# Organelle-specific photoactivation and dual-isotope labeling strategy reveals phosphatidylethanolamine metabolic flux

**DOI:** 10.1101/2022.11.03.514994

**Authors:** Clémence Simon, Antonino Asaro, Suihan Feng, Howard Riezman

## Abstract

Phosphatidylethanolamine metabolism plays essential roles in eukaryotic cells but has not been completely resolved due to its complexity. This is because lipid species, unlike proteins or nucleic acids, cannot be easily manipulated at the single molecule level or controlled with subcellular resolution, two of the key factors toward understanding their functions. Here, we use the organelle-targeting photoactivation method to study PE metabolism in living cells with a high spatiotemporal resolution. Containing predefined PE structures, we designed probes which can be selectively introduced to the ER or mitochondria to compare their metabolic products according to their subcellular localization. We combined photo-uncaging method with dual stable isotopic labeling to track PE metabolism in living cells by mass spectrometry analysis. Our results reveal that both mitochondrial- and ER-released PE participate in phospholipid remodeling, and that PE methylation can be detectable only under particular conditions. Thus, our method provides a framework to study phospholipid metabolism at subcellular resolution.

## Introduction

Despite their essential functions in health and disease, lipid metabolism is still less studied than metabolism of other biomolecules. Unlike proteins or nucleic acid structures, the highly diverse lipid species are not directly genetically encoded and cannot be selectively overexpressed/reduced. Moreover, delivery of exogenous lipids to living cells is a great challenge in lipid biology as many of them are highly hydrophobic, resulting in low solubility in aqueous solutions. In addition, some of them also bear charges on their headgroup, preventing their crossing of the plasma membrane. Thus, it cannot be predicted if and where they will be transported into cells. Lastly, lipid metabolic processes are very fast and occur in precise subcellular locations, making them difficult to detect. Over the past twenty years, chemical tools have been developed and combined with bioanalytical techniques in order to address lipid chemical complexity (for example chemical reporters, photocrosslinkable, photoswitchable or photocaged lipids).^1,2^ The photocaging method allows to deliver native species on demand with light and is therefore ideal to study lipid biological processes with high temporal resolution. It consists of adding a photolabile protecting group on the lipid headgroup to inhibit its activity. The caged lipid can be incorporated and accumulated in living cells in which it won’t be recognized by enzymes. Then, a rapid UV illumination removes the chemical protecting group, and delivers the bioactive lipid in a non-invasive manner. Photocaging has been applied to a number of lipid messengers like phosphoinositides,^3^ diacylglycerides,^4^ ceramides^5^ and fatty acids.^6,7^ While these studies focused on lipid roles in signalling processes, examples of photocaging tools to trace their metabolism are rare. In the past years, our group has developed a series of photocaged lipids targeted to different subcellular compartments including lysosomes and mitochondria. We combined this approach with stable-isotope labeling to trace sphingosine metabolism, and found that its metabolic conversion to ceramide versus sphingosine-1-phosphate depends on its original location to subcellular compartments, showing a metabolic bias based on subcellular location.^8,9^ Here we extended this approach of tracking local lipid metabolic flux to phosphatidylethanolamine (PE).

PE represents 25% of all phospholipids in mammals and corresponds to the second most abundant glycerophospholipid (GPL) behind phosphatidylcholine (PC).^10,11^ Enriched in the mitochondrial inner membrane, PE has a myriad of biological roles in cells.^12–16^ Given its biological importance, abnormal PE metabolism can lead to severe diseases like cancers or diabetes.^17,18^ PE metabolism holds a unique place in lipid mammalian cell biology and has not been completely resolved due to its complexity.^10,11^ First, it can be synthesized by 2 major pathways localized either in the ER by the CDP-ethanolamine pathway starting with ethanolamine or in mitochondrial inner membranes by phosphatidylserine (PS) decarboxylation mediated by phosphatidylserine decarboxylase (PSD). Then, PE metabolism involves its conversion into other GPL species and its remodeling into other PE species with different acyl chain compositions (figure S1). Studies of PE metabolism in mammals, based for a long time on radioactive precursors^19–22^ and more recently on stable isotope-labeled precursors^23–25^ suggest that its metabolism, transport and functions depend on its localization between ER and mitochondria. However, little is known in terms of quantitative and qualitative contribution of each pathway.

Here, we use an organelle-targeting photoactivation method to study PE metabolism in living cells with a high spatiotemporal resolution. We designed different PE probes which can be selectively introduced to the ER versus mitochondria to compare their metabolic products. We combined the photo-uncaging method with dual-stable isotopic labeling, which enabled us to track PE transformations in living cells with high resolution mass spectrometry (figure 1).

**Figure 1.**
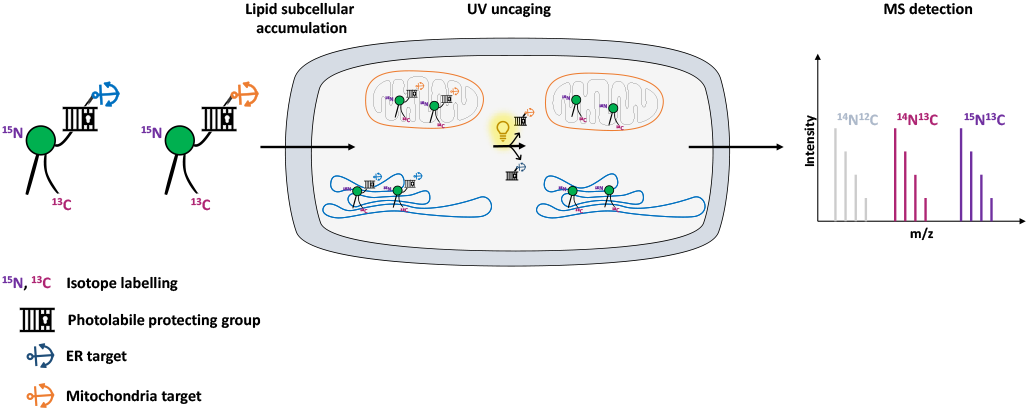
Schematic illustration of organelle-specific uncaging and MS-based metabolic tracking of PE probes. Isotope-labeled PE is equipped with a photocaging group functionalized with targeting groups that localize it to specific compartments (mitochondria or ER) in living cells, where the phospholipid can subsequently be photoreleased.

**Scheme S1.**
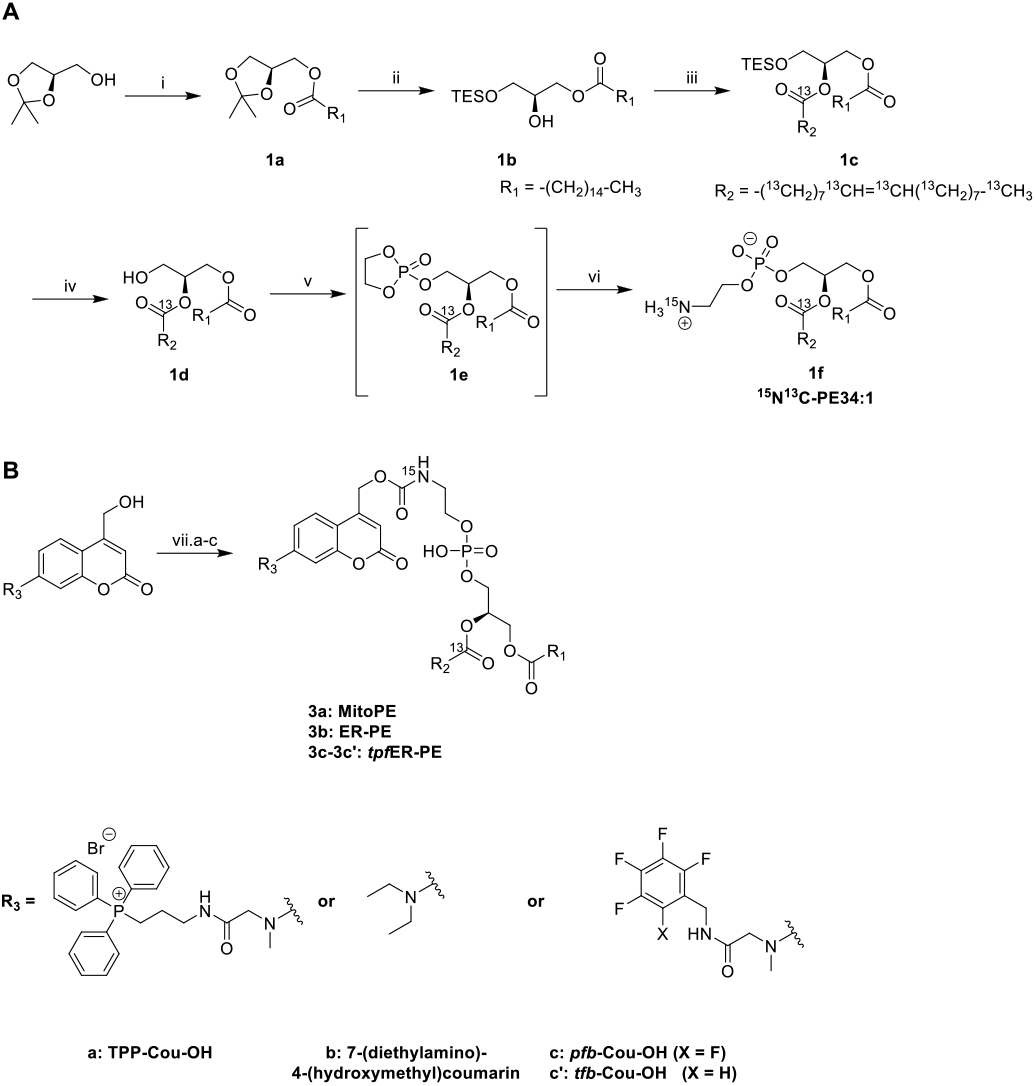
A) Synthesis of ^15^N^13^C-PE34:1. Conditions: (i) palmitic acid, DCC, DMAP, DCM, rt, 18 h - 85%; (ii) 1) TESOTf, DIEA, DCE, 95°C, 36 h; 2) I_2_, Na_2_CO_3_, H_2_O, THF, rt, 4 h - 54% over 2 steps; (iii) oleic acid-^13^C_18_, EDC, DMAP, DCM, rt, 12 h - 93%; (iv) FeCl_3_, MeOH, DCM, rt, 1 h - 92%; (v) 2-chloro-2-oxo-l,3,2-dioxaphospholane, TEA, ACN, 0°C to rt, 12 h; (vi) ^15^NH_3_, ACN, 0°C to 65°C, 24 h - 6% over 2 steps. B) Synthesis of MitoPE, ER-PE and *tpfER*-PE (compounds 3a, 3b and 3c respectively). Condition (vii.a) for MitoPE: 1) bis-(4-nitrophenyl)carbonate, DIEA, DCM, rt, 3 h; 2) ^15^N^13^C-PE34:1, DCM, 38°C, 6 h - 41%. Condition (vii.b) for ER-PE: (i) 1) triphosgene, DIEA, THF, 0°C, 4h; 2) ^15^N^13^C-PE34:1, DIEA, DCM, 0°C to rt, 12 h - 22%. Condition (vii.c) for *tpfER*-PE: 1) bis-(4-nitrophenyl)carbonate, DIEA, DCM, rt, 12 h; 2) ^15^N^13^C-PE34:1, DCM, 39°C, 24 h -43%.

## Results

The first step of the strategy was to design photocaged PE probes. Concerning its composition, the synthesized PE bears a palmitoyl and an oleoyl chain on sn1 and sn2 position respectively, giving PE34:1 (or PE16:0-18:1), one of the most abundant PE species in mammals. Most importantly, we labeled PE with two different heavy isotopes to ensure the tracing of different metabolic events using mass spectrometry: eighteen ^13^C labeled carbons on the oleoyl acyl chain and one ^15^N on the amine headgroup, giving ^15^N^13^C-PE34:1 (or ^15^N-PE16:0-^13^C18:1). PE synthesis is described in scheme S1A and starts with the (*R*) enantiomer of protected glycerol. The first key step of the synthetic route is the esterification of the hydroxyl at sn2 position of compound **1b** with commercial oleic acid-^13^C_18_. The second key step is the nucleophilic addition of ^15^N-ammonia on cyclic phosphotriester intermediates (**1e**) to get the final ^15^N^13^C-PE34:1 (compound **1f**). Measured optical rotation of the isotope labeled PE was similar to its non-labeled naturally occurring commercial analog (*R*)-POPE (+4.5 and +5.1 respectively), confirming its high enantiospecificity.

To capture the rapid turnover of PE, we synthesized different caged PE probes from ^15^N^13^C-PE34:1 to deliver it to specific subcellular compartments (scheme S1B). Bioactive small molecules are usually caged with nitrophenyl or coumarin derivatives to inhibit their activity. Coumarin was used in our case because of its intrinsic fluorescence properties visible by optical microscopy and was added on the PE headgroup. First, we synthesized the triphenyphosphonium-labeled-coumarin-alcohol (TPP-Cou-OH) using a previously described protocol to deliver PE specifically into mitochondria.^8^ Compound **3a** was synthesized by coupling TPP-Cou-OH with ^15^N^13^C-PE34:1 thought a carbamate linkage. In the same way, we combined the commercial 7-(diethylamino)-4-(hydroxymethyl)coumarin bearing no organelle tag to PE in order to see if this global-caged lipid (compound **3b**) could be delivered in a more diffuse way in cells. Aiming to develop an ER-specific probe, we also coupled ^15^N^13^C-PE34:1 with a coumarin functionalized with a mixture of pentafluorobenzyl and tetrafluorobenzyl (*pfb*-and *tfb*-Cou-OH) to accumulate PE specifically in the ER (compounds **3c-3c’**).^26^ We first tested the photo-cleavage efficiency of the synthesized probes. Upon UV illumination, compounds **3a, 3b** and **3c-3c’** were quickly hydrolysed in chloroform/methanol (1:1) to give free ^15^N^13^C-PE34:1 within 2 min. (figure S2). Next, we sought to determine the localization of each compound in living cells. In HeLa cells, we incubated compound **3a** (25 μM), which was complexed with liposomes (POPC/cholesterol 1:1) and methyl-β-cyclodextrin (mβCD), and compared its fluorescent signals with the ones from MitoTracker. The fluorescence signal of compound **3a** showed a good colocalization with a MitoTracker, suggesting it is mainly introduced into mitochondria in living cells (figure 2A-B), and leading us to rename it “MitoPE” to highlight its subcellular localization. Importantly, optimization experiments showed that MitoPE needed to be complexed first with a correct ratio of mβCD/liposomes to be efficiently introduced into HeLa cells (figure S3D). Indeed, previous studies showed that mβCD is a good donor to transfer long-chain fluorescent or natural phospholipids from vesicles (composed of POPC/cholesterol) to cultured cells.^27,28^ In the absence of mβCD or non-optimal donor/vesicles ratio, there is no specific staining inside cells, but blue dots representing insoluble aggregates inside and outside cells (figure S3A-C).

**Figure 2.**
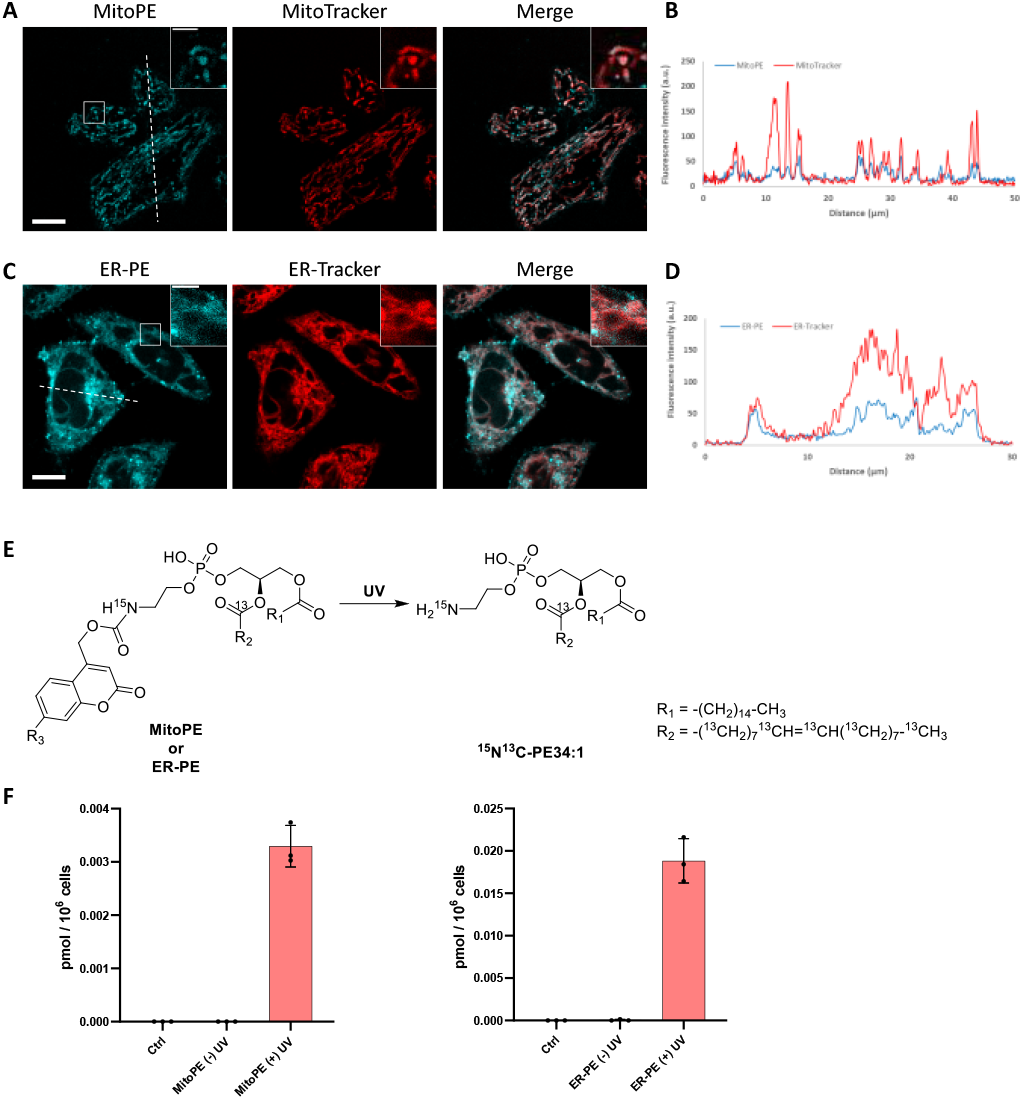
Localization and uncaging efficiency of organelle-targeted caged PE probes in living HeLa cells. A, C) Representative fluorescence image of HeLa cells stained with A) MitoPE (25 μM, previously complexed with cholesterol/POPC 1:1 and mβCD, see detailed description in the ESI) and MitoTracker (50 nM) or C) ER-PE (25 μM) and ER-Tracker (1 μM). Scale bar: 10 μm and 3 μm (insert). B, D) Intensity profiles of the white dotted lines in B) MitoPE and Mitotracker channels, or D) ER-PE and ER-Tracker channels. E) Illustration of photoactivation of PE probes. F) Uncaging efficiency of MitoPE and ER-PE in living cells. Fully confluent cells in 35 mm culture dishes were incubated with the caged probes i) MitoPE 5 μM, previously complexed with cholesterol/POPC 1:1 and mβCD or ii) ER-PE 5 pμM for 15 min, washed, irradiated under 350–450 nm UV light for 2 min on ice. Cells were collected immediately after UV irradiation. Lipids were extracted and measured by LC-MS. Values were normalized with respect to the amount of internal standards and cell numbers. Data represents the average of three independent experiments. Error bars represent SD.

Concerning compound **3b**, no vesicle carrier was necessary to introduce it into living HeLa cells. The absence of the three hydrophobic benzenes forming the TPP cation could explain its better solubility compared to MitoPE, even if the latter has a supplementary positive charge. Hence, HeLa cells were incubated with the global-caged PE (25 μM) for 15 min before washing. Despite the presence of some nonspecific dots, its fluorescence staining mostly overlaps with an ER-Tracker, meaning that it is mainly localized to the ER (figure 2C-D). We therefore renamed the compound **3b** to “ER-PE” to highlight its subcellular localization. Similarly, compounds **3c-3c’** complexed with liposomes/mβCD accumulates mainly in the ER (figure S4A-B), so we renamed it by “*tpf*ER-PE”. However, *tpf*ER-PE seemed to present more non-specific puncta than ER-PE, which can be explained by the low solubility of the fluorinated groups. So, we used the latter during our study.

Next, we sought to determine the efficiency of ^15^N^13^C-PE34:1 released in living cells, the isotope labeling allowing to distinguish it from its endogenous analogue by mass spectrometry (figure 2E). HeLa cells were incubated with the probe of interest and exposed to UV. Directly after the uncaging, cells were collected for lipid extraction and analysis by LC-MS. For both MitoPE and ER-PE, data showed a robust appearance of the ^15^N^13^C-PE34:1 signal after UV irradiation compared to the cell treated with caged-lipid without UV exposure and control cells treated with no caged-molecule (figure 2F). The signal specificity of ^15^N^13^C-PE34:1 in cells and of all following labeled species was further confirmed by LC-MS/MS analysis (see LC/MS-MS spectrum in the supporting information). Therefore, the uncaging was efficient for both probes. Interestingly, the level of released ^15^N^13^C-PE34:1 after UV uncaging is higher (approximatively 6 times) in cells incubated with ER-PE than MitoPE. This could be explained by the difference in size of the organelles where the probe is localized (mitochondria *vs* ER) and perhaps by differences in solubility between the two probes. Even though we treated the cells with equal amount of probes, only MitoPE needs to be complexed beforehand with a donor carrier, meaning that it will not enter in the HeLa cells as easily as ER-PE. Taken together, these results show that MitoPE and ER-PE can selectively be introduced into mitochondria and the ER of HeLa cells respectively, and they can efficiently release 15N13C-PE34:1 under UV illumination. To our knowledge, this is the first time that a PE can be delivered with a high spatiotemporal resolution in living cells.

Immediately after photo-uncaging, 15N13C-PE34:1 can be released at specific places in living HeLa cells, recognized by enzymes and recycled, degraded or converted into different metabolites. These potential metabolites can be traced by high resolution mass spectrometry due to the 15N and 13C isotopic labeling on amine headgroup and oleoyl sn2 acyl chain respectively, allowing us to study phospholipid remodeling. Also called the Lands’ cycle, this pathway consists of deacylation/reacylation steps of newly synthesized phospholipids performed by phospholipases and acyltransferases respectively to modulate their composition and maintain membrane diversity in mammalian cells.^29^ To monitor phospholipid remodeling, we treated cells either with MitoPE, or with ER-PE. After the uncaging steps cells were incubated at 37°C during different times (0, 20 or 90 min) and then collected for lipid extraction and LC-MS analysis. To be able to compare the data coming from MitoPE and ER-PE photolysis, we normalized the signals of interest by the quantity of released ^15^N^13^C-PE. PE34:1 originating in the two locations was involved in sn2 remodeling, but this pathway was more prevalent for the ER localized lipid (figure 3A-B). For the ER localized lipid, a signal corresponding to ^15^N-LysoPE16:0-0:0 is visible from 0 min to 90 min with a slight increase at time 20, meaning that phospholipase A2 enzymes are able to cleave the ^13^C_18_-oleic acid at sn2 position and generate ^15^N-LysoPE 16:0-0:0. Then ^13^C_18_-oleic acid must be converted to oleoyl-CoA and used by lysophospholipid acyltransferases (LPLAT) to acylate the sn2 position of endogenous phospholipids like PC, giving ^13^C-labeled PC species. The composition of detected ^13^C-PC species follows the order of the more abundant species with an oleoyl chain at the sn2 position in HeLa cells. Five species are detected after ER-PE uncaging (^13^C-PC34:1; 36:2; 36:1; 34:1-O and 34:2) and seem to continue to increase up to 90 min. Three species are visible after MitoPE photolysis (^13^C-PC34:1, 36:2 and 34:2) and they seem to reach a plateau already at 20 min. Some species are already visible at 0 min, meaning that the remodeling can take place already during the photoactivation period which last 2 min.

**Figure 3.**
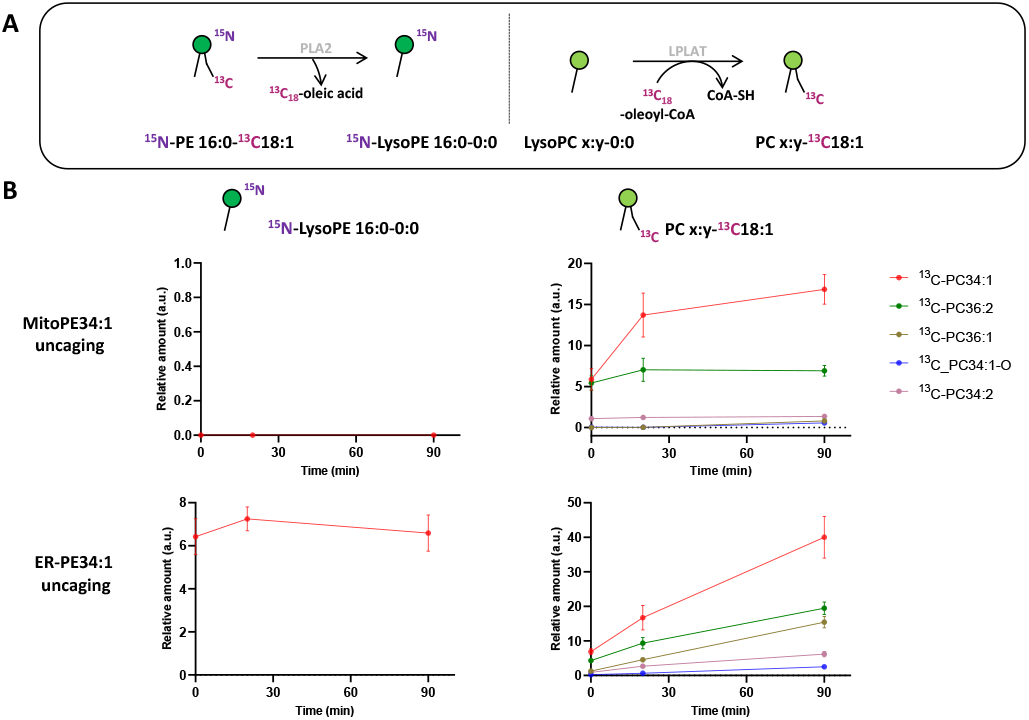
Released ^15^N^13^C-PE34:1 participates in sn2 remodeling of endogenous lysoPC species in living cells. MitoPE or ER-PE were introduced in HeLa cells as outlined in “experimental procedures” for 15 min. Then cells were washed, irradiated by UV light for 2 min on ice, incubated at 37°C, and collected for lipid extraction at different time points (0, 20 or 90 min). A) The ^13^C_18_-oleoyl acyl chain of released ^15^N^13^C-PE34:1 is cleaved by phospholipase A2 (PLA2) and used by lysophospholipid acyltransferase (LPLAT) to acetylate the sn2 position of endogenous lysoPC species. B) LC-MS analysis of labeled species involved in sn2 remodeling of endogenous lysoPC species. Values were normalized with respect to the amount of internal standards and photo-released ^15^N^13^C-PE34:1. Data represent the average of three independent experiments. Error bars represent SD.

Longer incubation time of ^15^N^13^C-PE34:1 after probe photo-release leads to sn2 reacylation of other GPL classes with ^13^C-oleic acid. After 4 h and 16 h of ER-PE uncaging, we observed significant appearance of the more abundant PE species (^13^C-PE36:2; 34:1 and 36:1 after ER-PE uncaging; ^13^C-PE34:1 after MitoPE uncaging), PI species (^13^C-PI36:2; 36:1; 34:1; and 34:2 after ER-PE uncaging only) and of the more abundant PS species as well (^13^C-PS36:1; 36:2 and 34:1 after ER-PE uncaging only)(figure S5). Although ^13^C-PS 36:1 and 36:2 are necessarily generated by reacylation of their respective lysoPS forms, we cannot exclude the possibility that ^13^C-PS34: could also come from base exchange reaction between ^15^N-ethanolamine of ^15^N^13^C-PE34:1 and endogenous serine. The solution to verify this hypothesis would be to synthesize a new PE probe with a labeling of the glycerol skeleton in addition to the oleoyl acyl chain. Interestingly, while we can detect sn2 remodeling of endogenous lysoPE species with the cleaved ^13^C-oleic acid, reacylation of ^15^N-LysoPE16:0 to give ^15^N-PE PE16:0-18:1 or 15N-PE16:0-18:2 as abundant PE species is not detectable (data not shown).

Tracing metabolism of released 15N13C-PE34:1 after long incubation times allows to follow its sn1 remodeling. After the uncaging of ER-PE or MitoPE, cells were incubated at 37°C during 4 h or 16 h. First, for ER-PE photo-release, we can observe a signal specific of 15N-LysoPE0:0-13C18:1 at times 4 h and 16 h (figure 4B). We can also detect specific signals for 15N-PE18:0-13C18:1 and 15N-PE18:1-13C18:1. Together, these data suggest that phospholipase A1 has cleaved the palmitoyl acyl chain at sn1 position and an LPLAT is able to reacetylate the 15N-LysoPE0:0-13C18:1 with stearic acid or oleic acid (figure 4A). In the case of MitoPE, a signal of 15N-LysoPE0:0-13C18:1 is not visible, most likely because it is below the detection limit.

**Figure 4.**
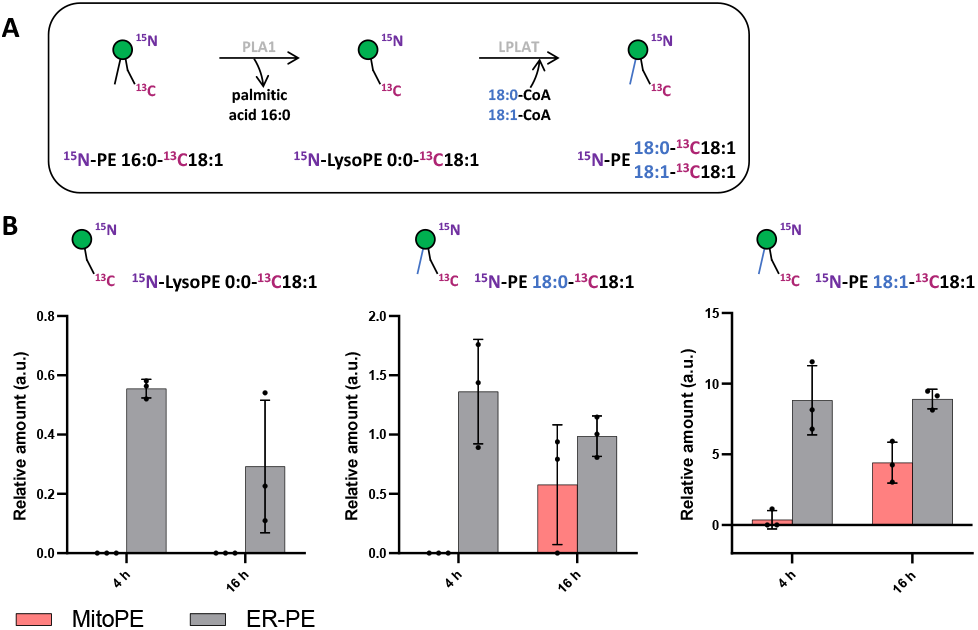
Sn1 remodeling of photo-released ^15^N^13^C-PE34:1 in living cells. MitoPE or ER-PE were introduced in HeLa cells as outlined in “experimental procedures” for 15 min. Then cells were washed, irradiated by UV light for 2 min on ice, incubated at 37°C, and collected for lipid extraction and LC-MS analysis at 4 or 16 h. A) Illustration of sn1 remodeling of released ^15^N^13^C-PE34:1 by phospholipase A1 (PLA1) and lysophospholipid acyltransferase (LPLAT). B) Analysis of labeled species involved in ^15^N^13^C-PE sn1 remodeling after its release from different sources as indicated. Values were normalized with respect to the amount of internal standards and photo-released ^15^N^13^C-PE34:1. Data represent the average of three independent experiments. Error bars represent SD.

However, a signal of ^15^N-PE18:0-^13^C18:1 starts to appear and a signal of ^15^N-PE18:1-^13^C18:1 is significant at 16 h. As phospholipid reacylation by LPLAT usually occur in the ER, this implies that ^15^N^13^C-PE34:1 released in mitochondria from MitoPE can be transported to the ER in HeLa cells. Together, these results show that ^15^N^13^C-PE34:1 is remodeled at the sn1 position and participates as well in the sn2 remodeling of endogenous lysoPL species.

As expected, ^15^N^13^C-PE released from *tpf*ER-PE after UV illumination is also involved in sn2 remodeling of endogenous PC species and undergoes its own sn1 remodeling (figure S4C-D).

Another important aspect of PE metabolism is its transformation into PC by the phosphatidylethanolamine-*N*-methyl transferase (PEMT, figure S1).^30^ PEMT is an integral membrane protein enriched in the ER and mitochondria-associated membranes (MAM) contact sites which converts PE into PC through 3 consecutive transmethylation reactions from S*-*adenosyl-methionine (SAM). This process is one of the three mammalian PC biosynthesis pathways and is essential in the liver where it accounts for 30% of total PC production. Contrary to yeast in which PEMT is a major pathway, its activity is less clear in mammals and seems to be organ-dependent. Moreover, study of PE methylation is complex because of the lack of suitable mammalian cell line that significantly expresses PEMT. Indeed, the photo-release of ^15^N^13^C-PE34:1 from the different sources (mitochondria or ER) during different incubation times (from 0 min until 16h) in HeLa cells did not lead to detection of any ^15^N^13^C-PC34:1 (*i*.*e*. ^15^N-PC16:0-^13^C18:1), signal which would come from PEMT methylation. This nevertheless demonstrates the importance to label PE at two different places of the chemical structure (here the oleoyl acyl chain *versus* the headgroup) to discriminate ^13^C-PC34:1 involving sn2 remodeling of endogenous lysoPC species from ^15^N^13^C-PC34:1 which would come from PEMT methylation. Therefore, we decided to overexpress PEMT in HeLa cells by transfecting a plasmid containing the isoform 2 of PEMT under the control of a constitutive promoter. Immunofluorescence of PEMT and western blot analysis (figure 5A-B) confirmed a high rate of transfection. According to the immunofluorescence, overexpressed PEMT is localized into ER and Golgi. To test for overexpression of PEMT activity, we incubated control cells and PEMT overexpressing cells with d_4_-ethanolamine (d_4_-Etn) for 1.5 h or 5 h before cell collection and lipid extraction. The obtained lipidomic data showed the incorporation of d_4_-Etn into d_4_-PE species, and a significant increase of d_4_-labeled PC species in overexpressed cells compared to control cells, confirming the increase of PEMT activity (figures S6 and 5C). After 5 h of incubation, d_4_-PC34:1 signal increased approximatively 114-fold in PEMT overexpressing cells compared to mock-transfected cells, corresponding approximatively to 20 % of d_4_-PE34:1 signal (figure 5C).

**Figure 5.**
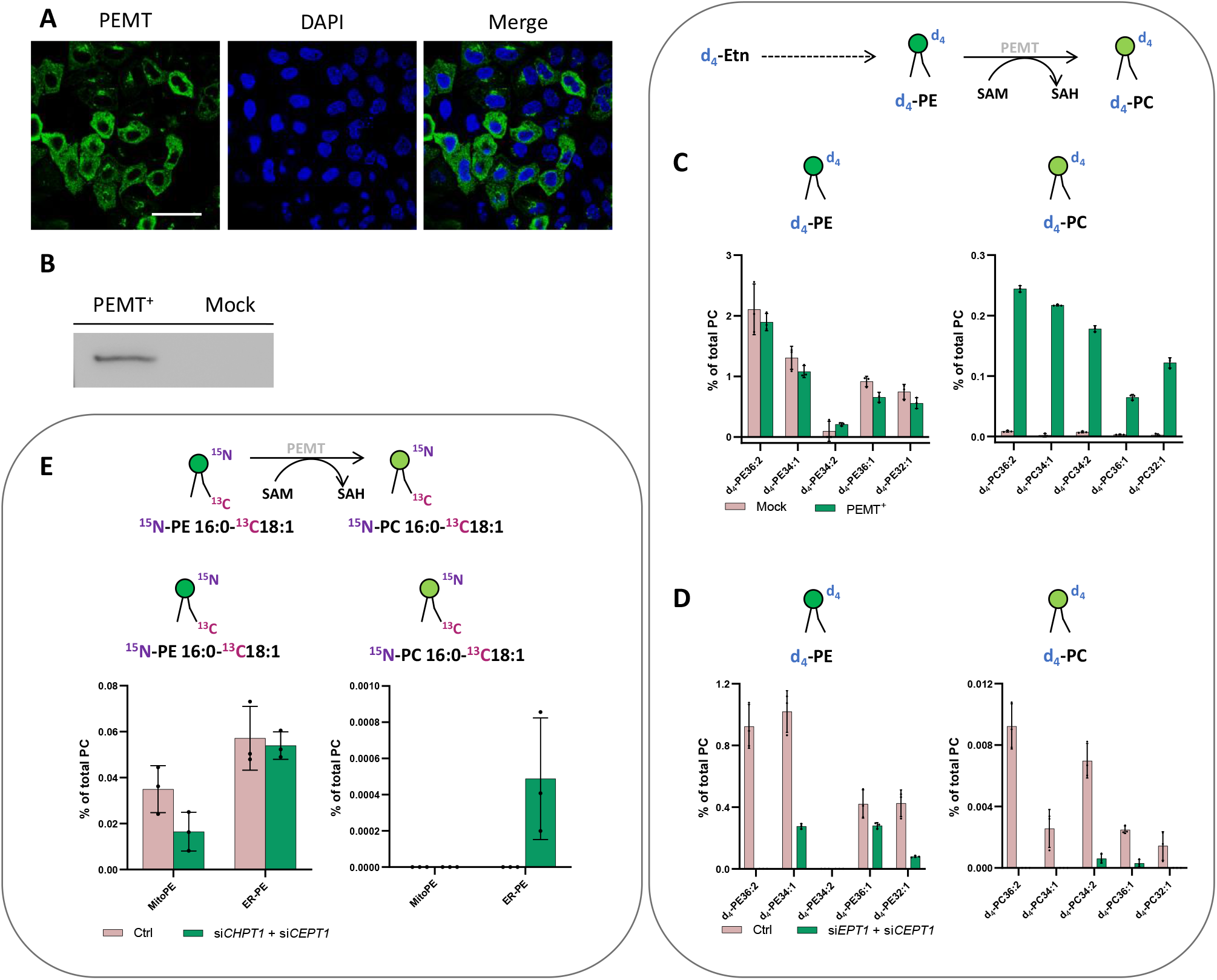
Analysis of PE methylation. A-B) PEMT gene overexpression in HeLa cells. A) HeLa cells were grown on glass dishes and transfected with PEMT-myc expression construct. 24 h after transfection, cells were fixed and stained with anti-Myc antibody (green). Cell nuclei were counterstained with DAPI (blue). Scale bar: 50 μm. B) Western blot analysis of PEMT-myc levels in cell extracts from transiently transfected cells. C) Incorporation of d_4_-Etn into lipid metabolites of HeLa cells transfected with PEMT2. HeLa control cells and transiently transfected with PEMT2 were washed once with PSB and incubated with d_4_-Etn-HCI (0.5 mM) for 5 h at 37°C. Then cells were washed three times with PBS and collected for lipid extraction. Top: illustration of d_4_-PE methylation to d_4_-PC. Bottom: LC-MS analysis of labeled species involved in PEMT pathway. D) Incorporation of d_4_-Etn into lipid metabolites of HeLa cells presenting PEMT overexpression and CDP-Ethanolamine pathway downregulation. PEMT overexpressing cells treated with negative siRNA (Ctrl) or siRNA for EPT1 (*siFPTl*) and CEPT1 (*siCEPTl*) were washed once with PSB and incubated with d_4_-Etn-HCI (0.5 mM) for 90 min at 37°C. Then cells were washed three times with PBS and collected for lipid extraction. E) Formation of ^15^N^13^C-PC34:1 in HeLa cells presenting PEMT overexpression and CDP-Choline pathway downregulation after photoactivation. PEMT overexpressing cells treated with negative siRNA (Ctrl) or siRNA for CPT1 (*siCHPTl*) and CEPT1 (*siCEPTl*) were washed once with PSB and incubated with MitoPE or ER-PE as outlined in “experimental procedures” for 15 min. Then cells were washed, irradiated by UV light for 2 min on ice, incubated for 4 h at 37°C and collected for lipid extraction. Top: illustration of ^15^N^13^C-PE34:1 methylation to ^15^N^13^C-PC34:1 by PEMT. Bottom: LC-MS analysis of labeled species involved in PEMT pathway. For each LC-MS analysis, values were normalized with respect to the amount of internal standards and total PC species. Data represent the average of three independent experiments. Error bars represent SD.

In some organisms including plants, PEMT enzyme can methylate the ethanolamine headgroup not only when part of the phospholipid but also directly after its activation to phosphoethanolamine in the CDP-ethanolamine pathway.^31^ To prove that increased PEMT activity is working on PE and not one of its precursors in HeLa cells, the latter were transfected with two siRNA targeting *EPT1* and *CEPT1*, afterwards PEMT was overexpressed and cells were incubated with d_4_-Etn for 1.5 h. *EPT1* and *CEPT1* encode respectively phosphotransferase EPT1 and CEPT1 that act at the last step of the CDP-ethanolamine pathway to generate PE from CDP-choline. Their effective silencing in transfected cells was confirmed by RT-qPCR (figure S7A). Then, the lipidomic data confirmed a decrease of PE synthesis in cells presenting PEMT overexpression combined with CDP-ethanolamine pathway downregulation compared with control cells with no siRNA silencing (figure 5D). Signal of d_4_-PC species significantly decreased as well, meaning that in HeLa cells, PEMT enzyme seems mainly methylate PE phospholipids after their complete biosynthesis and cannot act upstream at the start of the CDP-ethanolamine pathway.

Despite PEMT overexpression, the use of our probes to photo-release ^15^N^13^C-PE34:1 did not lead to any detectable ^15^N^13^C-PC34:1 signal (figure S8). The major pathway for PC biosynthesis in mammalian cells is the CDP-choline pathway by analogy to the CDP-ethanolamine pathway for PE biosynthesis. We therefore decided to inhibit the CDP-choline pathway in addition to PEMT overexpression to further promote PE methylation. The final step of the CDP-choline pathway is mediated by two choline phosphotransferases (CPT1 and CEPT1 respectively encoded by *CHPT1* and *CEPT1*) able to transfer phosphocholine from CDP-choline into a diacylglycerol backbone to form PC. Hence, HeLa cells were transfected with two siRNA targeting *CHPT1* and *CEPT1*, afterwards they were transfected with PEMT2 plasmid. The effective silencing of targeted genes in transfected cells was confirmed by RT-qPCR (figure S7B). Then, after MitoPE or ER-PE incubation and uncaging, transfected cells were incubated at 37°C for 4 h (figure 5E). ^15^N^13^C-PC34:1 signal was finally detectable when ^15^N^13^C-PE34:1 was released from the ER in cells presenting PEMT overexpression combined with CDP-Choline pathway downregulation, meaning that ER-PE probe can be used to specifically study PE methylation. Signal of ^15^N^13^C-PC34:1 corresponds to only 0.9 % of ^15^N^13^C-PE34:1 released signal. No signal was detected under the same conditions with MitoPE. This difference between both probes can be explained by the localization of PEMT in the ER, directly accessible for ER-PE.

## Discussion

PE metabolism holds a unique place in lipid cell biology and exemplifies the complexity of lipid biosynthesis and transport mechanisms. Although some studies suggest that its metabolism and function depend on its biosynthetic origin,^17,22,23^ the experimental means to directly address this issue are limited. Here, we developed novel photo-activation techniques, which enables delivering PE into different subcellular compartments, and thereafter releasing PE molecules with a high spatiotemporal resolution. We combined it with dual isotopic labeling to quantitatively analyze and compare PE metabolic conversion in HeLa cells. Installation of the labeling on two strategic positions of the lipid molecule is essential to discriminate the remodeling pathway from the PEMT pathway.

Phospholipid remodeling was mainly studied in the past by using radioactive labeled or stable isotopically labeled precursors (choline, ethanolamine or glycerol). However, due to the lack of specificity, it needs details on the remodeling pathways and the kinetics. Only a few papers describe instead the used of isotopically labeled phospholipids in mammalian cells or in yeast.^32,33^ In mammalian cells, the authors showed that atypical phospholipids were rapidly remodeled to endogenous abundant species whereas the introduced species identical to endogenous ones were remodeled only to a limited degree. Our probes allow to study phospholipid remodeling with a high spatiotemporal resolution due to the presence of functionalized photolabile protecting groups on the PE. The use of two different stable isotopes at two strategic positions of the lipid allows to differentiate the sn1 remodeling of the labeled lipid itself from sn2 remodeling of endogenous phospholipids. While sn2 remodeling has been well described,^29,34^ it is not clear how the composition of saturated fatty acid is controlled at sn1 position. It has been recently shown that lysophosphatidylglycerol acyltransferase 1 (LPGAT1) is a key remodeling enzyme located in the ER and involved in the regulation of stearate/palmitate ratio at sn1 position of LysoPE species.^35^ It is possible that LPGAT1 is responsible of the ^15^N-PE16:0-^13^C18:1 (*i*.*e*. ^15^N^13^C-PE34:1) remodeling into ^15^N-PE18:0-^13^C18:1. This could be investigated by the use of LPGAT1 knock out cell lines.

The PEMT pathway represents the second biosynthetic pathway of PC via PE methylation in the ER. However, PEMT activity has not been clearly implicated in a number of tissues because this pathway is mainly liver specific, and cell culture models usually express only traces of PEMT activity. To our knowledge, PE methylation into PC in mammalian models was demonstrated with GPL precursors and never with complete GPLs.^36,37^ Some previous studies also postulate that only the newly made PE could be methylated.^38^ As Hela cells only present a minor PEMT activity, we overexpressed the gene and confirmed the increase of d_4_-PE methylation into d_4_-PC using d_4_-Etn as PE precursors. However, overexpression alone was not sufficient to detect any signal of ^15^N^13^C-PC34:1 coming from ^15^N^13^C-PE34:1 methylation, suggesting that this pathway is very inefficient on consuming PE. We then combined PEMT overexpression with disruption of CDP-Choline pathway to further promote methylation of the PE probe. This condition allowed detection of the formation of ^15^N^13^C-PC34:1 when PE was photo-released from the ER, meaning that ER-PE probe can be used to study PEMT activity. We also photo-released PE from the ER-PE probe in other cell lines including mouse primary hepatocytes, in which PEMT initial level should be very high. However, we did not detect any signal of ^15^N^13^C-PC coming from PEMT activity (figure S9). The striking inefficiency of this pathway suggests that only a very small portion of the PE in cells is substrate for PEMT. When compared to our measurements of the efficiency of PE to PC conversion by labeling with d_4_-Etn, we must conclude that there are limitations on the metabolism of PE that we do not understand. Two possible explanations could be entertained as limitations for accessibility to PEMT. First, it could be that only newly synthesized PE can be converted to PC by PEMT. Second, it could be that only not all PE can access the location where PEMT is localized, suggesting that PE cannot freely diffuse anywhere in the ER. This is consistent with our previous data on sphingosine transport.^9^ Sphingosine coming from lysosomes can access ceramide synthases, but not sphingosine kinases, both of which are found in the ER. Our data shows that PE metabolism is far more complex than originally thought, which could only be seen using tools like we have prepared.

The diversity of PE metabolism implies its transport between the ER and mitochondria, probably involving the MAM and some lipid transfer proteins.^39^ The lipid probes we developed presented a chemical tool that can be combined with genetic tools to investigate PE transport and its impact on PE metabolism.

## Conclusions

To summarize, we developed organelle-targeted photoactivated probes to study PE metabolism in living cells with a high spatiotemporal resolution. We designed different PE probes which can be selectively introduced in the ER *versus* mitochondria to compare their metabolism according to their subcellular localization. LC-MS results revealed that both mitochondria-and ER-photo-released PE are involved in sn1 and sn2 phospholipid remodeling. It also showed that PEMT methylation was very inefficient but could be detected with ER-PE under specific conditions, suggesting unknown limitations on the PE methylation pathway. Our data thus provides a framework to study phospholipid metabolism at subcellular resolution.

## Data availability

Experimental data associated with this work have been provided in the ESI.

## Author Contributions

C.S. and H.R. designed all the experiments; C.S. and A.A. performed research; C.S. analyzed data; C.S. wrote the manuscript; C.S., A.A., S.F. and H.R. corrected the manuscript; C.S., S.F. and H.R. conceived and supervised the project.

## Acknowledgements

We thank Takeshi Haramaya for helpful discussion, Isabelle Riezman and Emmanuel Varesio for technical support in lipidomic experiments. We thank the Bioimaging Center (University of Geneva) for technical assistance. This work was supported by the Human Frontier Science Program (LT000552/2020-L), the Leducq Foundation, the Swiss National Science Foundation (310030_184949), the NCCR Chemical Biology (51NF40-185898) and the Deutsche Forschungsgemeinschaft. S.F. was supported by Shanghai Municipal Science and Technology Major Project (2019SHZDZX02) and Shanghai Pujiang Program (22PJ1414500).

