## Supplementary information for "Organelle-specific photoactivation and dual-isotope labeling strategy reveals phosphatidylethanolamine metabolic flux"

#### General material and methods

(*R*)-1-palmitoyl-2-oleoyl-sn-glycero-3-phosphoethanolamine (PE16:0-18:1) was purchased from Avanti. Oleic acid-$^{13}C_{18}$ was purchased from Sigma Aldrich. Ammonia gas $^{15}N$ (98% $^{15}N$) was purchased from Eurisotop. Other chemical reagents were purchased from Sigma Aldrich, TCI and Fluorochem and were used with no further purification.

Reactions were monitored by TLC, using pre-coated 0.20 mm ALUGRAM XtraSil G UV254 sheets from Macherey-Nagel. Detection was done under UV light at 254 nm or 365 nm and/or by chemical development of the plates with solutions of phosphomolybdic acid or potassium permanganate.

Purifications were performed by manual chromatography.

$^{1}H$ and $^{13}C$-NMR spectra were recorded on a Bruker AMX-400 MHz or 500 MHz spectrometer. $^{31}P$ and $^{19}F$-NMR spectra were recorded on a Bruker AMX-300 MHz. Chemical shifts are given in ppm (δ) using the NMR solvent as internal references and J values are reported in Hz. Splitting patterns are designated as follows: s, singlet; d, doublet; t, triplet; q, quartet; p, pentuplet; m, multiplet; br, broad. Spectra of final compounds are provided in the final section of the present document.

All mass spectra were measured by electrospray ionization (ESI) using either a Thermo Electron Corporation HPLC coupled with a Thermo Scientific Finnigan Surveyor MSQ Spectrometer System (LC-ESI-MS), or with a TSQ Vantage Triple Stage Quadrupole Mass Spectrometer (Thermo Scientific) by direct infusion (ESI-MS).

High-resolution mass spectra (HR-MS) data were recorded on a Xevo G2 Tof spectrometer equipped with orthogonal electrospray interface (ESI).

Optical rotations were measured as $CHCl_3$ solutions (c = 0.01 g/mL) on a JASCO P1030 polarimeter at 20°C using a Na lamp (589 nm) and converted to specific rotations $[\alpha]_D$. (*R*) configuration of $^{15}N^{13}C$-PE34:1 was confirmed by comparing its specific rotation with that of commercial (*R*)-1-palmitoyl-2-oleoyl-sn-glycero-3-phosphoethanolamine (non-isotopically labeled analogue).

### Synthesis procedures

1. Synthesis of $^{15}N^{13}C$-PE34:1

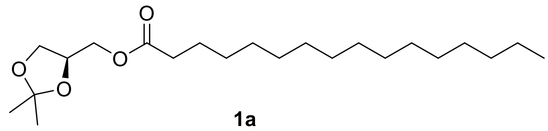

**1a**

**Compound 1a.** Into a solution of palmitic acid (3.8 g, 14.8 mmol), *N*,*N'*-dicyclohexylcarbodiimide (DCC, 3.7 g, 17.9 mmol) and 4-(dimethylamino)pyridine (DMAP, 0.5 g, 4.1 mmol) dissolved in DCM (20 mL) was added (*R*)-(−)-2,2-dimethyl-1,3-dioxolane-4-methanol (2.0 g, 15.1 mmol). After 18 h of stirring at rt, the mixture was evaporated under reduced pressure. The residue was purified by chromatography (cyclohexane/EtOAc, 9.5:0.5) to yield compound **1a** as a white solid (4.7 g, 19.4 mmol, 85%).

$^{1}$H NMR (400 MHz, $CDCl_3$) δ 4.30 (m, 1H), 4.16 (dd, *J* = 11.5, 4.7 Hz, 1H), 4.09 (dd, *J* = 11.5, 4.7 Hz, 1H), 4.06 (dd, *J* = 8.5, 6.5 Hz, 1H), 3.73 (dd, *J* = 8.5, 6.2 Hz, 1H), 2.33 (t, *J* = 7.6 Hz, 2H), 1.62 (p, *J* = 7.4 Hz, 2H), 1.43 (s, 3H), 1.36 (s, 3H), 1.26 (m, 24H), 0.87 (t, *J* = 7.0 Hz, 3H). $^{13}$C NMR (101 MHz, $CDCl_3$) δ 173.76, 109.94, 73.81, 66.50, 64.65, 34.25, 32.06, 29.83, 29.82, 29.81, 29.79, 29.78, 29.73, 29.59, 29.50, 29.38, 29.26, 26.82, 25.54, 25.03, 22.83, 14.25.

ESI-MS: m/z $[M + Na]^+$ calcd for $C_{22}H_{42}NaO_4$: 393.30; found: 393.15; m/z $[M + K]^+$ calcd for $C_{22}H_{42}KO_4$: 409.27; found: 409.04.

$[\alpha]_D^{20}$ = + 1.06 (c = 0.01 g/mL, $CHCl_3$).

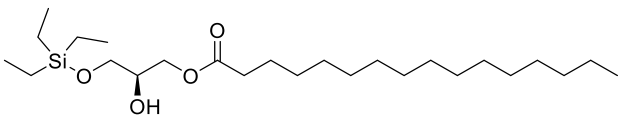

**1b**

**Compound 1b.** To a solution of compound **1a** (2.0 g, 5.4 mmol) dissolved in dichloroethane (20 mL) DIEA (2.8 mL, 16.2 mmol) and TESOTf (1.7 g, 6.5 mmol) were added. After 12 h of stirring at reflux an additional portion of TESOTf (0.7 g, 2.7 mmol) was added. Stirring at reflux was continued for additional 24 h and the reaction mixture allowed to reach rt. The reaction mixture was put into a mixture of EtOAc and $H_2O$ (1:1, 300 mL), the layers were separated, the organic layer washed with $H_2O$ (1 x 100 mL) and brine (1 x 100 mL), dried over anhydrous sodium sulfate, filtered and evaporated under reduced pressure. The residue was dissolved in a mixture of THF (44 mL) and $Na_2CO_3$ solution (10% in $H_2O$, 19 mL). The resulting suspension was treated with iodine (2.1 g, 8.3 mmol). After 4 h of stirring the reaction mixture was transferred onto a mixture of EtOAc and saturated $Na_2S_2O_3$ solution (1:1, 150 mL). The layers were separated, the organic layer was washed with $H_2O$ (1 x 100 mL) and brine (1 x 100 mL), dried over anhydrous sodium sulfate, filtered and evaporated under reduced pressure. The residue was purified by chromatography (cyclohexane/EtOAc gradient, 9.5:0.5 to 9:1) to yield compound **1b** as a light-yellow oil (1.3 g, 2.9 mmol, 54%).

$^{1}$H NMR (400 MHz, $CDCl_3$) δ 4.19 – 4.07 (m, 2H), 3.88 (p, *J* = 4.8 Hz, 1H), 3.67 (dd, *J* = 4.6, 10.1 Hz, 1H), 3.60 (dd, *J* = 5.8, 10.1 Hz, 1H), 2.33 (t, *J* = 7.6 Hz, 2H), 1.62 (p, *J* = 7.4 Hz, 2H), 1.25 (m, 25H), 0.96 (t, *J* = 7.9 Hz, 9H), 0.88 (t, *J* = 7.0 Hz, 3H), 0.62 (q, *J* = 8.1 Hz, 6H). $^{13}$C NMR (101 MHz, $CDCl_3$) δ 174.13, 70.19, 65.14, 63.52, 34.35, 32.07, 29.84 (2C), 29.82, 29.80 (2C), 29.76, 29.61, 29.51, 29.42, 29.29, 25.09, 22.84, 14.26, 6.81 (3C), 4.43 (3C). ESI-MS: m/z $[M + H]^+$ calcd for $C_{25}H_{53}O_4Si$: 445.37; found: 445.63; m/z $[M + Na]^+$ calcd for $C_{25}H_{52}NaO_4Si$: 467.35; found: 467.67.

$[\alpha]_D^{20}$ = + 1.00 (c = 0.01 g/mL, $CHCl_3$).

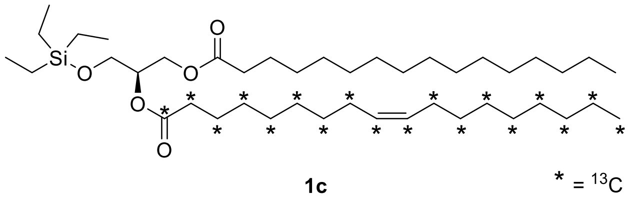

**1c**

**Compound 1c.** A solution of commercial oleic acid-$^{13}C_{18}$ (487 mg, 1.62 mmol), EDC (629 mg, 4.05 mmol) and DMAP (135 mg, 1.10 mmol) in dry DCM (15 mL) was stirred for 15 min at rt and subsequently treated with a solution of **1b** (600 mg, 1.35 mmol) in dry DCM (12 mL) and the solution was stirred at rt overnight. Upon completion, the DCM was removed under reduced pressure and the residue was dissolved in a mixture of EtOAc and $H_2O$ (1:1, 150 mL). The mixture was extracted once with EtOAc (75 mL). The organic layer was washed with $H_2O$ (75 mL) and brine (75 mL), dried over anhydrous sodium sulfate, filtered and evaporated under reduced pressure. The residue was purified by chromatography (cyclohexane/EtOAc gradient, 9.75:0.25 to 8:2) to yield compound **1c** as a colorless oil (913 mg, 1.26 mmol, 93%).

$^{1}$H decoupled $^{13}$C NMR (400 MHz, $CDCl_3$) δ 5.60 – 5.17 (m, 2H), 5.07 (p, *J* = 5.4 Hz, 1H), 4.35 (dd, *J* = 11.8, 3.7 Hz, 1H), 4.16 (dd, *J* = 11.8, 6.3 Hz, 1H), 3.72 (dd, *J* = 5.4, 1.7 Hz, 2H), 2.30 (t, *J* = 7.5 Hz, 4H), 2.00 (m, 4H), 1.60 (m, 4H), 1.33 – 1.21 (m, 44H), 0.95 (t, *J* = 7.9 Hz, 9H), 0.88 (t, *J* = 6.7 Hz, 6H), 0.59 (q, *J* = 8.0 Hz, 6H). $^{13}$C NMR (101 MHz, $CDCl_3$) δ 173.27 (2C), 129.98 (2C), 71.87, 62.62, 61.36, 34.75 (2C), 32.04 (2C), 30.24 – 28.69 (18C), 27.37 (2C), 25.03 (2C), 22.81 (2C), 14.24 (2C), 6.80 (3C), 4.44 (3C). Low-intensity signals of the sn1 acyl chain (non-enriched carbon-13) are overlapping with high-intensity labeled carbon-13 signals of the sn2 acyl chain.

ESI-MS: m/z $[M + H]^+$ calcd for $C_{25}{}^{13}C_{18}H_{85}O_5Si$: 727.68; found: 727,72; m/z $[M + Na]^+$ calcd for $C_{25}{}^{13}C_{18}H_{84}NaO_5Si$: 749.66; found: 749,35.

$[\alpha]_D^{20}$ = +7.49 (c = 0.01 g/mL, $CHCl_3$).

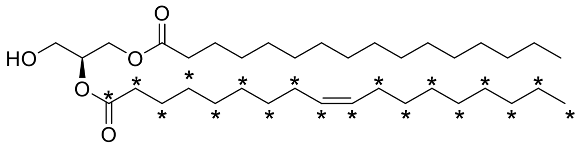

**1d** $* = {}^{13}C$

**Compound 1d.** To a solution of compound **1c** (890 mg, 1.22 mmol) dissolved in DCM (8.9 mL) was added a solution of $FeCl_3$ x $6H_2O$ (5 mM) in MeOH/DCM (3:1, 12 mL). The reaction was stirred at rt for 1 h. The mixture was then extracted once with EtOAc/$H_2O$ (1:1, 400 mL). The organic phase was washed with $H_2O$ (150 mL) and brine (150 mL), dried over anhydrous sodium sulfate, filtered and evaporated under reduced pressure. The residue was purified by chromatography (cyclohexane/EtOAc gradient, 9:1 to 8.5:1.5) to yield compound **1d** as a colorless oil (688 mg, 1.12 mmol, 92%).

${}^{1}H$ decoupled ${}^{13}C$ NMR (400 MHz, $CDCl_3$) δ 5.54 – 5.18 (m, 2H), 5.08 (p, *J* = 5.1 Hz, 1H), 4.32 (dd, *J* = 11.9, 4.6 Hz, 1H), 4.24 (dd, *J* = 11.9, 5.6 Hz, 1H), 3.73 (m, 2H), 2.32 (m, 4H), 2.01 (m, 4H), 1.61 (m, 4H), 1.28 (m, 44H), 0.88 (t, *J* = 6.7 Hz, 6H). ${}^{13}C$ NMR (101 MHz, $CDCl_3$) δ 173.83 (2C), 129.99 (2C), 72.26, 62.12, 61.73, 34.86 (2C), 32.04 (2C), 30.33 – 28.93 (18C), 27.35 (2C), 25.04 (2C), 22.81 (2C), 14.07 (2C). Low-intensity signals of the sn1 acyl chain (non-enriched carbon-13) are overlapping with high-intensity labeled carbon-13 signals of the sn2 acyl chain.

ESI-MS: m/z $[M + Na]^+$ calcd for $C_{19}{}^{13}C_{18}H_{70}NaO_5$: 635.57; found: 635.44; m/z $[M + OAc]^-$ calcd for $C_{21}{}^{13}C_{18}H_{73}O_7$: 671.60; found: 671.70.

$[\alpha]_D^{20}$ = -2.82 (c = 0.01 g/mL, $CHCl_3$).

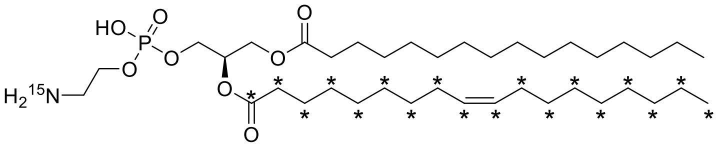

**1f**

**${}^{15}N{}^{13}C$-PE34:1** $* = {}^{13}C$

**Compound 1f.** To a solution of compound **1d** (380 mg, 0.62 mmol) and dry TEA (173 µL, 1.24 mmol) dissolved in dry toluene (8 mL) was added dropwise at 0°C a solution of 2-chloro-2-oxo-1,3,2-dioxaphospholane (97 mg, 0.68 mmol). The resulting mixture was stirred at rt overnight. The obtained white precipitate was filtered off over and the filtrate was evaporated while keeping the temperature at 20°C. The resulting residue (380 mg) was transferred with dry ACN (11 mL) into a microwave tube purged with argon. The tube was sealed and cooled down at 0°C. Dry ${}^{15}NH_3$ was bubbled into the tube until the solution became white (2 min). The mixture was heated at 65°C for 24 h. The reaction mixture was cooled down to rt and evaporated under reduced pressure. The residue was purified by chromatography (cyclohexane/EtOAc/$H_2O$ gradient 9.75:0.25:0 to 8:2:0.01) to yield compound **1f** (*i.e.* ${}^{15}N{}^{13}C$-PE34:1) as a white wax (32 mg, 0.38 mmol, 6%).

$^{1}H$ decoupled $^{13}C$ NMR (400 MHz, $CDCl_3$) δ 8.32 (br s, 3H), 5.33 (m, 2H), 5.21 (m, 1H), 4.37 – 3.84 (m, 6H), 3.17 (m, 2H), 2.29 (m, 4H), 2.00 (m, 4H), 1.58 (m, 4H), 1.28 – 1.25 (s, 44H), 0.88 (t, *J* = 7.0 Hz, 6H). $^{13}C$ NMR (101 MHz, $CDCl_3$) δ 173.01 (2C), 129.94 (2C), 70.41, 64.08, 62.73, 62.29, 40.56, 34.29 (2C), 32.05 (2C), 30.24 – 29.10 (18C), 27.38 (2C), 25.07 (2C), 22.82 (2C), 14.39 (2C). Low-intensity signals of the sn1 acyl chain (non-enriched carbon-13) are overlapping with high-intensity labeled carbon-13 signals of the sn2 acyl chain.

$^{31}P$ (162 MHz, $CDCl_3$) δ -1.37.

HR-ESI-MS (neg): m/z $[M - H]^-$ calcd for $C_{21}{}^{13}C_{18}H_{75}{}^{15}NO_8P$: 735.5804; found: 735.5835.

$[\alpha]_D^{20}$ = +4.5 (c = 0.01 g/mL, $CHCl_3$); commercial (*R*)-PE34:1 $[\alpha]_D^{20}$ = +5.1 (c = 0.01 g/mL, $CHCl_3$).

#### 2. Synthesis of cages

The 7-(diethylamino)-4-(hydroxymethyl)coumarin used for synthesis of ER-PE probe is commercial.

##### 2.i. Synthesis of TPP-Cou-OH

(3-Aminopropyl)triphenylphosphonium bromide, 7-[(carboxymethyl)-methylamino]-4-hydroxymethyl)coumarin and TPP-Cou-OH were synthesized according to previously described protocols.[1,2]

##### 2.ii. Synthesis of *pfb*-Cou-OH and *tfb*-Cou-OH (compounds **2b** and **2b'**)

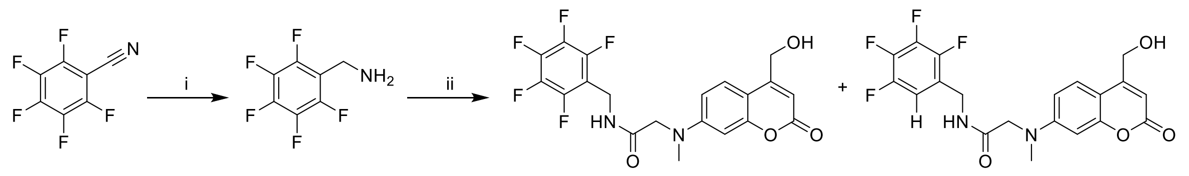

**2a** **2b** **2b'**

Scheme S1. Synthesis of *pfb*- and *tfb*-Cou-OH (compounds **2b** and **2b'**). (i) pentafluorobenzonitrile, $BH_3$.THF, THF, 66°C, 12 h - 77%; (ii) 7-[(carboxymethyl)-methylamino]-4-(hydroxymethyl)coumarin,[2] DIEA, HOBt, EDC.HCl, DCM, DMF, rt, 12 h - 66%.

**Compound 2a**. To a solution of commercial pentafluorobenzonitrile (1.0 g, 5.18 mmol) in anhydrous THF (80 ml) was added dropwise a solution of $BH_3$.THF 1M (26 ml, 26 mmol). The resultant solution was stirred and heated to reflux overnight. Then the solution was cooled down, 2.6 M HCl (100 ml) was carefully added and heating was continued at reflux for 30 min. THF was evaporated under reduced pressure. Excessive HCl solution (50 ml) was added and was extracted with $Et_2O$ (3 x 200 ml). The aqueous phase was made alkaline with a NaOH solution (10 % in $H_2O$) up to pH 10–11 and was again extracted with $Et_2O$ (3 x 100 mL). The combined organic layers were washed with brine, dried over anhydrous sodium sulfate, filtered and evaporated under reduced pressure to give pentafluorobenzylamine **2a** (783 mg, 3.97 mmol, 77%) as a yellow liquid which was used without further purification.

$^{1}$H NMR (400 MHz, $CDCl_3$) δ 3.93 (d, *J* = 17.3 Hz, 2H), 1.50 (s, 2H). $^{19}$F NMR (282 MHz, $CDCl_3$) δ -145.50 (m), -156.21 (m), -162.80 (m). $^{13}$C signals stemming were not well resolved due to the presence of complex fluorine-carbon coupling patterns.

LC-ESI-MS: m/z $[M + H]^+$ calcd for $C_7H_5F_5N$: 198.03; found: 197.89.

**Compounds 2b-2b'**. To a solution of 7-[(carboxymethyl)-methylamino]-4-(hydroxymethyl)coumarin[2] (70 mg, 0.27 mmol) dissolved in dry DCM (5 mL) with one drop of DMF were successively added pentafluorobenzylamine **2a** (105 mg, 0.53 mmol), DIEA (313 µL, 1.80 mmol), HOBt (72 mg, 0.53 mmol) and EDC.HCl (102 mg, 0.53 mmol). The reaction mixture was stirred overnight at rt under nitrogen. Then 50 mL of DCM was added and the reaction mixture was washed with saturated ammonium chloride solution (2 x 50 mL). The organic layer was dried over anhydrous sodium sulfate, filtered and evaporated under reduced pressure. The residue was purified by chromatography (DCM/MeOH gradient, 9.9:0.1 to 9.5:0.5) to yield a mixture (1:1.4) of the two compounds *pfb*-Cou-OH **2b** and *tfb*-Cou-OH **2b'** respectively as a yellow powder (78 mg, 0.18 mmol, 66%).

**Data for compound 2b**

$^{1}$H NMR (400 MHz, $CD_3OD$) δ 7.45 (br s, 1H), 7.24 (d, *J* = 9 Hz, 1H), 6.46 (dd, *J* = 9.0, 2.5 Hz, 1H), 6.37 (d, *J* = 2.4 Hz, 1H), 6.17 (s, 1H), 4.64 (s, 2H), 4.38 (s, 2H), 3.91 (s, 2H), 3.21 (br s, 1H), 3.01 (s, 3H). $^{13}$C NMR (101 MHz, $CD_3OD$) δ 170.08, 163.08, 156.36, 154.93, 151.71, 146.26, 143.23, 123.89, 121.84, 111.24 (2C, $CF_{ortho}$), 109.11, 107.59, 105.21, 98.31, 59.33, 55.55, 39.08, 30.68. LC-ESI-MS: m/z $[M + H]^+$ calcd for $C_{20}H_{16}F_5N_2O_4$: 443.10; found: 443.15.

**Data for compound 2b'**

$^{1}$H NMR (400 MHz, $CD_3OD$) δ 7.45 (br s, 1H), 7.24 (d, *J* = 9 Hz, 1H), 6.82 (m, 1H, $CH_{ortho}$), 6.51 (dd, *J* = 8.9, 2.5 Hz, 1H), 6.43 (d, *J* = 2.4 Hz, 1H), 6.17 (s, 1H), 4.64 (s, 2H), 4.30 (s, 2H), 3.96 (s, 2H), 3.21 (br s, 1H), 3.05 (s, 3H). $^{13}$C NMR (101 MHz, $CD_3OD$) δ 170.49, 163.05, 156.36, 154.96, 151.64, 147.65, 145.28, 140.48, 137.86, 123.99, 121.84, 110.28 (dt, $J^{13}C\text{-}^{19}F$ = 20.2, 3.6 Hz, $CH_{ortho}$), 109.14, 107.67, 105.29, 98.42, 59.33, 55.79, 39.19, 35.66. LC-ESI-MS: m/z $[M + H]^+$ calcd for $C_{20}H_{17}F_4N_2O_4$: 425.11; found: 425.16.

3. Synthesis of photocaged $^{15}N^{13}C$-PE34:1 probes

3.i. Synthesis of MitoPE (compound **3a**)

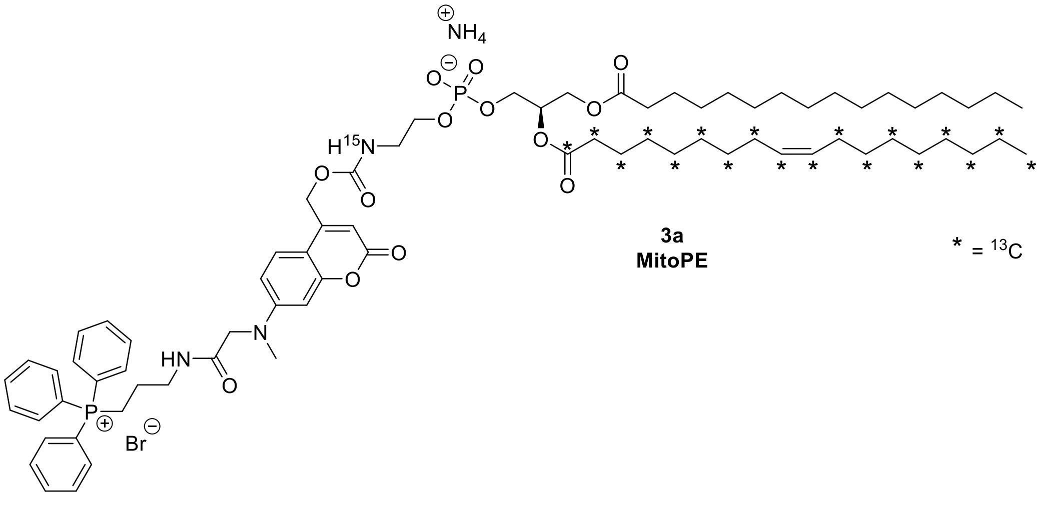

**Compound 3a.** To a solution of TPP-Cou-OH (33 mg, 0.051 mmol)[2] dissolved in dry DCM (5 mL) were added bis-(4-nitrophenyl)carbonate (16 mg, 0.051 mmol) and DIEA (27 µL, 0.153 mmol). The reaction mixture was stirred at rt in the dark. After 3 h, $^{15}N^{13}C$-PE34:1 (compound **1f**, 30 mg, 0.037 mmol) dissolved in dry DCM (2.5 mL) was added and the reaction was stirred at 38°C for 6 h in the dark. The reaction mixture was cooled down to rt and evaporated under reduced pressure. The residue was purified by two successive chromatography steps (first column: DCM/MeOH gradient 9.5:0.5 to 9:1; second column: DCM/MeOH/$NH_4OH$ gradient 9.5:0.5:0.02 to 8:2:0.02) to yield compound **3a** (*i.e.* MitoPE) as a yellow oil (20 mg, 0.015 mmol, 41%).

$^{1}H$ decoupled $^{13}C$ NMR (400 MHz, $CDCl_3$) δ 9.71 (br s, 1H), 7.75 (m, 3H), 7.69 – 7.56 (m, 12H), 6.88 (m, 1H), 6.69 (m, 1H), 6.35 (m, 1H), 5.81 (m, 1H), 5.49 – 5.05 (m, 3H), 4.84 (m, 2H), 4.46 – 3.86 (m, 8H), 3.43 (m, 6H), 3.13 (s, 3H), 2.25 (t, J = 7.5 Hz, 4H), 1.99 (m, 4H), 1.85 (m, 2H), 1.56 (m, 4H), 1.24 (m, 44H), 0.87 (t, J = 6.9 Hz, 6H). $^{13}C$ NMR (126 MHz, $CDCl_3$) δ 173.43 (2C), 170.50, 161.58, 155.98, 155.50, 152.35, 150.61, 135.22 (d, $J(^{13}C\text{-}^{31}P)$ = 2.6 Hz, 3C), 133.53 (d, $J(^{13}C\text{-}^{31}P)$ = 10.0 Hz, 6C), 130.60 (d, $J(^{13}C\text{-}^{31}P)$ = 12.9 Hz, 6C), 129.96 (2C), 124.17, 118.38 (d, $J(^{13}C\text{-}^{31}P)$ = 86.1 Hz, 3C), 109.63, 106.56, 106.09, 98.06, 70.68, 63.99, 63.71, 62.90, 61.65, 55.39, 43.11, 40.18, 38.86 (d, $J(^{13}C\text{-}^{31}P)$ = 17.6 Hz, 1C), 34.30 (2C), 32.02 (2C), 30.20 – 29.02 (18C), 27.28 (2C), 25.01 (2C), 22.79 (2C), 22.50, 20.15 (d, $J(^{13}C\text{-}^{31}P)$ = 52.7 Hz, 1C), 14.10 (2C). Low-intensity signals of the sn1 acyl chain (non-enriched carbon-13) are overlapping with high-intensity labeled carbon-13 signals of the sn2 acyl chain.

$^{31}P$ (162 MHz, $CDCl_3$) δ 24.35, 1.02.

HR-ESI-MS (pos): m/z $[M]^+$ calcd for $C_{56}{}^{13}C_{18}H_{108}N_2{}^{15}NO_{13}P_2$: 1327.7931; found: 1327.7983; m/z $[M - H + Na]^+$ calcd for $C_{56}{}^{13}C_{18}H_{107}NaN_2{}^{15}NO_{13}P_2$ : 1349.7751; found 1349.7759.

3.ii. Synthesis of ER-PE (compound **3b**)

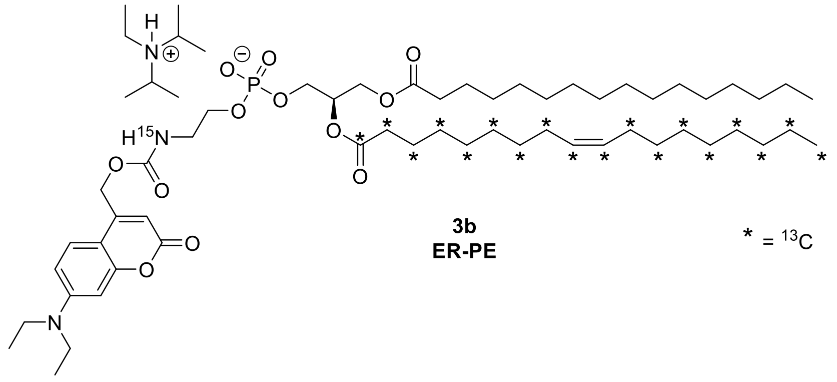

**Compound 3b.** To a solution of commercial 7-(diethylamino)-4-(hydroxymethyl)coumarin (19 mg, 0.077 mmol) dissolved in dry THF (2.4 mL) were added at 0°C triphosgene (42 mg, 0.142 mmol) and DIEA (71 µL, 0.410 mmol). The reaction mixture was stirred at 0°C in the dark. After 4 h, the reaction mixture was extracted once with EtOAc/$H_2O$ (1:1, 20 mL), the layers were separated and the organic layer was washed with brine (2 x 20 mL), dried over anhydrous sodium sulfate, filtered and evaporated under reduced pressure. The resulting 7-(diethylamino)-coumarin-4-yl)-methyl chloroformate was immediately used without further purification.

$^{15}N^{13}C$-PE34:1 (compound **1f**) (15 mg, 0.018 mmol) was dissolved in dry DCM (1.2 mL) and cooled down to 0°C. DIEA (44 µL, 0.253 mmol) was added and the reaction mixture was stirred for 10 min. Then crude 7-(diethylamino)-coumarin-4-yl)-methyl chloroformate in dry DCM (0.75 mL) was added and the solution was stirred overnight at rt. The reaction mixture was washed with saturated ammonium chloride solution (3 x 10 mL), dried over anhydrous sodium sulfate, filtered and evaporated under reduced pressure. The residue was purified by two successive chromatography steps (first column: DCM/MeOH gradient 10:0 to 9:1; second column: DCM/MeOH/$H_2O$ gradient 9.25:0.75:0.02 to 8:2:0.02) to yield compound **3b** (*i.e.* ER-PE) as a yellow oil (4.3 mg, 0.004 mmol, 22%).

$^{1}H$ decoupled $^{13}C$ NMR (400 MHz, $CDCl_3$) δ 7.46 (br s, 1H), 7.29 (m, 1H), 6.57 (m, 1H), 6.49 (m, 1H), 6.12 (m, 1H), 5.33 (m, 2H), 5.25 (m, 3H), 4.41 – 3.92 (m, 6H), 3.44 (m, 2H), 3.41 (m, 4H), 2.27 (m, 4H), 2.00 (m, 4H), 1.57 (m, 4H), 1.25 (m, 44H), 1.20 (t, *J* = 6.4 Hz, 6H), 0.87 (t, *J* = 6.5 Hz, 6H). $^{13}C$ NMR (126 MHz, $CDCl_3$) δ 173.52 (2C), 162.05, 156.36, 154.77, 150.87, 149.27, 129.96 (2C), 124.55, 108.92, 106.26, 105.91, 97.92, 70.46, 68.23, 64.77, 63.89, 62.76, 44.90 (2C), 42.37, 34.29 (2C), 32.04 (2C), 30.22 – 29.05 (18C), 27.37 (2C), 25.02 (2C), 22.82 (2C), 14.11 (2C), 12.59 (2C). Low-intensity signals of the sn1 acyl chain (non-enriched carbon-13) are overlapping with high-intensity labeled carbon-13 signals of the sn2 acyl chain.

$^{31}P$ (162 MHz, $CDCl_3$) δ -1.28.

HR-ESI-MS (neg): m/z $[M - H]^-$ calcd for $C_{36}{}^{13}C_{18}H_{90}N^{15}NO_{12}P$: 1008.6805; found: 1008.6700.

3.iii. Synthesis of *tpf*ER-PE (compound **3c** and **3c'**)

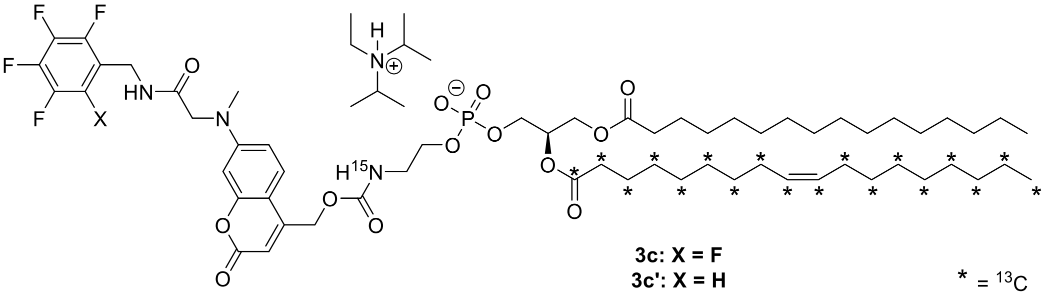

Compounds **3c** and **3c'**. To a solution of compounds **2b** and **2b'** (21 mg, 0.047 mmol) dissolved in dry DCM (4 mL) were added bis-(4-nitrophenyl)carbonate (14 mg, 0.046 mmol) and DIEA (33 µL, 0.188 mmol). The reaction mixture was stirred at rt in the dark. After 1 night, $^{15}N^{13}C$-PE34:1 (compound **1f**) (21 mg, 0.028 mmol) dissolved in dry DCM (1.9 mL) was added and the reaction was stirred at 39°C for 24 h in the dark. The reaction mixture was washed with saturated ammonium chloride solution (3 x 10 mL), dried over anhydrous sodium sulfate, filtered and evaporated under reduced pressure. The residue was purified by chromatography (DCM/MeOH gradient 10:0 to 9:1) to yield *tpf*ER-PE as a mixture (1:1.5) of the two compounds **3c** and **3c'** respectively as a yellow oil (16 mg, 0.012 mmol, 43%).

$^{1}H$ decoupled $^{13}C$ NMR (400 MHz, $CDCl_3$)

Data for compound **3c**

δ 7.20 (m, 1H), 6.55 (m, 1H), 6.37 (s, 1H), 5.94 (s, 1H), 5.45 – 5.21 (m, 3H), 5.03 (s, 2H), 4.59 (s, 2H), 4.41 – 3.88 (m, 8H), 3.47 (m, 2H), 3.11 (s, 3H), 2.27 (m, 4H), 1.99 (m, 4H), 1.56 (m, 4H), 1.25 (m, 44H), 0.87 (t, *J* = 6.2 Hz, 6H).

Data for compound **3c'**

δ 7.20 (m, 1H), 7.04 (s, 1H) ($CH_{ortho}$), 6.55 (m, 1H), 6.37 (s, 1H), 5.94 (s, 1H), 5.45 – 5.21 (m, 3H), 5.03 (s, 2H), 4.51 (s, 2H), 4.41 – 3.88 (m, 8H), 3.47 (m, 2H), 3.11 (s, 3H), 2.27 (m, 4H), 1.99 (m, 4H), 1.56 (m, 4H), 1.25 (m, 44H), 0.87 (t, *J* = 6.2 Hz, 6H).

$^{13}C$ NMR (126 MHz, $CDCl_3$)

Due to their proximity, $^{13}C$ peaks cannot be assigned to molecule **3c** or **3c'**. Only one script is therefore represented for the mixture of molecules **3c** and **3c'**.

$^{13}C$ NMR (126 MHz, $CDCl_3$) δ 173.04 (2C), 169.84, 162.06, 156.22, 155.38, 151.99, 150.71, 129.95 (2C), 124.82, 122.16, 111.57 ($CF_{ortho}$ of compound **3c**), 111.16 (d, $J_{^{13}C-^{19}F}$ = 19.7 Hz, $CH_{ortho}$ of compound **3c'**), 109.70, 107.88, 104.49, 98.95, 70.18, 65.31, 64.31, 62.55, 61.83, 57.10, 42.05, 39.81, 36.47, 34.25 (2C), 32.04 (2C), 30.21 – 29.04 (18C), 27.31 (2C), 25.01 (2C), 22.81 (2C), 14.38 (2C). Signals stemming from the pentafluorobenzylamide structure (δ 148.09 - 136.56) were not well resolved due to the presence of complex fluorine-carbon coupling patterns. Low-intensity signals of the sn1 acyl chain (non-enriched carbon-13) are overlapping with high-intensity labeled carbon-13 signals of the sn2 acyl chain.

$^{31}$P (162 MHz, $CDCl_3$) δ -1.19.

HR-ESI-MS

Data for compound **3c**
HR-ESI-MS (neg): m/z $[M - H]^-$ calcd for $C_{42}{}^{13}C_{18}H_{88}F_5N_2{}^{15}NO_{13}P$: 1203.6549; found: 1203.6539.
Data for compound **3c’**
HR-ESI-MS (neg): m/z $[M - H]^-$ calcd for $C_{42}{}^{13}C_{18}H_{89}F_4N_2{}^{15}NO_{13}P$: 1185.6643; found: 1185.6569.

#### Cell culture

HeLa cells were cultured at 37˚C and 5% $CO_2$ in Dulbecco's Modified Eagle Medium (GLUTAMAX, DMEM, Cat# 31966021, Gibco) supplemented with 10% fetal bovine serum (FBS, Cat# 10270106, Gibco) and 1% Penicillin/Streptomycin (Cat# 10378016, Gibco). Cell numbers were quantified by Countess II Automated Cell Counter (Invitrogen) following manufacturers protocol. For photo-uncaging experiments, cells were grown in 35 mm dishes to reach ~ 80% confluency.

Cryopreserved hepatocytes derived from male mouse (Cat# MSCS10, Gibco) were thawed at 37°C, transferred into Williams' Medium E (Cat# A1217601, Gibco), centrifuged (100 x *g* rpm for 5 min), and cell pellet was directly used for ER-PE uncaging assay.

**Cell transfection**

1. Plasmid and siRNAs

The plasmid encoding human PEMT2 tagged with Myc was purchased from ORIGENE, Cat# RC223190. Plasmids were transformed into chemically competent NEB 5-alpha bacterial cells (BioLabs), purified using PureLink® HiPure Plasmid Filter Midiprep kit (Invitrogen) and sequence verified by Sanger sequencing (Fasteris SA, Plan-les-Ouates, Geneva, Switzerland).

Silencer Select small interfering RNAs (siRNAs) for *EPT1*, *CEPT1* and *CHPT1* were from siTOOLs Biotech GmbH.

2. Cell transfection

Transfection was performed using lipofection reagent (*Trans*IT-X2® Dynamic Delivery System (Mirus Bio) for plasmid transfection and RNAiMAX (Cat# 13778075) for siRNA transfection) and OptiMEM, according to the manufacturer's instructions. Typically, HeLa cells were seeded in a 35 mm dish and cultured for 1 day. PEMT2 plasmid content was 0.69 µg per dish, and the two siRNAs targeting i) *EPT1* and *CEPT1* or ii) *CHPT1* and *CEPT1* were mixed at a final concentration of 12 nM each. Twenty-four hours after the plasmid transfection and 72 h after the siRNA transfection, cells were ready for experiments.
Expression levels were determined by Western Blot for PEMT and by quantitative RT-PCR for *EPT1*, *CHPT1* and *CEPT1*.

3. PEMT immunofluorescence

HeLa cells were grown in 35 mm glass bottom MatTek dishes. The day after, cells were transfected with PEMT2 plasmids using *Trans*IT-X2® (Mirus Bio). 24 h after transfection, cells were fixed with 4% paraformaldehyde. Fixed cells were blocked and permeabilized with 1% bovine serum albumin and

0.05% saponin in PBS, and then incubated with the appropriate primary antibodies and followed by fluorescently labeled secondary antibodies. Finally, cells were stained with Hoechst DNA dye and kept in PBS for confocal imaging.

4. Western blot

Cell lysates were analyzed by western blot using the following antibodies: anti-myc (Origene, Cat# TA150121, 1:1000). After incubation with secondary antibodies coupled to HRP, the chemiluminescent signal was acquired using FUSION FX imaging system (VILBER).

5. Quantitative RT-PCR

Total RNA from cells was extracted and purified using with RNeasy Mini Kit (Qiagen) and reversely transcribed into cDNA with the iScript cDNA Synthesis Kit (Bio-Rad). Quantitative RT-PCR analysis was performed on a CFX Connect Real-Time PCR system using SsoAdvanced Universal SYBR Green Supermix (Bio-Rad). The primers used were (5'→ 3'):

*EPT1* Forward AGGTATTTCCTACTTGGCTG
*EPT1* Reverse CAGGATCAAAGTATGCCATTAG
*CEPT1* Forward GGATCAACAATAGCAGGAAC
*CEPT1* Reverse TTCTTGTAGATCATTGCAGC
*CHPT1* Forward CACAGTATTTATGGCAGTGG
*CHPT1* Reverse CATGCCTGAAACATAAGTCTG
*GAPDH* Forward GGCCATCCACAGTCTTCTG
*GAPDH* Reverse TCATCAGCAATGCCTCCTG

Fold change in transcript levels was calculated using the cycle threshold (CT) comparative method ($2^{-ddCT}$) normalizing to CT values of internal control genes *GAPDH*.

#### Cell labeling for lipidomic experiments

1. Caged PE probes

1.i. MitoPE or *tpf*ER-PE

Preparation of donor vesicles

Palmitoyl-oleoyl-phosphatidylcholine (POPC) and cholesterol (1:1) were mixed in chloroform/MeOH (1:1) and the solvent was evaporated under a nitrogen stream followed by high vacuum for 1 h. Finally, sterile PBS was added and sonicated using an ultra-sound bath for 5 min. The sample was centrifuged for 5 min at 3000 *g* to pellet any undispersed lipid and probe particles. The small unilamellar vesicles (SUVs) in supernatant were used as the donors in cell labeling.

Introduction to cells

MitoPE or *tpf*ER-PE (10 µM or 5 µM respectively) was complexed with SUVs (500 µM) and methyl-β-cyclodextrine (mβCD, 10 mM) in sterile HBSS (Cat# 14025092, Gibco) supplemented with 1% FBS by stirring for 30 min at 37°C in the dark. HeLa cells at 80% of confluency were fed with MitoPE/SUVs/mβCD complex and incubated for 15 min at 37°C in the dark, washed by HBSS and finally replaced with new HBSS (or DMEM without FBS in time course experiment) before uncaging.

1.ii. ER-PE

HeLa cells at 80% of confluency were fed with ER-PE (5 µM) in sterile HBSS supplemented with 1% FBS and incubated for 15 min at 37°C in the dark, washed by HBSS and finally replaced with new HBSS (or DMEM without FBS in time course experiment) before uncaging.

Mouse hepatocyte pellet was resuspended into Williams' Medium E containing ER-PE (5 µM) and incubated for 15 min at 37°C in the dark, washed with Williams' Medium E and resuspended in Williams' Medium E in nine 35 mm dishes before uncaging.

1.iii. Photo-uncaging

Experiments were performed using a 1000 Watt Arc Lamp Source (#66924, NewPort) equipped with a dichromic mirror (350–450 nm, #66226). Cells were placed on the ice under the lamp at a distance of 20 cm, and irradiated 120 seconds at 1000 Watt. Cells were either placed back in a 37 ºC incubator for the indicated time periods, and/or immediately harvested for lipid extraction.

2. $d_4$-ethanolamine

HeLa cells at 80% of confluency were fed with $d_4$-ethanolamine hydrochloride (Sigma, 0.5 mM in DMEM without FBS) and incubated for different time (90 min or 5 h) at 37°C. Then, the cells were washed in PBS (3x) and harvested for total lipid extraction.

#### Lipid extraction and quantification

Lipids were extracted following previously described protocols with minor modifications.[3–5] Briefly, cells were washed with cold PBS and scraped off in 500 µl cold PBS on ice. The suspension was transferred to a 2 ml Eppendorf tube in which it was spin down at 2500 rpm for 5 min at 4°C. After taking off the PBS, samples were stored at -80˚C or directly used for further extraction.

For phospholipid analysis, samples were prepared following the MTBE protocol.[6] Briefly, cell pellets were re-suspended in 100 µL of MS-grade water. 360 µL of MeOH and a mixture of internal standards (0.4 nmol of DLPC, 1 nmol of PE31:1, 1 nmol of PI31:1, 3.3 nmol of PS31:1, 2.5 nmol of C12 sphingomyelin, 0.5 nmol of C17 ceramide and 0.1 nmol of C8 glucosylceramide) were added. Samples were vortexed, following the addition of 1.2 ml of MTBE. The samples were vigorously vortexed at maximum speed for 10 min at 4 ºC and incubated for 1 h at room temperature on a shaker. Phase separation was induced by addition of 200 µL MS-grade water and incubation for 10 min. Samples were centrifuged at 1,000 g for 10 min. The upper phase was transferred into a 13 mm glass tube and the lower phase was re-extracted with 400 µL of a MTBE/MeOH/$H_2O$ mixture (10:3:1.5, v/v). The extraction was repeated one more time. The combined upper phase was dried under nitrogen flow and frozen at -80°C.

Dried samples were resuspended by sonicating in 100 µl of LC-MS-grade $CHCl_3$:MeOH (1:1, v/v). Reversed-phase UHPLC-HRMS analyses were performed using a Q Exactive Plus Hybrid Quadrupole-Orbitrap mass spectrometer coupled to an UltiMate 3000UHPLC system (Thermo Fisher Scientific) equipped with an Accucore C30 column (150 x 2.1 mm, 2.6 µm) and its 20mm guard (Thermo Fisher Scientific). Samples were kept at 8°C in the autosampler, 10 µl were injected and eluted with a gradient starting at 10% B for 1 min, 10-70% B in 4 min, 70-100% B in 10 min, washed in 100 % B for 5 min and column equilibration for an additional 3 min. Eluents were made of 5 mM ammonium formate and 0.1% formic acid in water (solvent A) or in isopropanol/acetonitrile (2:1, v/v) (solvent B). Flow rate and column oven temperature were respectively at 350µl/min and 40°C. The mass spectrometer was operated using a heated electrospray-ionization (HESI) source in positive and negative polarity. Lipid concentrations were calculated with respect to the corresponding internal standards and were presented as percentage of total PC signal.

For identification of phospholipids, parallel reaction monitoring (PRM) measurement was performed using a predetermined inclusion list of corresponding lipid species. The following setting was used in HCD fragmentation: automatic gain control: 1e6; maximum injection time: 50 ms; resolution: 70,000; (N)CE: 20. Xcalibur.4.2 (Thermo Fisher Scientific) was used for data acquisition and processing.

#### Imaging experiments

1. Cell imaging

HeLa cells were grown in 35 mm glass bottom MatTek dishes until reaching ~ 50% confluency. Cells were washed with PBS (1x) and incubated with the following caged probes and their appropriate organelle tracker in HBSS imaging buffer supplemented with 1% FBS for 15 min at 37˚C:

- MitoPE (25 µM) previously complexed with POPC/cholesterol (1:1, 500 µM) and mβCD (10 mM) (see cell labeling section), and MitoTracker Orange (50 nM)

- *tpf*ER-PE (25 µM) previously complexed with POPC/cholesterol (1:1, 500 µM) and mβCD (10 mM) (see cell labeling section), and ER-Tracker Red (1 µM)
- ER-PE (25 µM) and ER-Tracker Red (1 µM)

Finally, cells were washed and resuspended in HBSS (1 mL) before fluorescence imaging.

For optimization of mβCD concentration, the protocol described above was used with different concentrations of mβCD (0, 2.5, 5 or 10 mM).

2. Image acquisition

Subcellular localization experiments were performed on a Leica SP8 confocal microscope using a 63 x oil immersion objective. A 405 nm and 532 nm laser were used with appropriate filter settings during image acquisition. The fluorescence images were analyzed by Fiji software.[7]

### Supplementary figures S1-S9

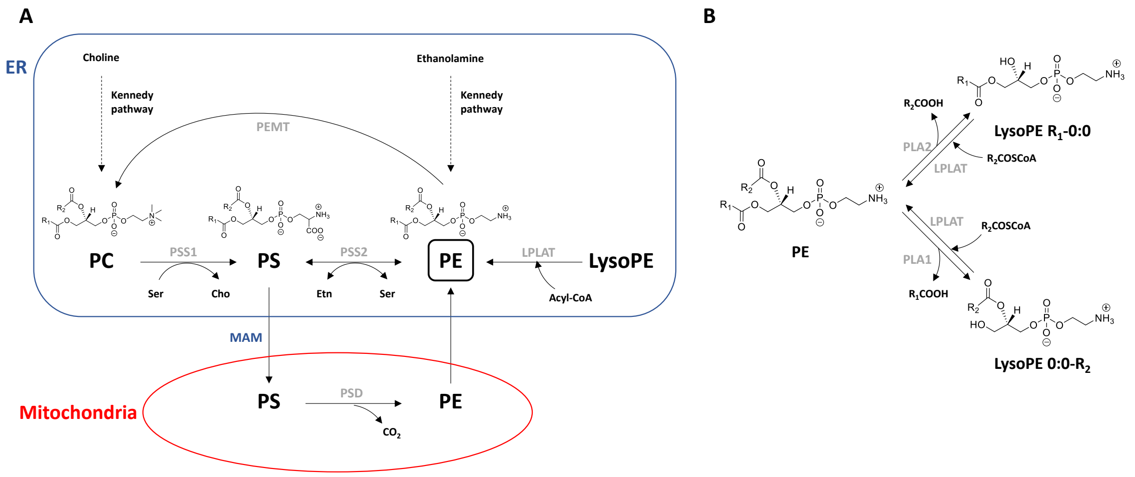

Figure S1. PE metabolism and sn1/sn2 acyl chain remodeling in mammalian cells. Cho, choline; Ser, serine; Etn, ethanolamine; PC, phosphatidylcholine; PS, phosphatidylserine; PE, phosphatidylethanolamine. Enzyme: PSS, phosphatidylserine synthase; LPLAT, lysophospholipid acyltransferase; PSD, phosphatidylserine decarboxylase; PEMT, phosphatidylethanolamine N-methyltransferase; PLA, phospholipase. PE sn1/sn2 remodeling can be transposed to other phospholipid species.

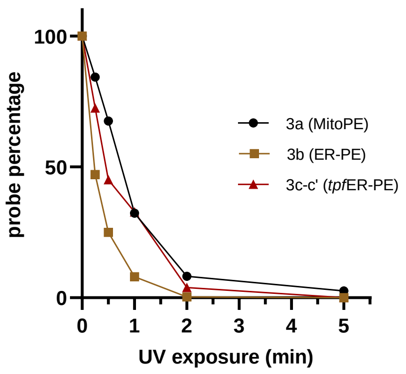

Figure S2. Uncaging efficiency of probes **3a** (MitoPE) **3b** (ER-PE) and **3c-3c'** (tpfER-PE). Each probe was dissolved in a mixture of chloroform/methanol (1:1 v/v) (~ 100 nM), illuminated by UV light (350-450 nm) on ice for 0 s, 15 s, 30 s, 1 min, 2 min and 5 min respectively and quantified by LC-MS.

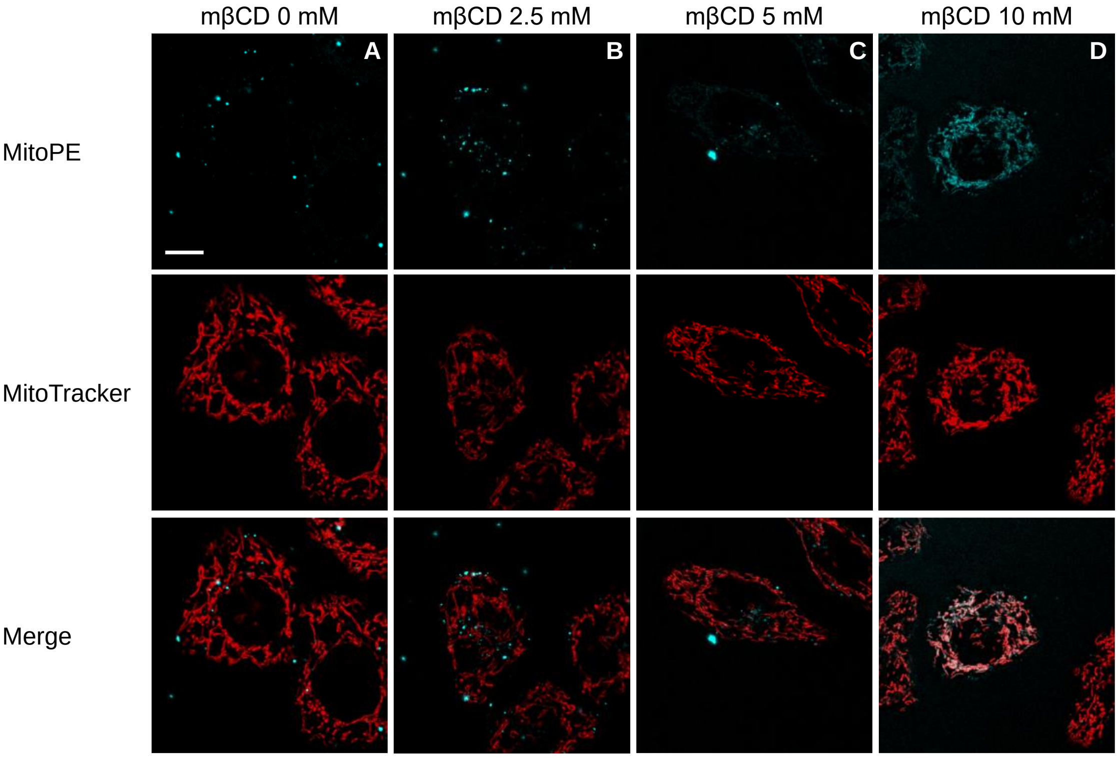

Figure S3. MitoPE incorporation in living HeLa cells, optimization of mβCD concentration. HeLa cells were treated with MitoPE (25 µM) previously complexed with liposomes (POPC/cholesterol 1:1, 500µM) and different concentrations of mβCD, in the presence of MitoTracker Orange (50 nM) for 15 min at 37°C. Then cells were washed by HBSS (1x), and replaced by HBSS prior to fluorescence imaging. Scale bar: 10 µm.

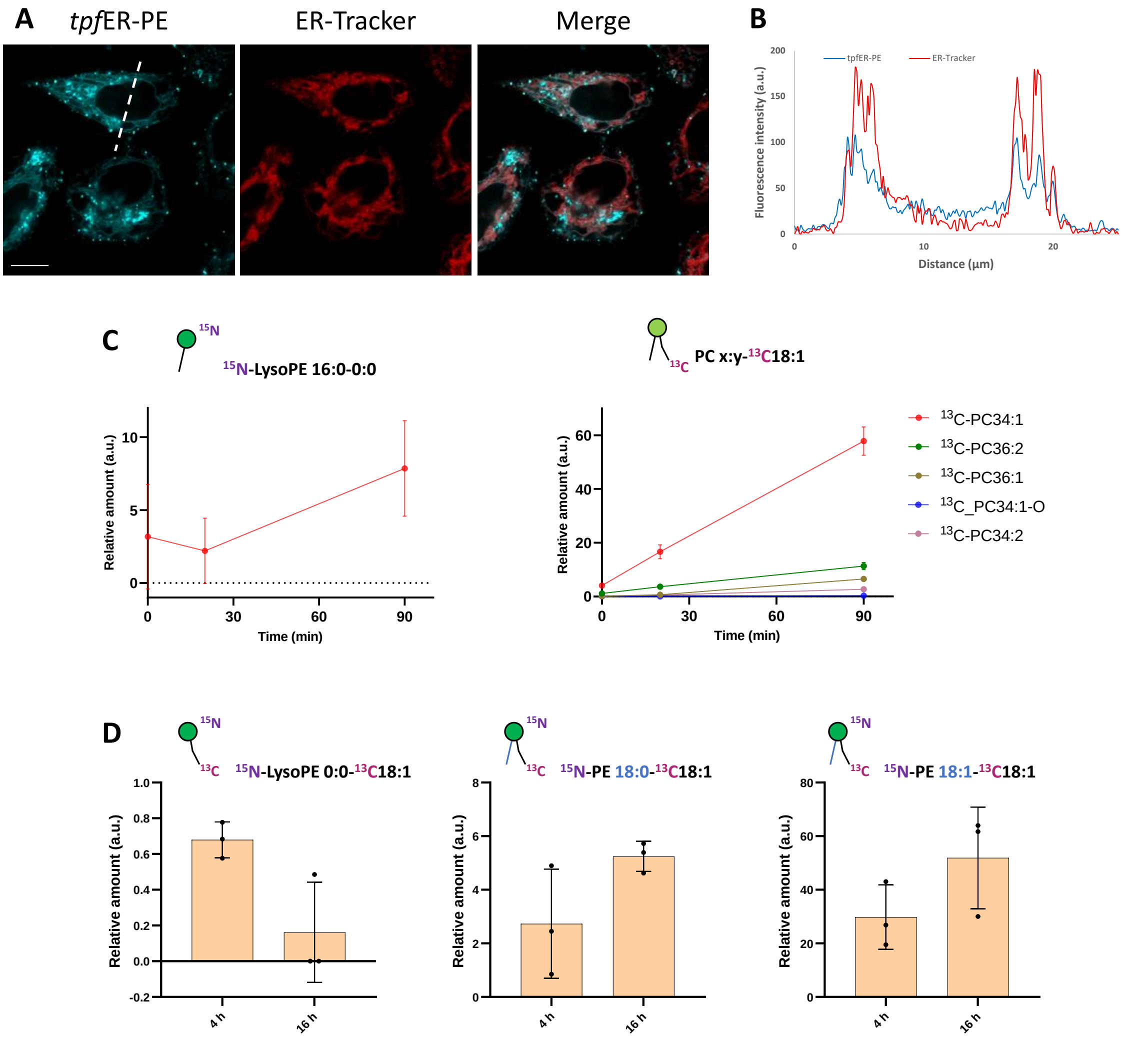

Figure S4. *tpf*ER-PE microscopy localization and lipidomics. A) Representative fluorescence image of HeLa cells stained *tpf*ER-PE (25 µM, previously complexed with cholesterol/POPC 1:1 and mβCD, see detailed description in the ESI) and ER-Tracker (1 µM) for 15 min at 37°C. Scale bar: 10 µm. B) Intensity profiles of the white dotted lines in *tpf*ER-PE and ER-Tracker channels. C-D) Photo-released $^{15}N^{13}C$-PE34:1 participates to phospholipid remodeling in living cells. *tpf*ER-PE was introduced in HeLa cells as outlined in "experimental procedures" for 15 min. Then cells were washed, irradiated by UV light for 2 min on ice, incubated at 37˚C and collected for lipid extraction at different time points: C) 0, 20 or 90 min to detect sn2 remodeling of endogenous lysoPC species; D) 4 h or 16 h to detect sn1 remodeling of photo-released $^{15}N^{13}C$-PE34:1. Values were normalized with respect to the amount of internal standards and released $^{15}N^{13}C$-PE34:1. Data represent the average of three independent experiments. Error bars represent SD.

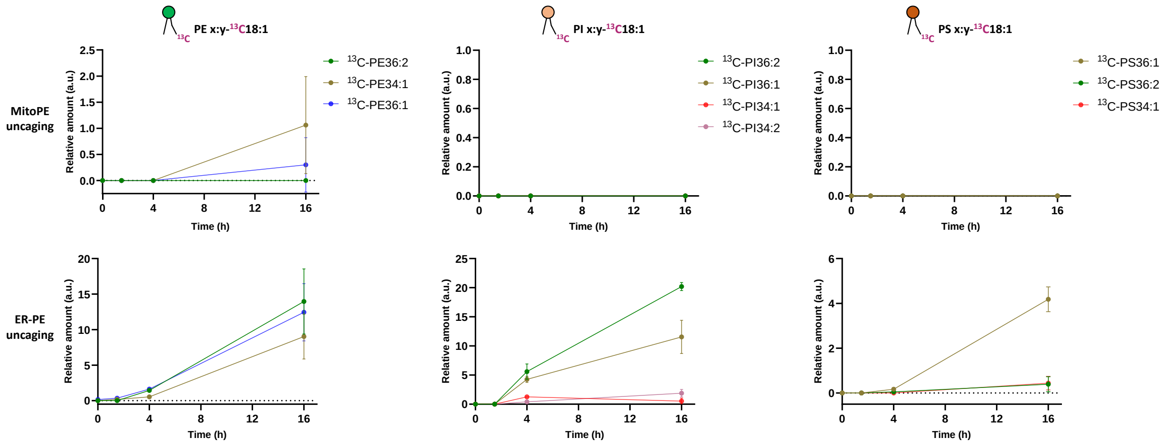

Figure S5. $^{15}N^{13}C$-PE34:1 released from ER participates to sn2 remodeling of endogenous lysoPE, lysoPI and lysoPS species in living cells. MitoPE or ER-PE were introduced in HeLa cells as outlined in "experimental procedures" for 15 min. Then cells were washed, irradiated by UV light for 2 min on ice, incubated at 37˚C, and collected for lipid extraction and LC-MS analysis at different time points (0, 1.5, 4 or 16 h). Values were normalized with respect to the amount of internal standards and photo-released $^{15}N^{13}C$-PE34:1. Data represent the average of three independent experiments. Error bars represent SD.

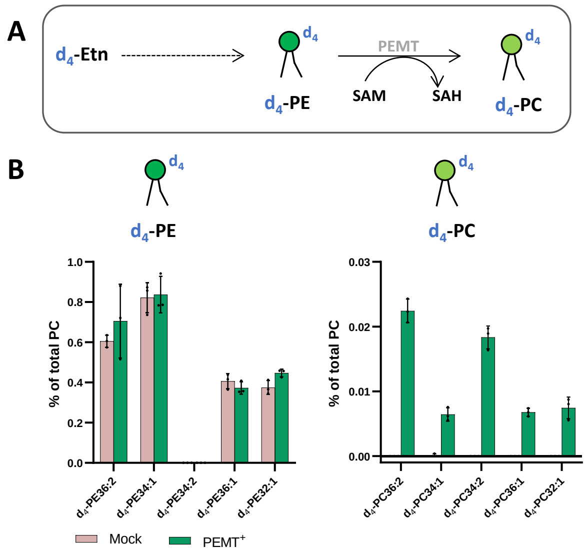

Figure S6. Incorporation of $d_4$-Etn into lipid metabolites of HeLa cells transfected with PEMT2. HeLa control cells and transiently transfected with PEMT2 were washed once with PSB and incubated with $d_4$-Etn-HCl (0.5 mM) for 1.5 h at 37°C. Then cells were washed three times with PBS and collected for lipid extraction. A) Illustration of $d_4$-PE methylation to $d_4$-PC by PEMT. B) LC-MS analysis of labeled species involved in PEMT pathway. Values were normalized with respect to the amount of internal standards and total PC species. Data represent the average of three independent experiments. Error bars represent SD.

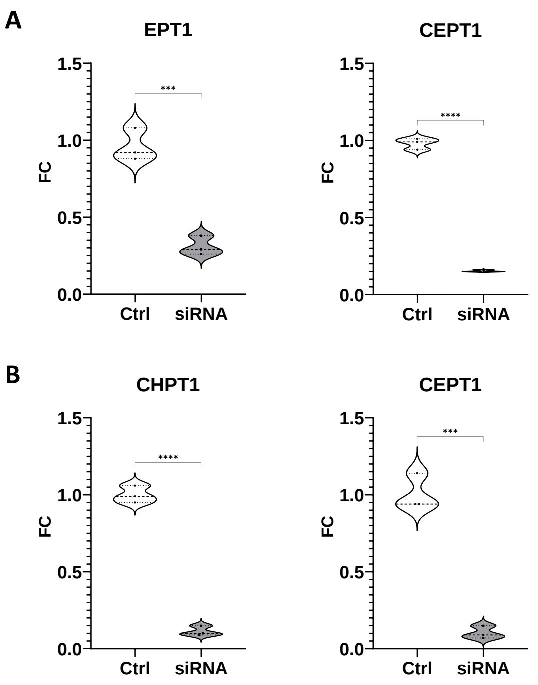

Figure S7. RT-qPCR analysis of A) *EPT1* and *CEPT1* mRNA levels in PEMT overexpressing HeLa cells transfected with negative siRNA control (Ctrl) or cotransfected with siRNAs targeting *EPT1* and *CEPT1* (siRNA); and B) *CHPT1* and *CEPT1* mRNA levels in PEMT overexpressing HeLa cells transfected with negative siRNA control (Ctrl) or cotransfected with siRNAs targeting *CHPT1* and *CEPT1* (siRNA). ***, p<0.001; ****, p<0.0001. Values are given as fold changes relative to ctrl set to value 1 (mean ± SD).

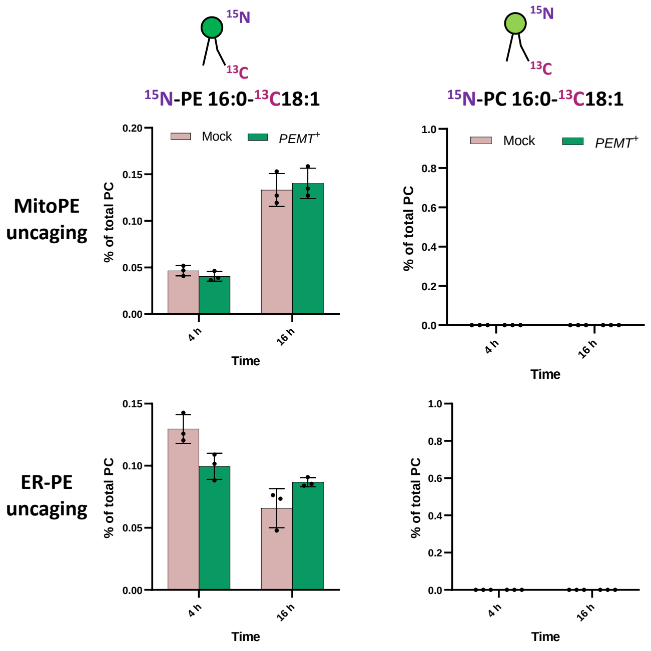

Figure S8. No detection of $^{15}N^{13}C$-PC34:1 in HeLa cells transfected with PEMT2. HeLa control cells and transiently transfected with PEMT2 were washed once with PBS and incubated with MitoPE or ER-PE as outlined in "experimental procedures" for 15 min. Then cells were washed, irradiated by UV light for 2 min on ice, incubated at 37°C, and collected for lipid extraction and LC-MS analysis at different time points. Values were normalized with respect to the amount of internal standards and total PC species. Data represent the average of three independent experiments. Error bars represent SD.

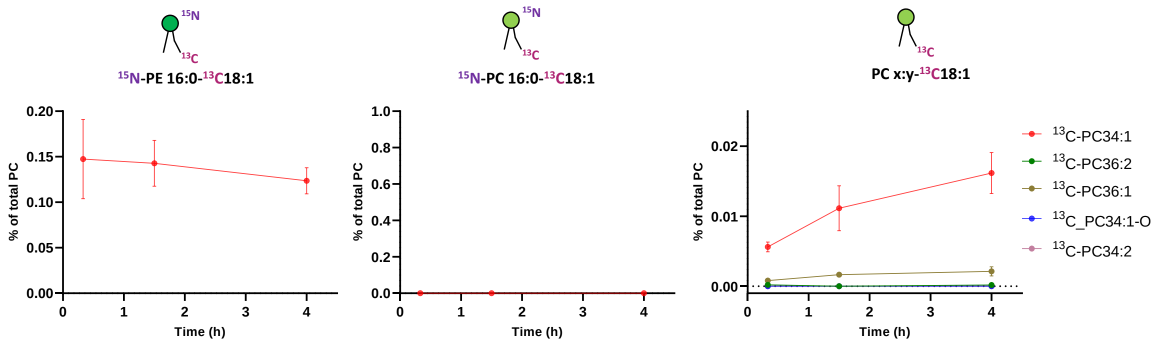

Figure S9. No detection of $^{15}N^{13}C$-PC34:1 in mouse CD1-hepatocytes. Resuspended male mouse CD1-hepatocytes were incubated with ER-PE for 15 min. Then cells were washed, irradiated by UV light for 2 min on ice, incubated at 37˚C, and collected for lipid extraction and LC-MS analysis at different time points (20 min, 90 min or 4 h). Values were normalized with respect to the amount of internal standards and total PC species.

### LC-MS/MS characterization of isotopically labeled phospholipid species

**$^{15}N^{13}C$-PE34:1** (MitoPE incorporation in HeLa cells followed by UV uncaging).

RT = 13.2 min; Positive mode

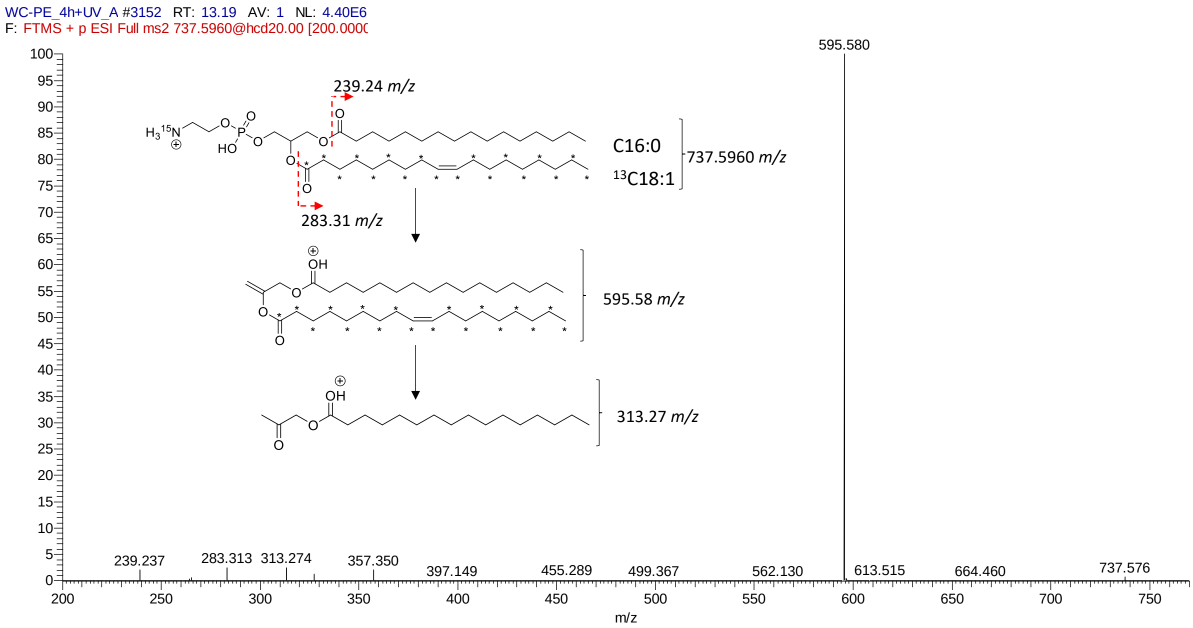

Negative mode

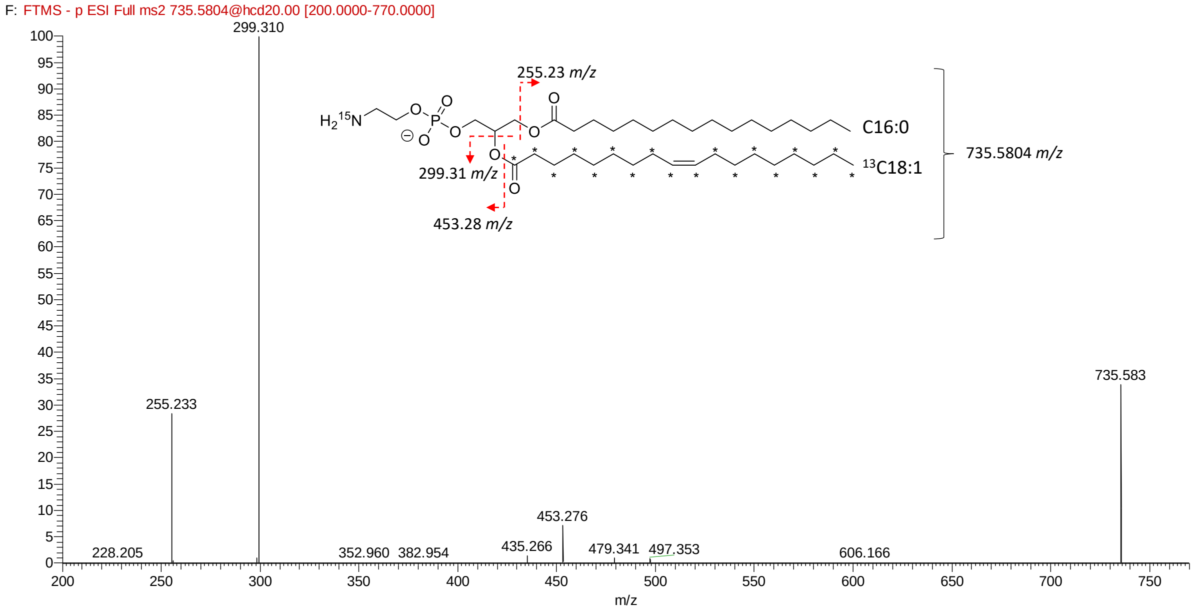

**Labeled lysoPE species** (ER-PE incorporation in HeLa cells followed by UV uncaging and chase of 90 min).

##### $^{15}$N-LysoPE 16:0-0:0

RT = 7.3 min; Negative mode

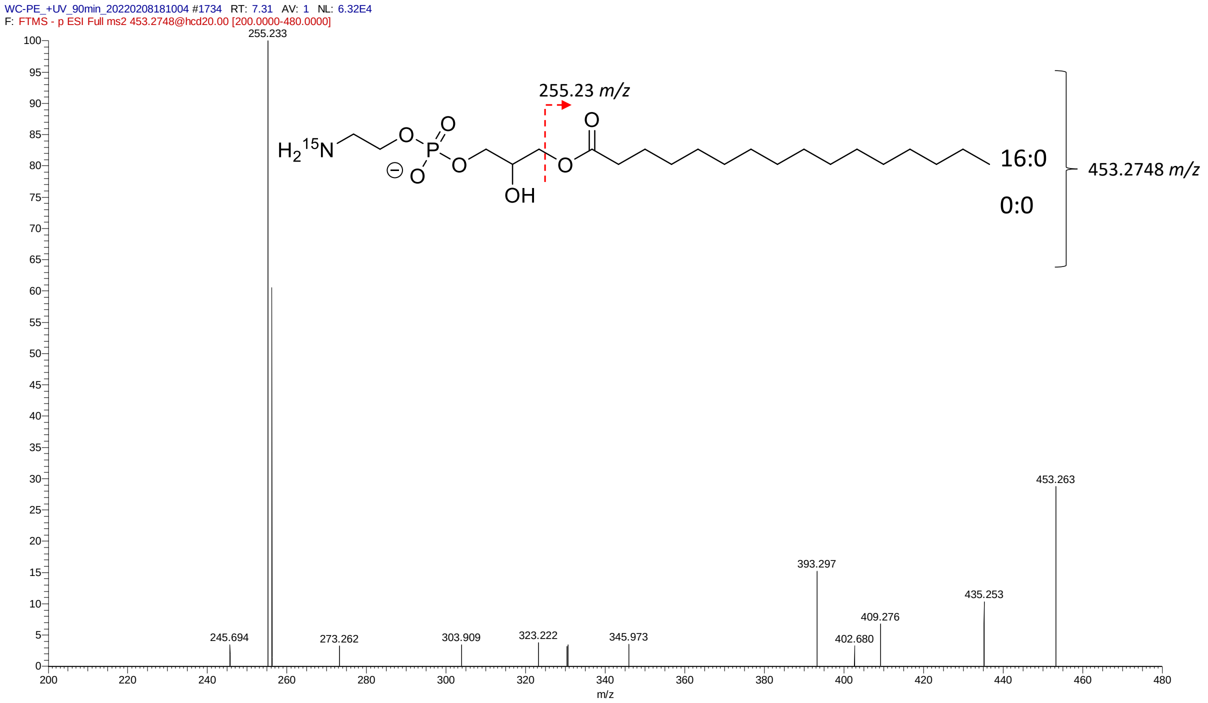

##### $^{15}$N-LysoPE0:0-$^{13}$C18:1

RT = 7.4 min; Negative mode

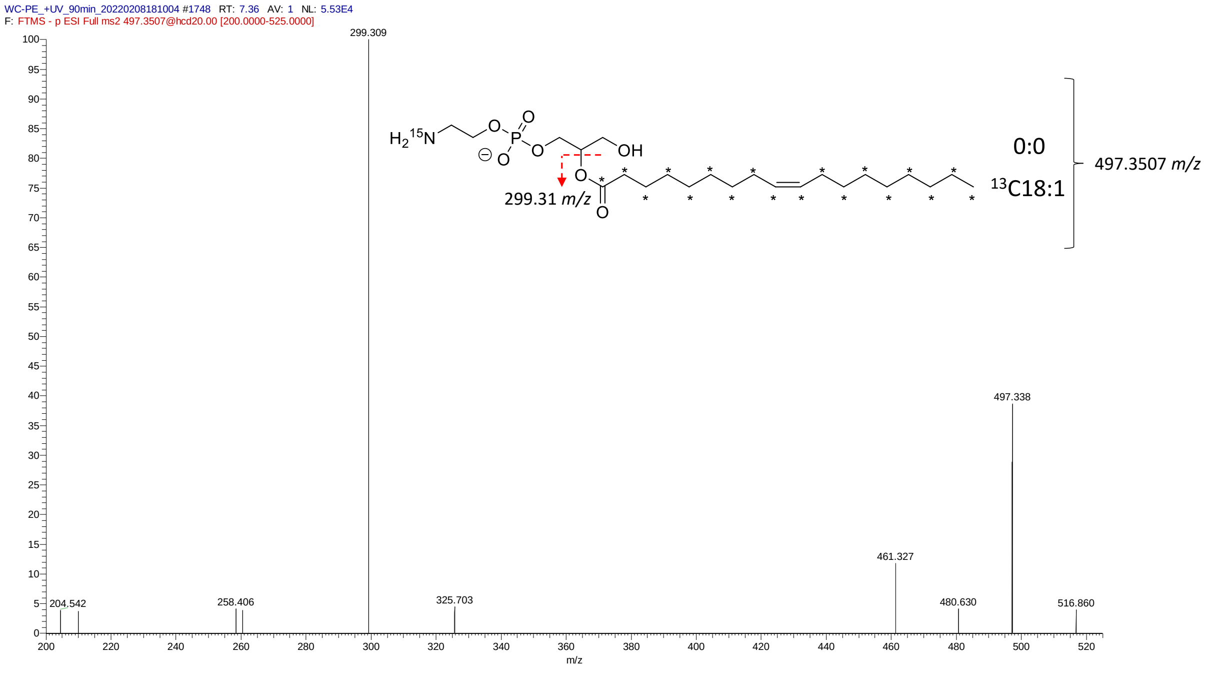

**[13]C-PC species** (ER-PE incorporation in HeLa cells followed by UV uncaging and chase of 90 min).

## [13]C-PC34:1

RT = 13.1 min; Negative mode

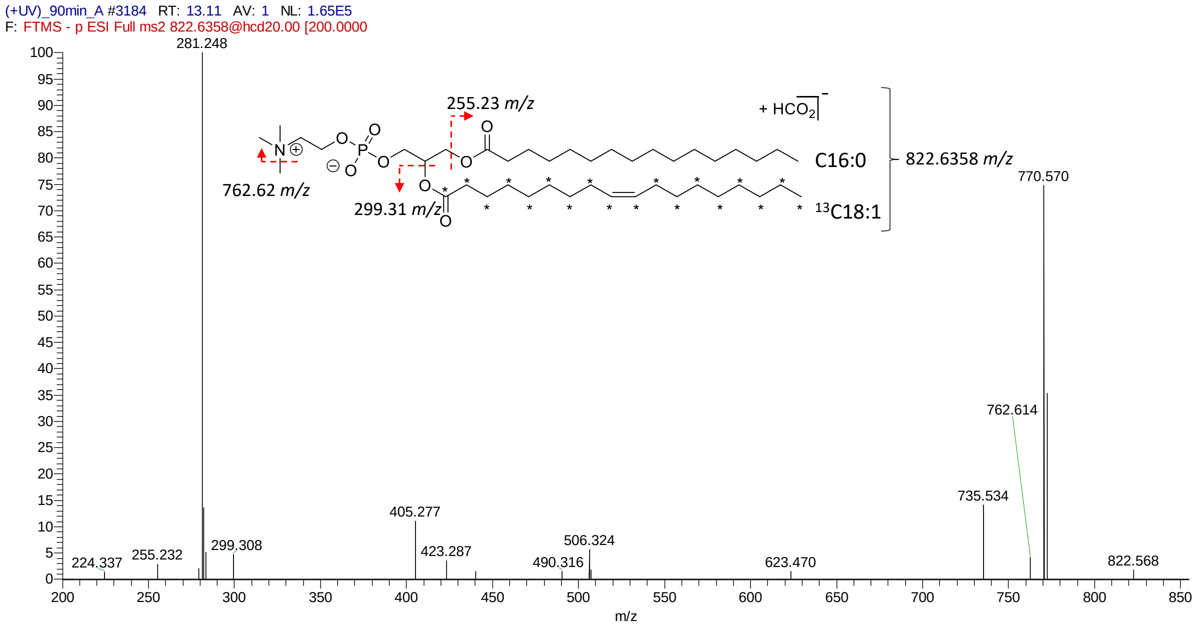

## [13]C-PC36:2

RT = 13.2 min; Negative mode

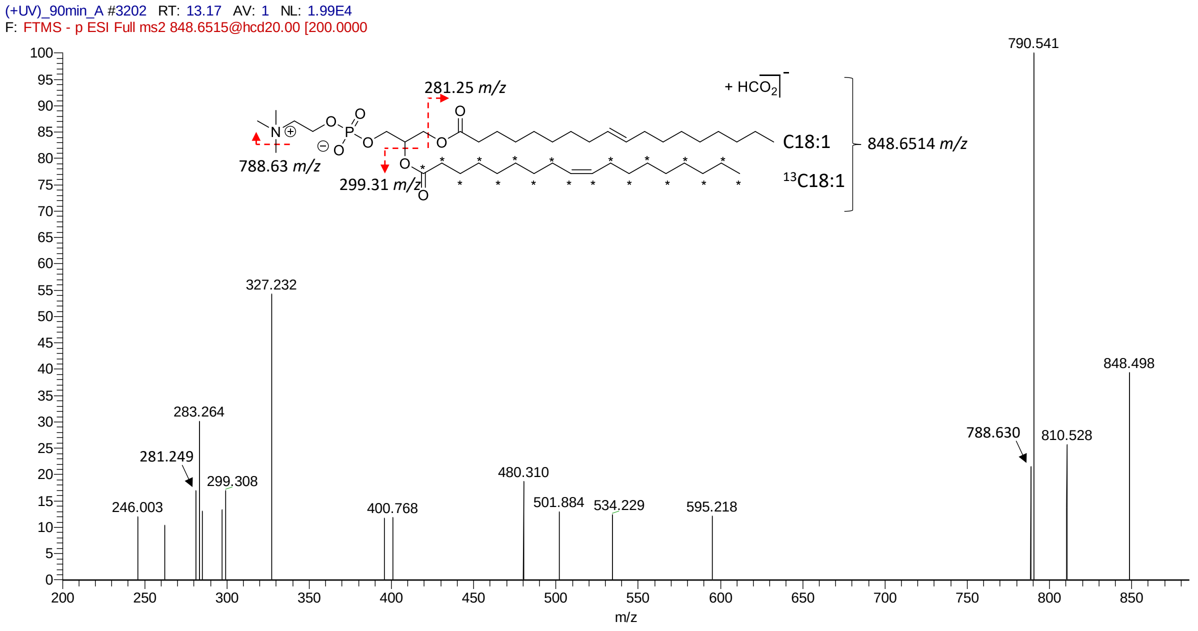

## $^{13}$C-PC36:1

RT = 13.9 min; Negative mode

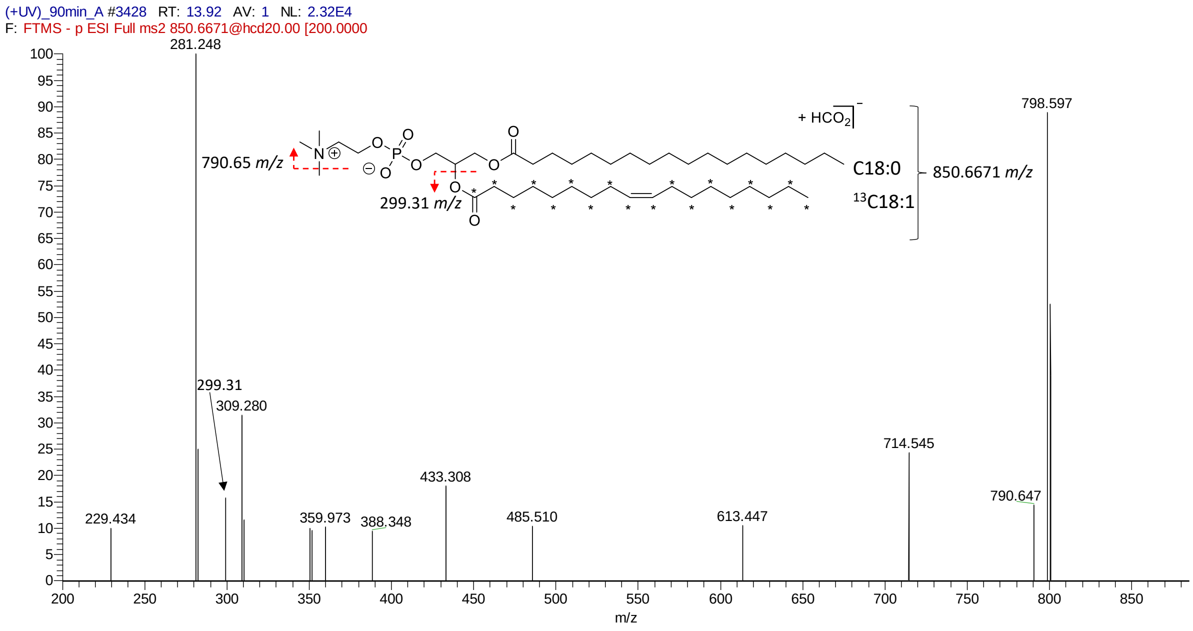

## $^{13}$C-PC34:1-O

RT = 13.6 min; Negative mode

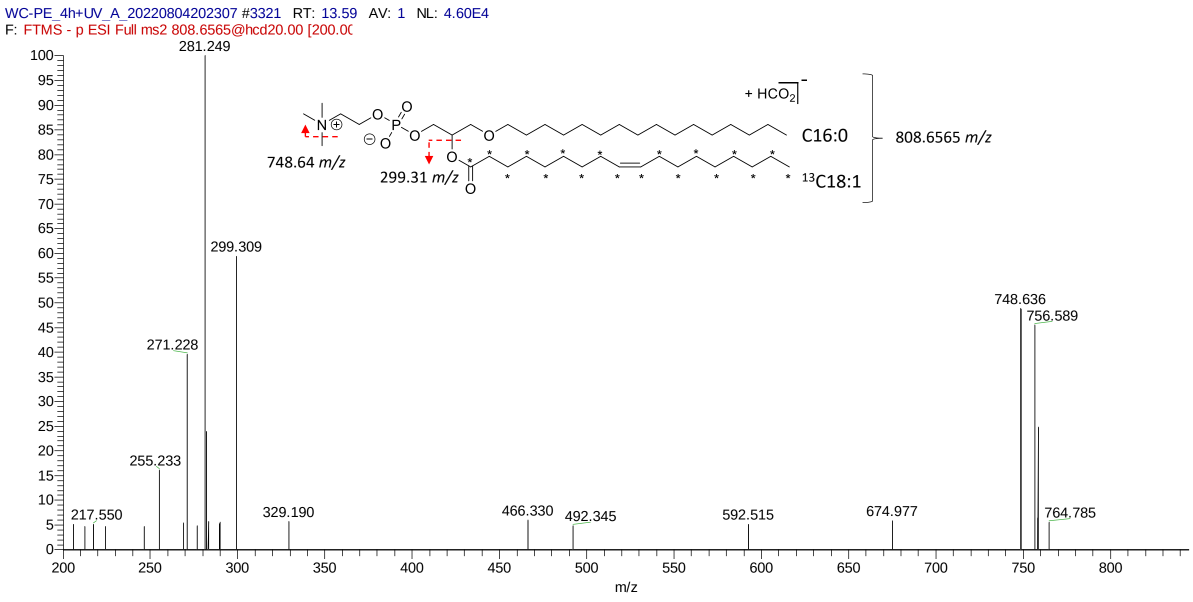

**$^{13}$C-PC34:2**

RT = 12.3 min; Negative mode

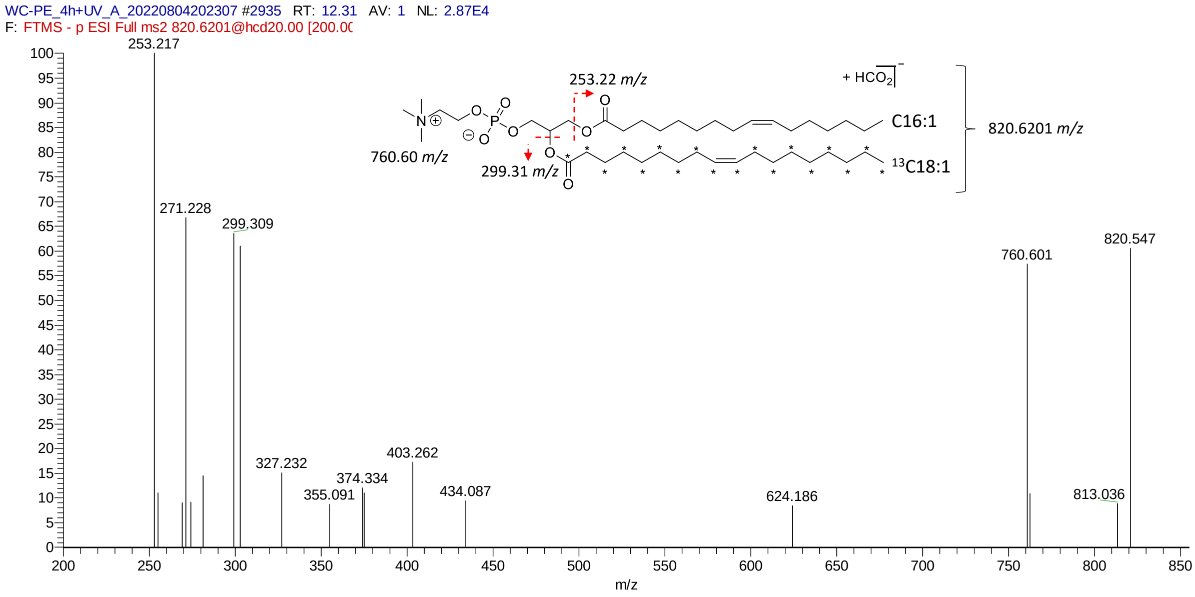

$^{13}$**C-PE species** (ER-PE incorporation in HeLa cells followed by UV uncaging and chase of 4 h).

## $^{13}$C-PE36:2

RT = 13.3 min; Negative mode

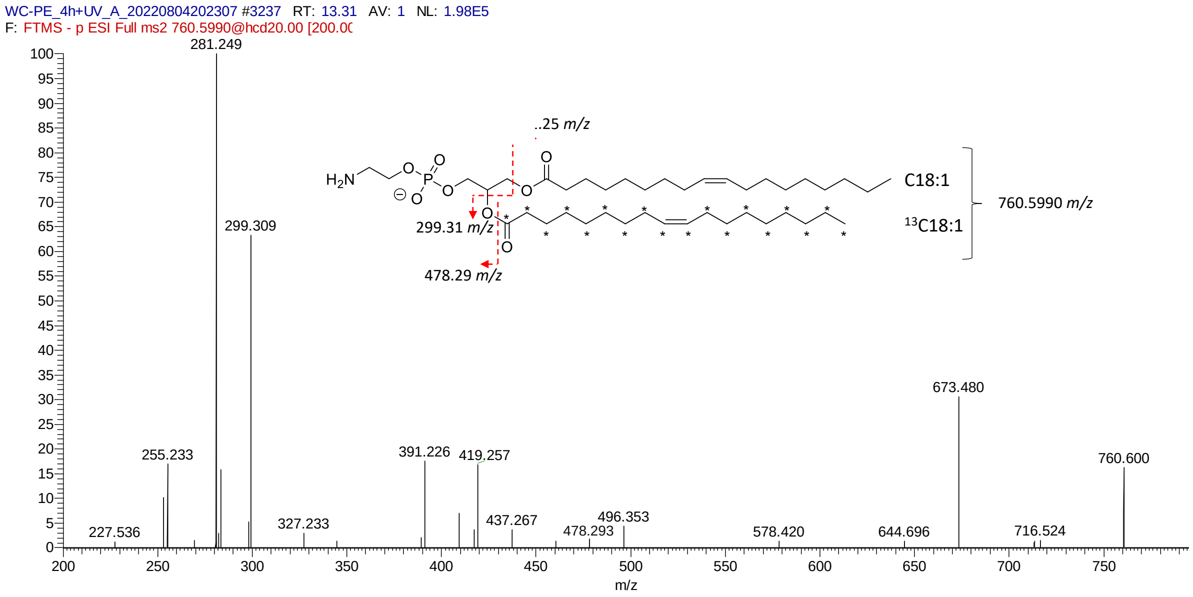

## $^{13}$C-PE34:1

RT = 13.3 min; Negative mode

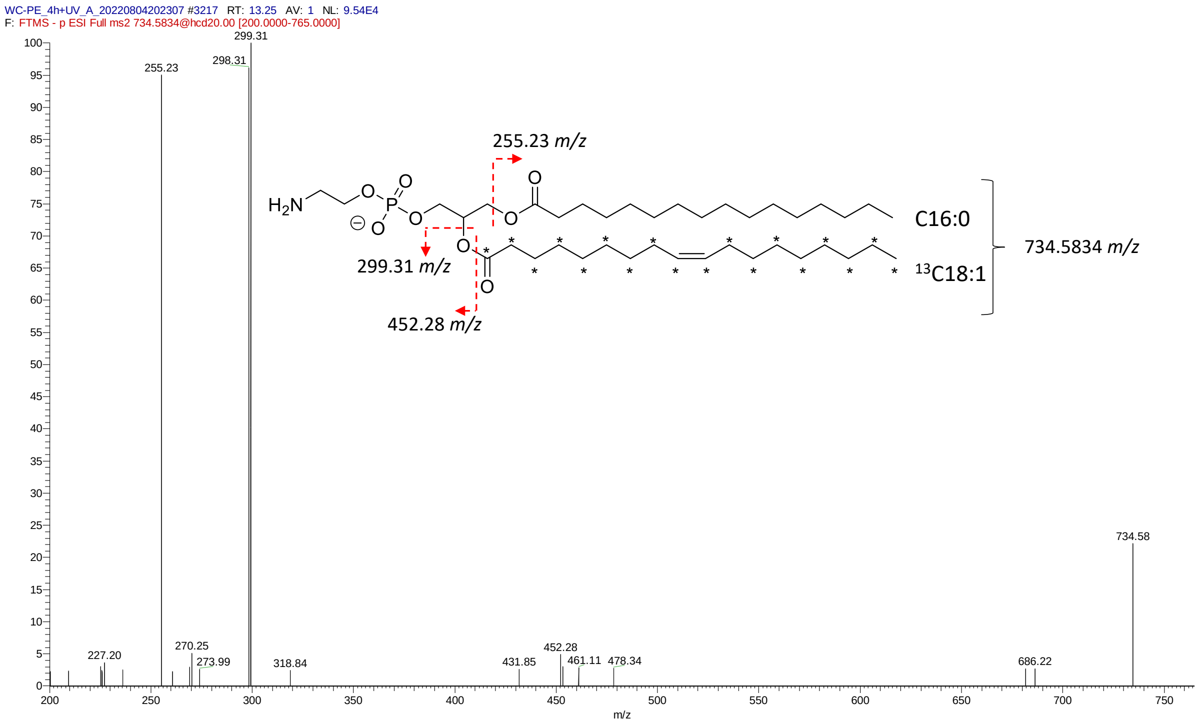

**$^{13}$C-PE36:1**

RT = 14.0 min; Negative mode

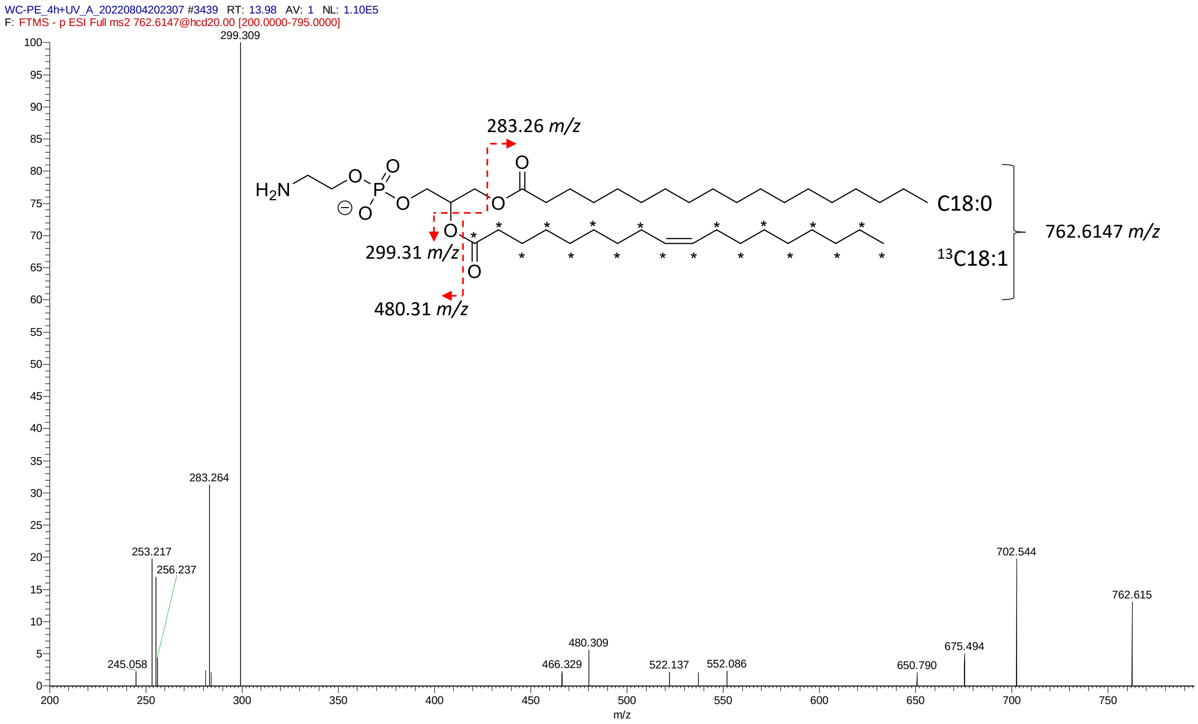

$^{13}$**C-PI species (**ER-PE incorporation in HeLa cells followed by UV uncaging and chase of 4 h).

## $^{13}$C-PI36:2

RT = 12.5 min; Negative mode

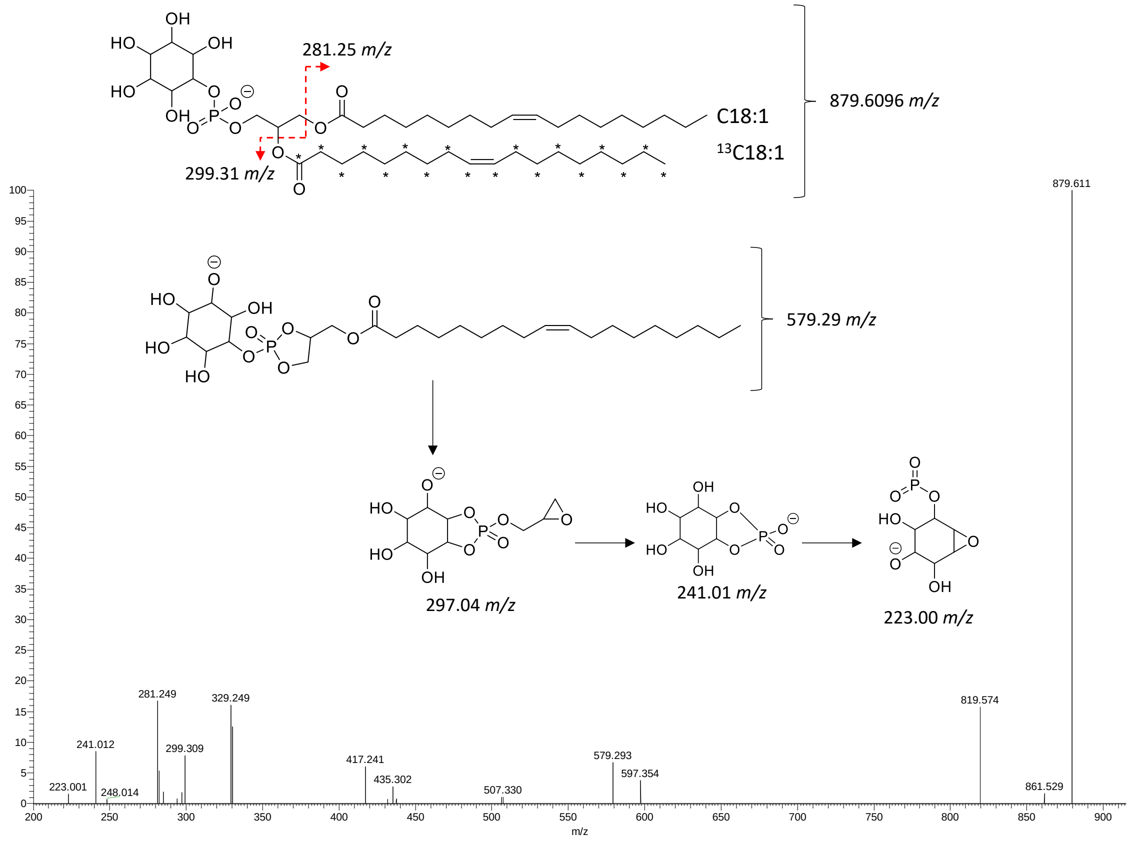

Negative mode, zoom 200-300 *m/z*

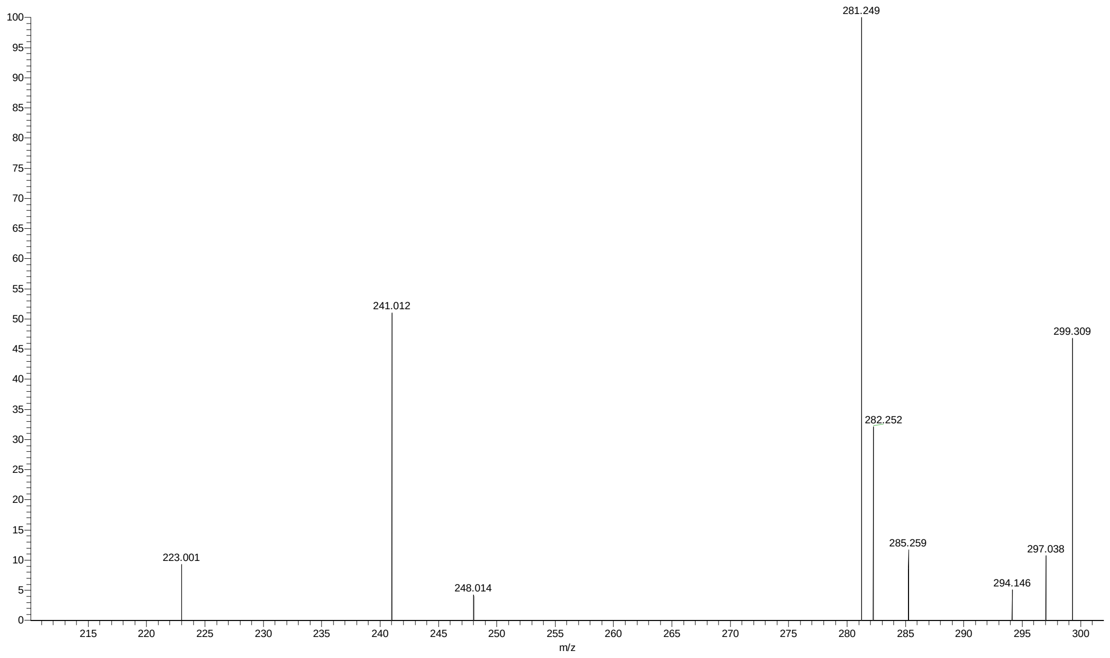

**$^{13}$C-PI36:1**

RT = 13.2 min; Negative mode

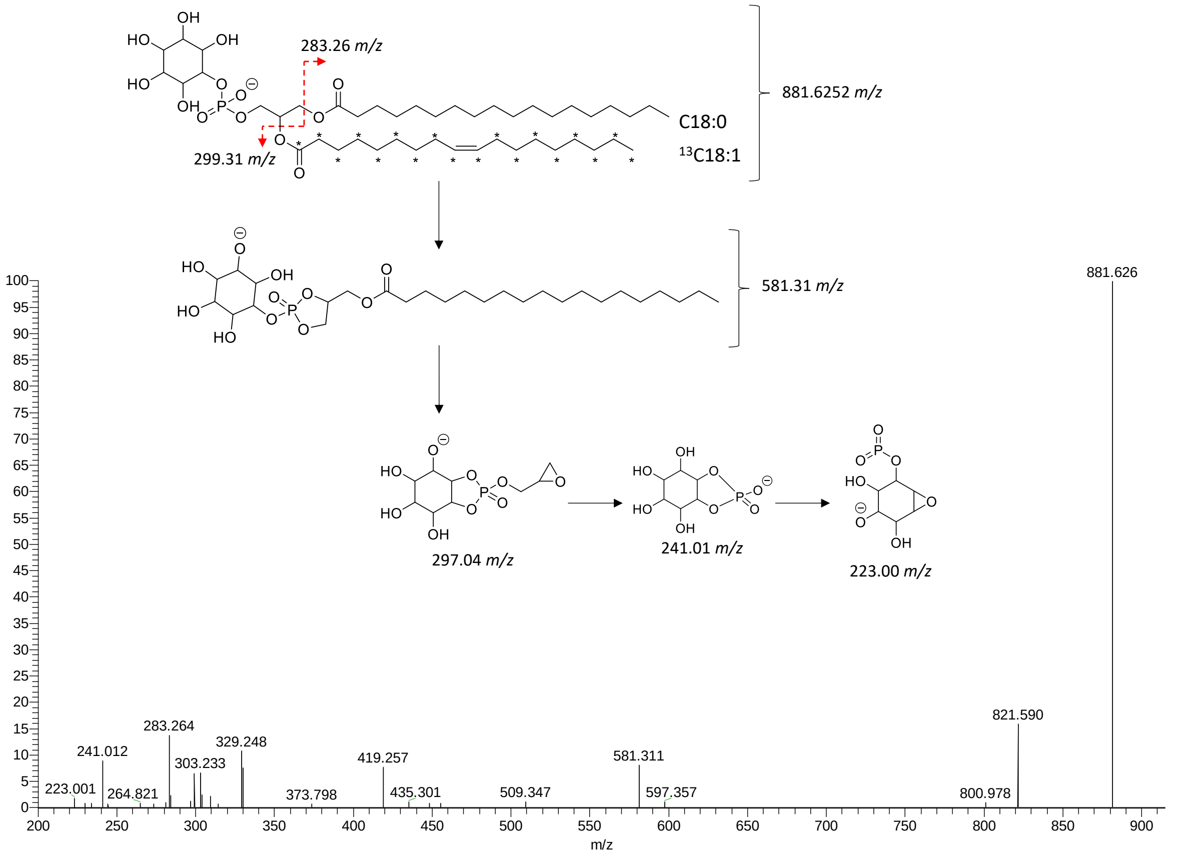

Negative mode, zoom 200-300 *m/z*

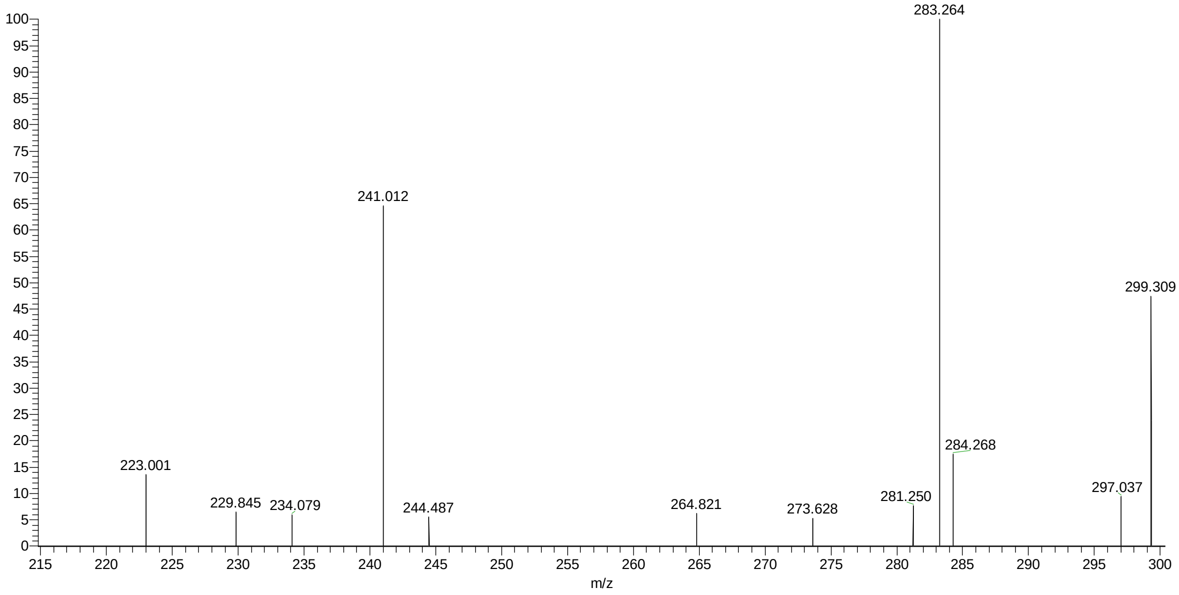

## $^{13}$C-PI34:1

RT = 12.4 min; Negative mode

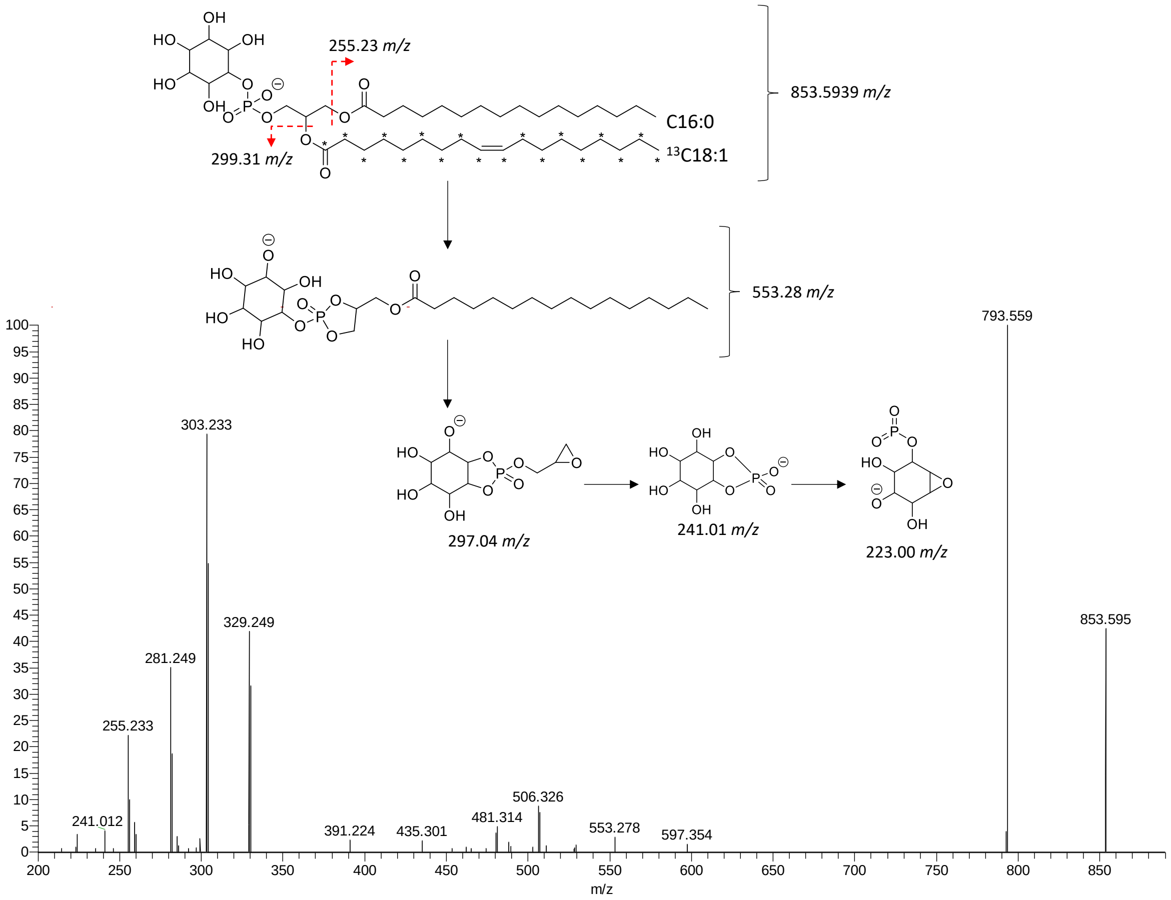

Negative mode, zoom 200-300 *m/z*

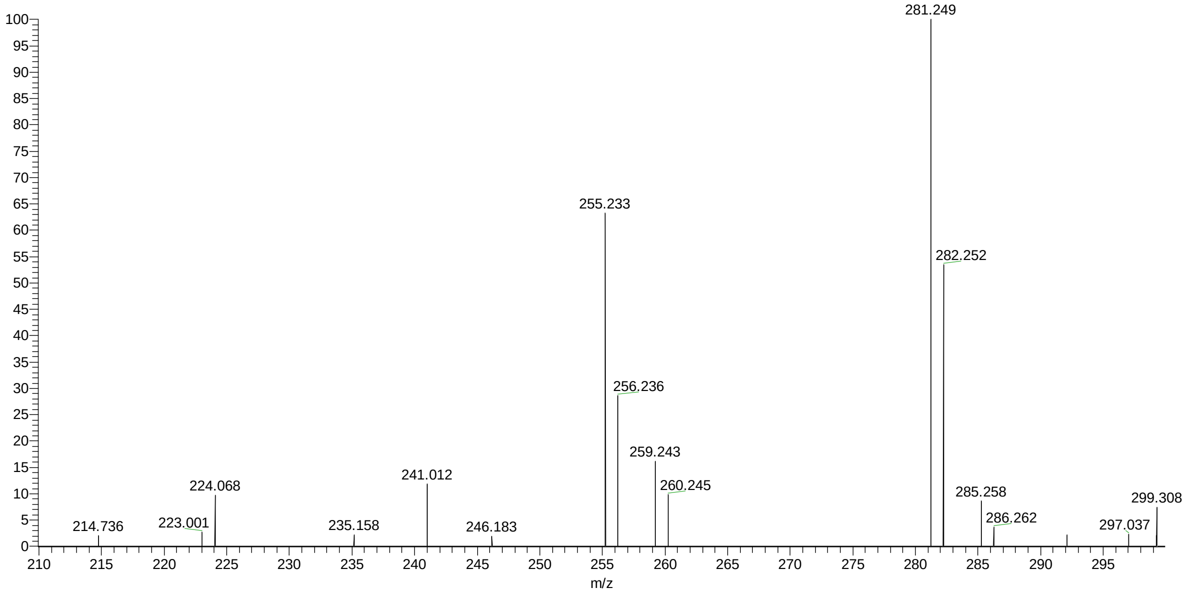

## $^{13}$C-PI34:2

RT = 11.6 min; Negative mode

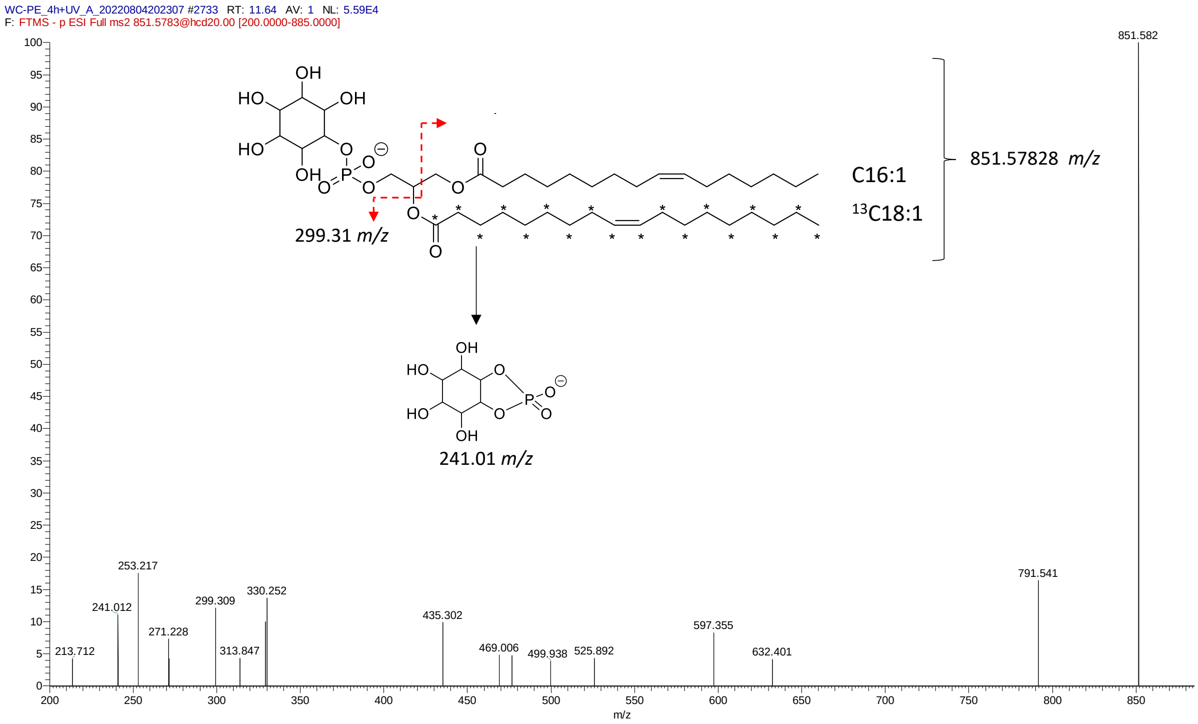

**[13]C-PS species (**ER-PE incorporation in HeLa cells followed by UV uncaging and chase of 16 h).

## [13]C-PS36:1

RT = 13.3 min; Negative mode

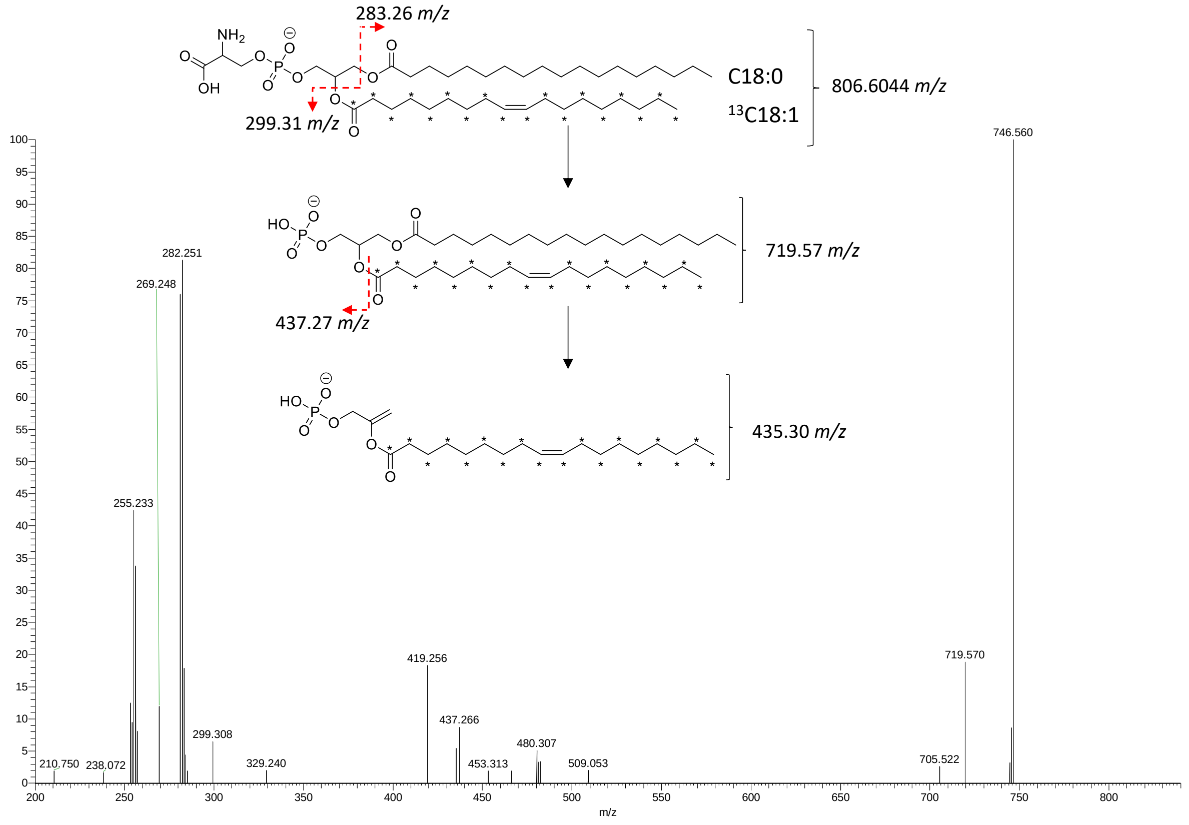

Negative mode, zoom 283-284 *m/z*

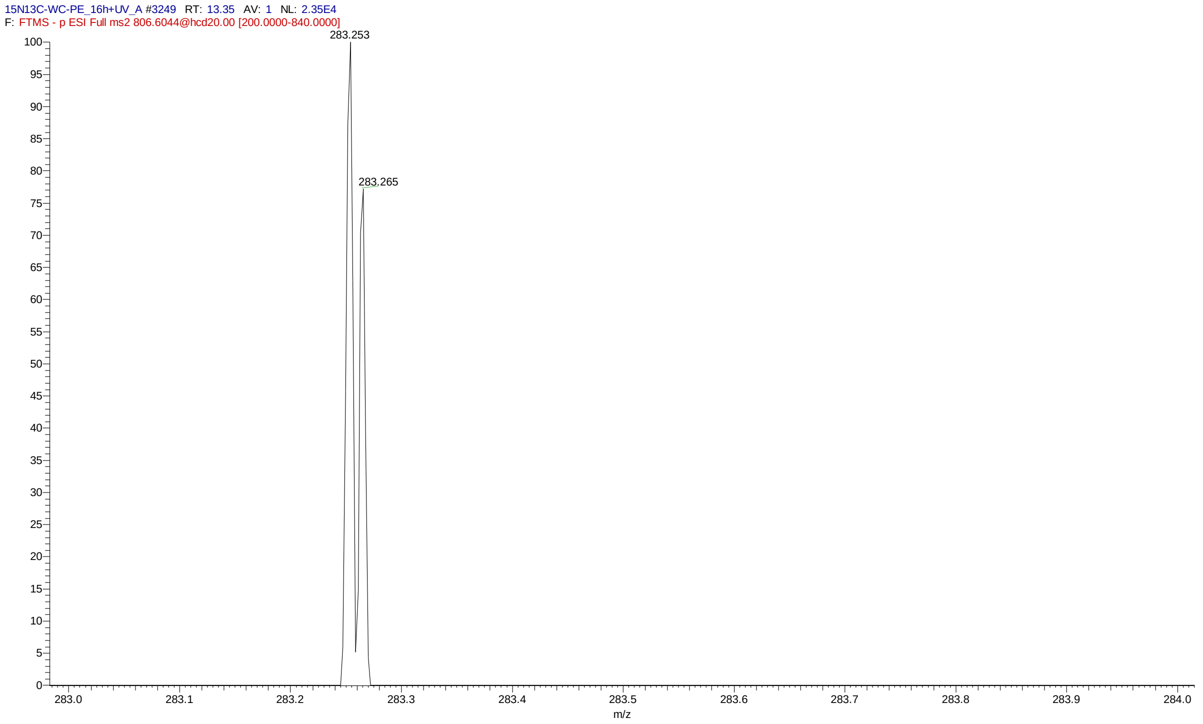

Negative mode, zoom 400-500 *m/z*

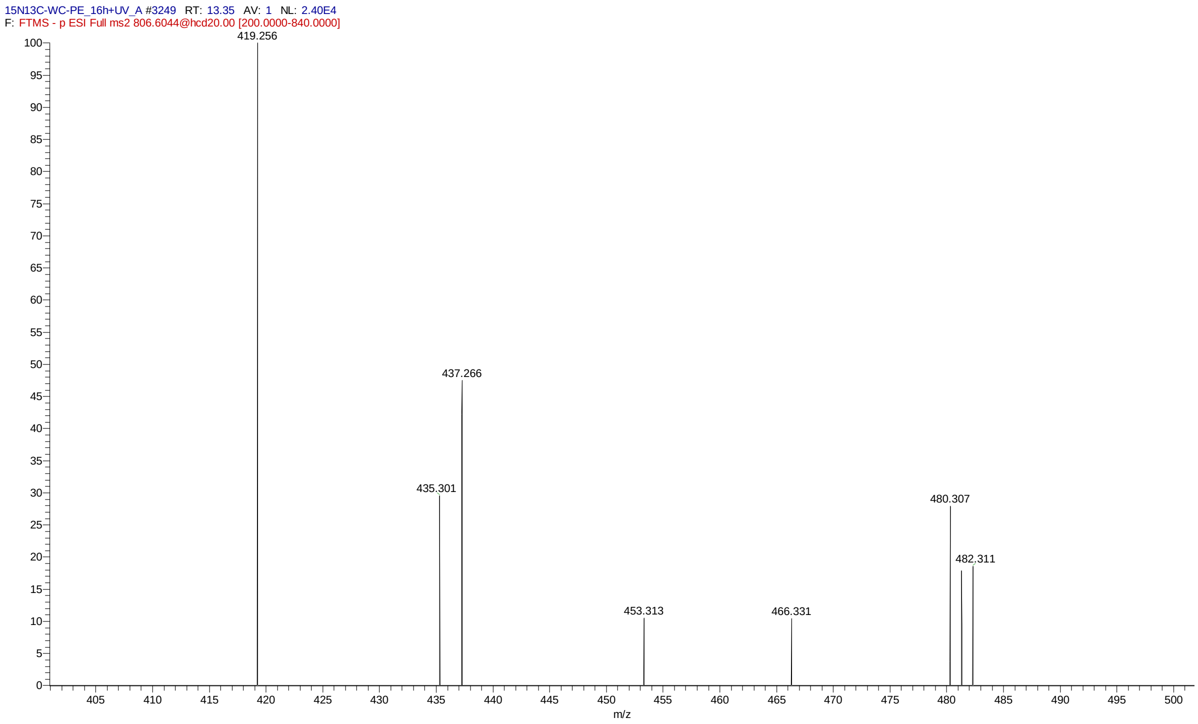

## $^{13}$C-PS36:2

RT = 12.6 min; Negative mode

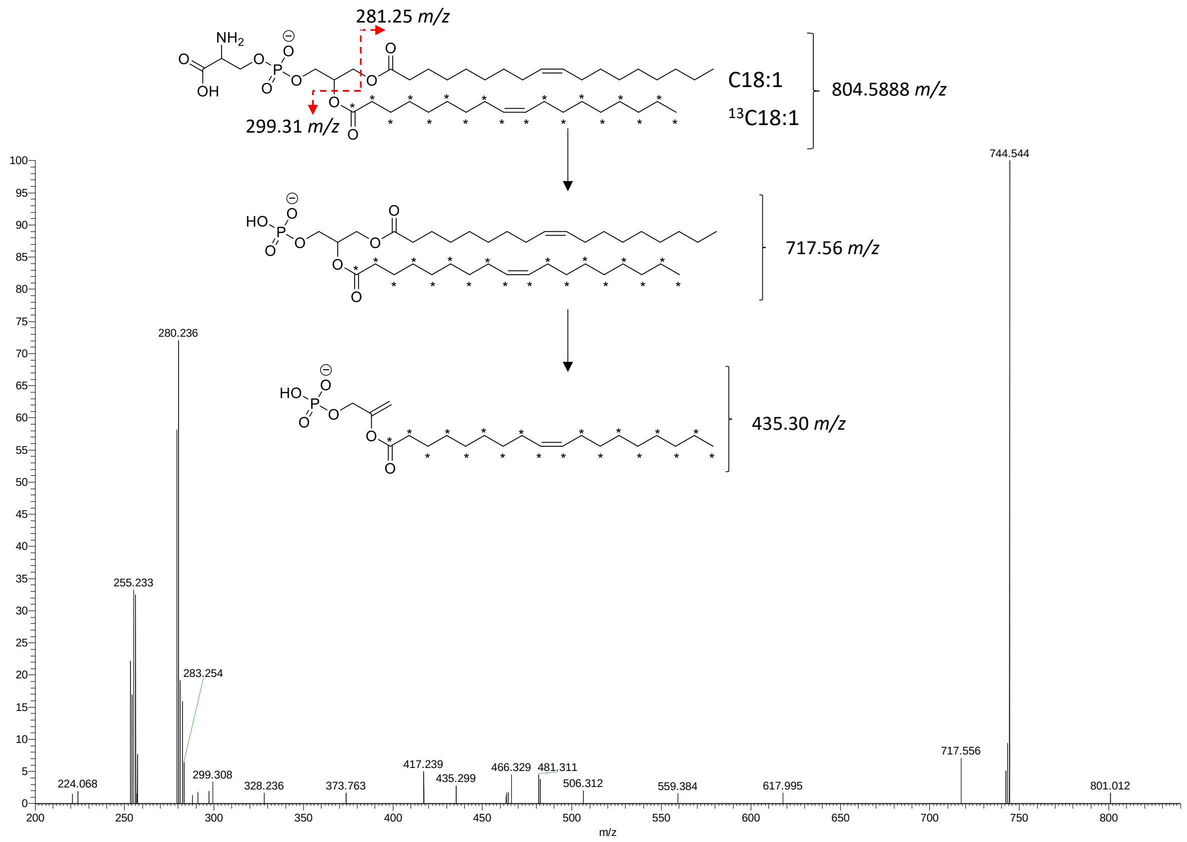

Negative mode, zoom 281-282 *m/z*

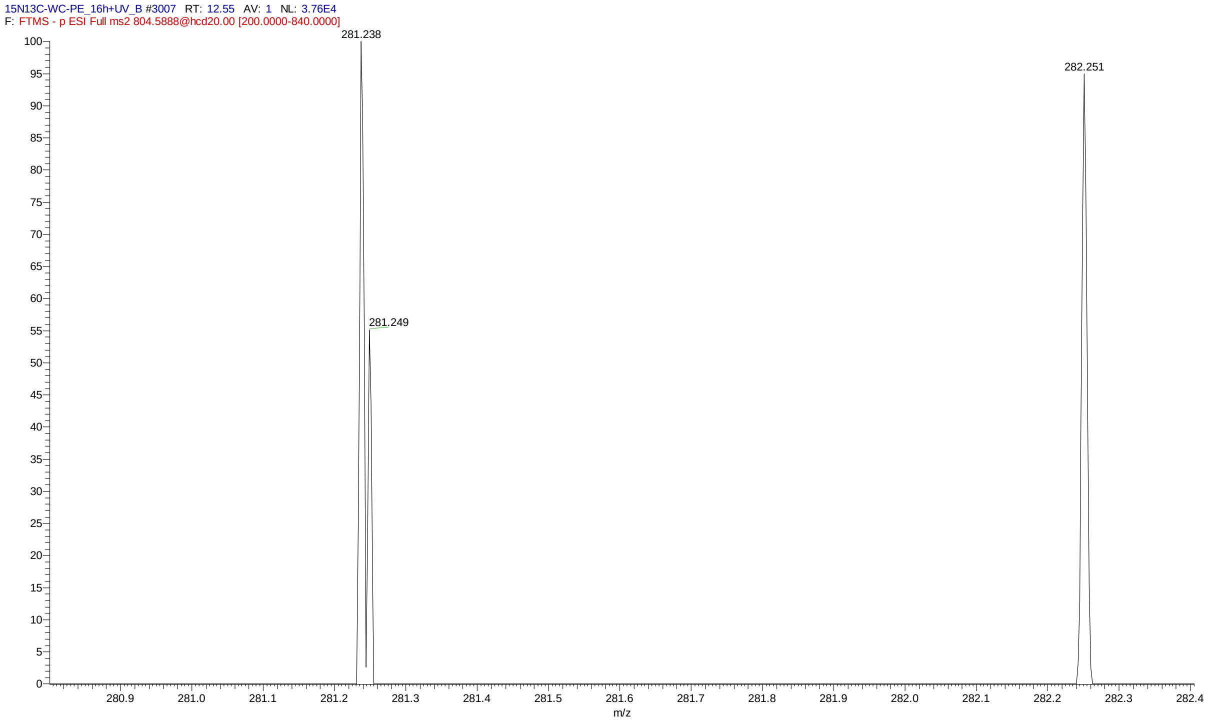

**$^{13}$C-PS34:1**

RT = 12.5 min; Negative mode

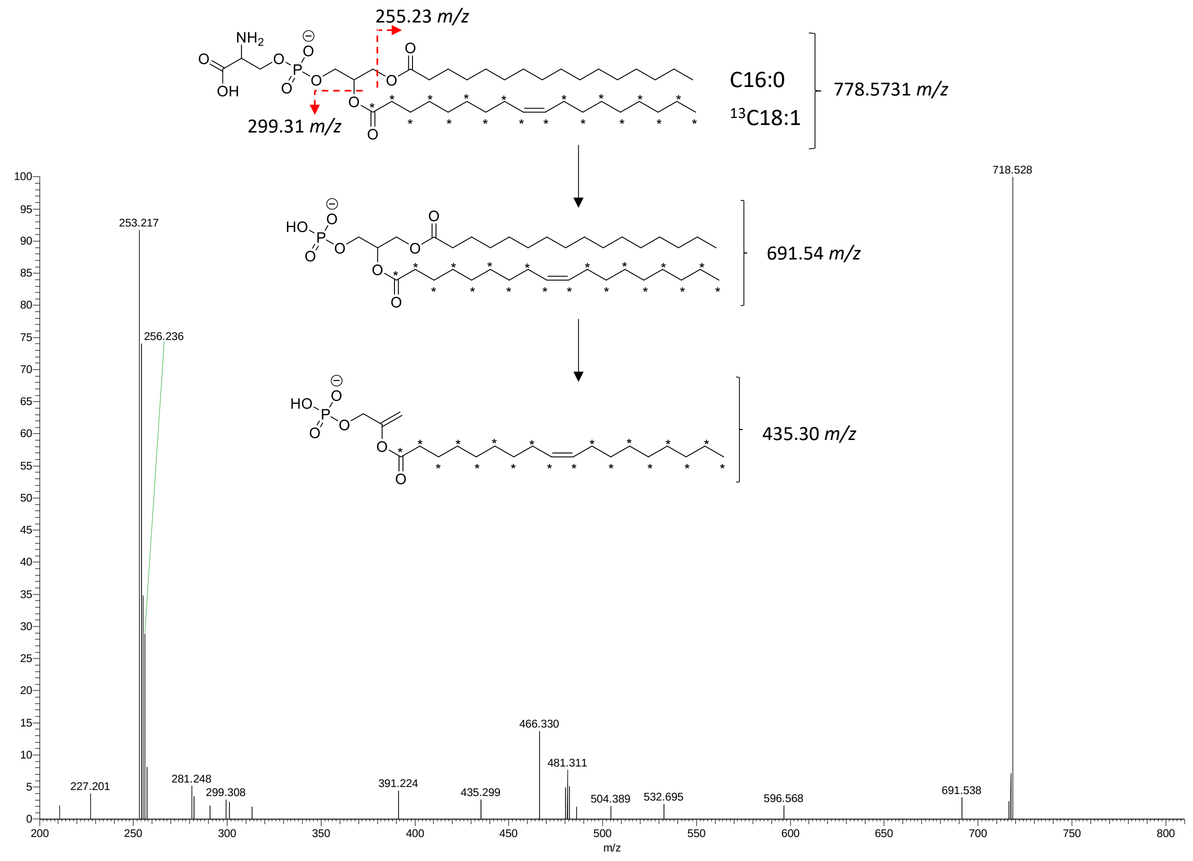

Negative mode, zoom 200-300 *m/z*

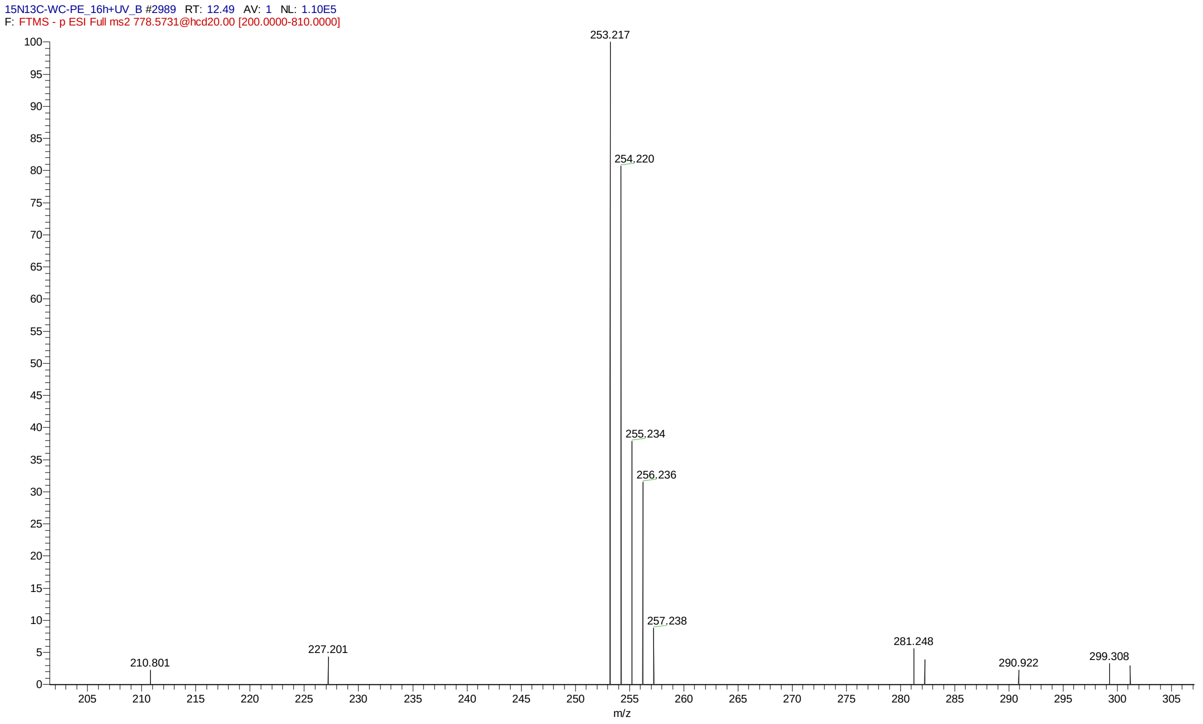

$^{15}N^{13}C$**-PE species** (ER-PE incorporation in HeLa cells followed by UV uncaging and chase of 4 h)

## $^{15}N\ ^{13}C$-PE36:1 = $^{15}N$-PE-18:0-$^{13}C18:1$

RT = 14.0 min; Positive mode

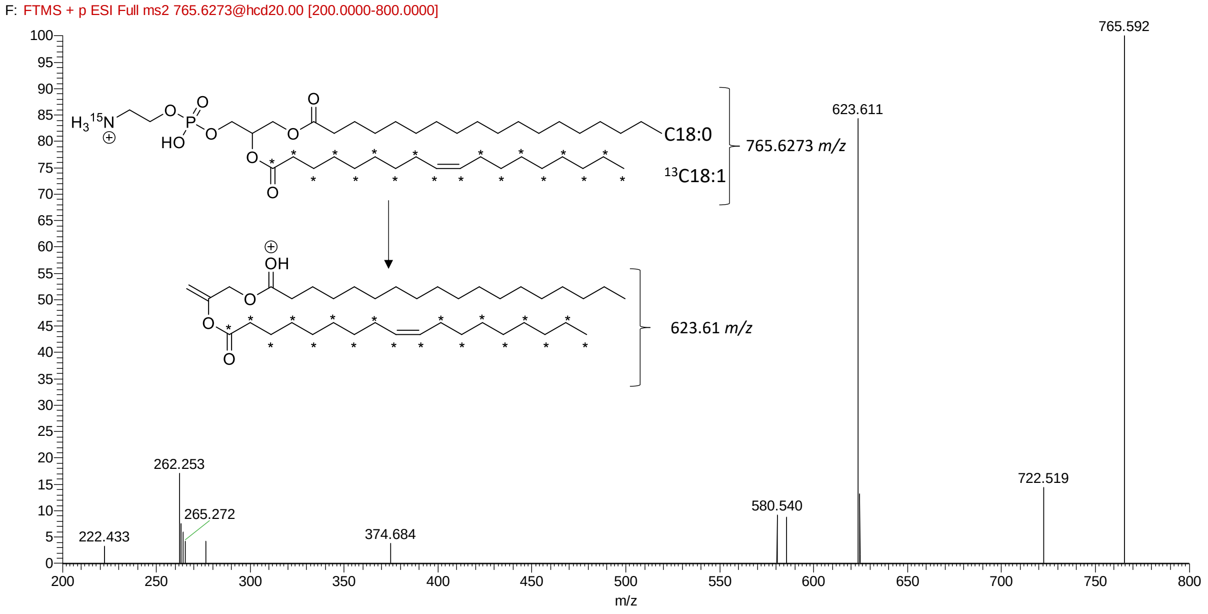

Negative mode

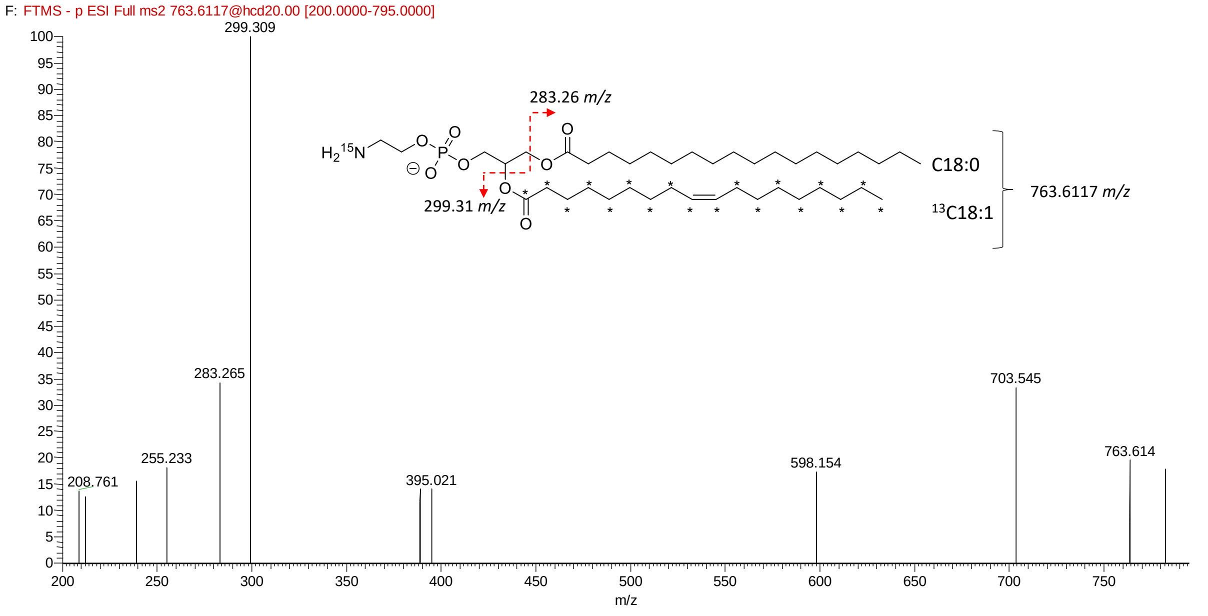

**$^{15}$N $^{13}$C-PE36:2 = $^{15}$N-PE-18:1-$^{13}$C18:1**

RT = 13.3 min; Positive mode

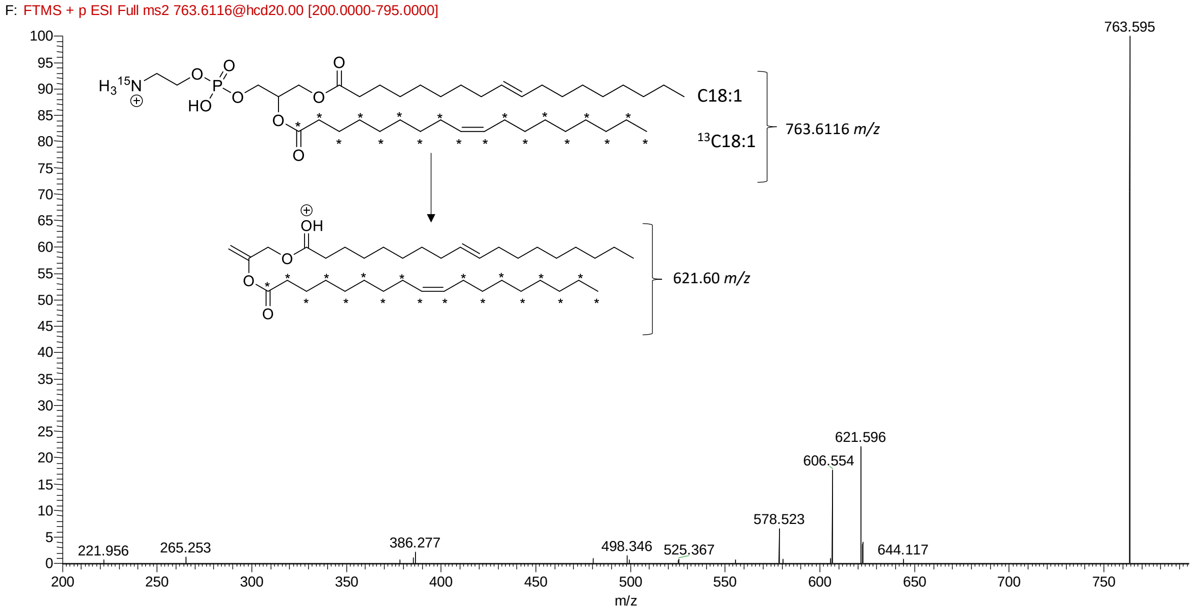

Negative mode

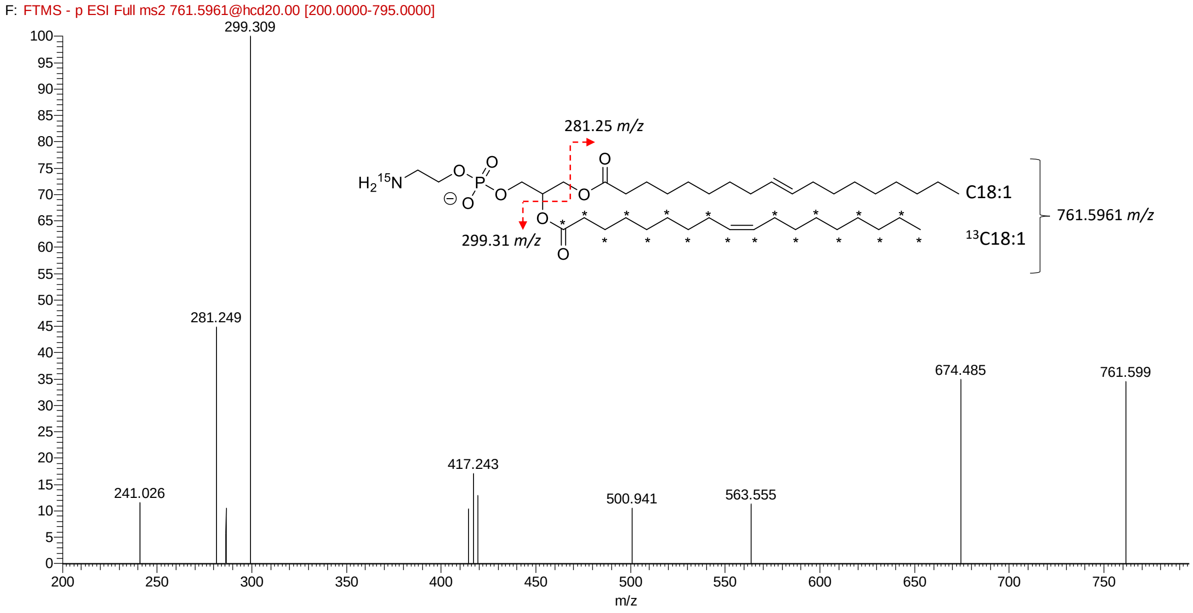

**$^{15}N^{13}C$-PC34:1 = $^{15}N$-PC-16:0-$^{13}C18:1$** (ER-PE incorporation in co-transfected HeLa cells (with siRNA targeting *CHPT1* and *CEPT1* and with PEMT2 plasmid) followed by UV uncaging and chase of 4 h

RT = 13.0 min; Negative mode

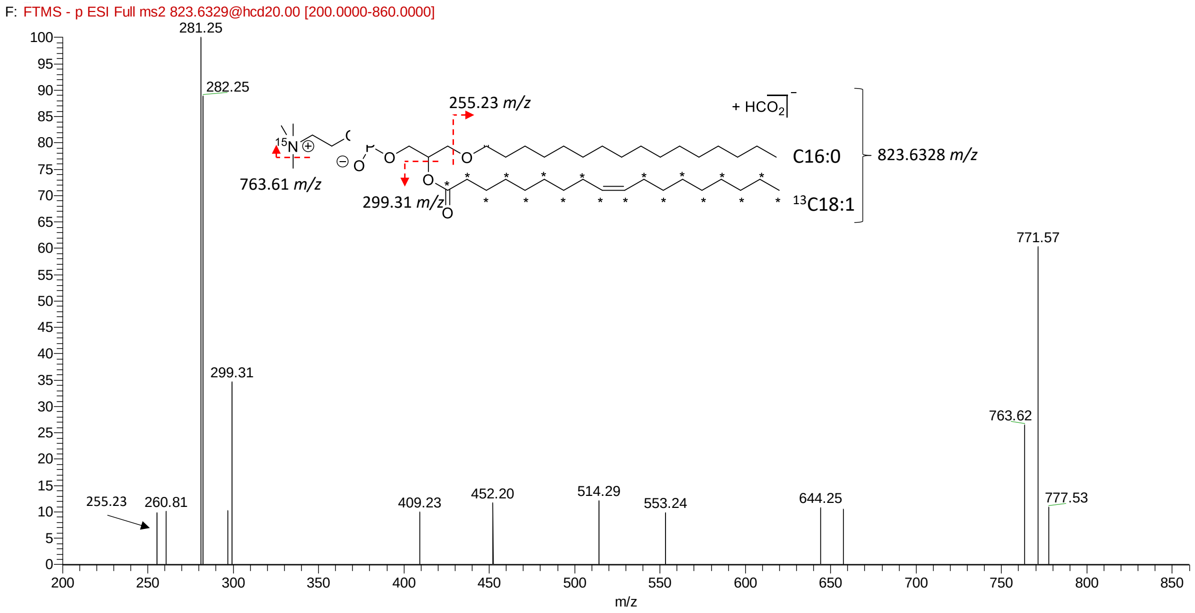

### Spectroscopic data of key compounds (NMR, HR-MS)

1. $^{15}N^{13}C$-PE34:1

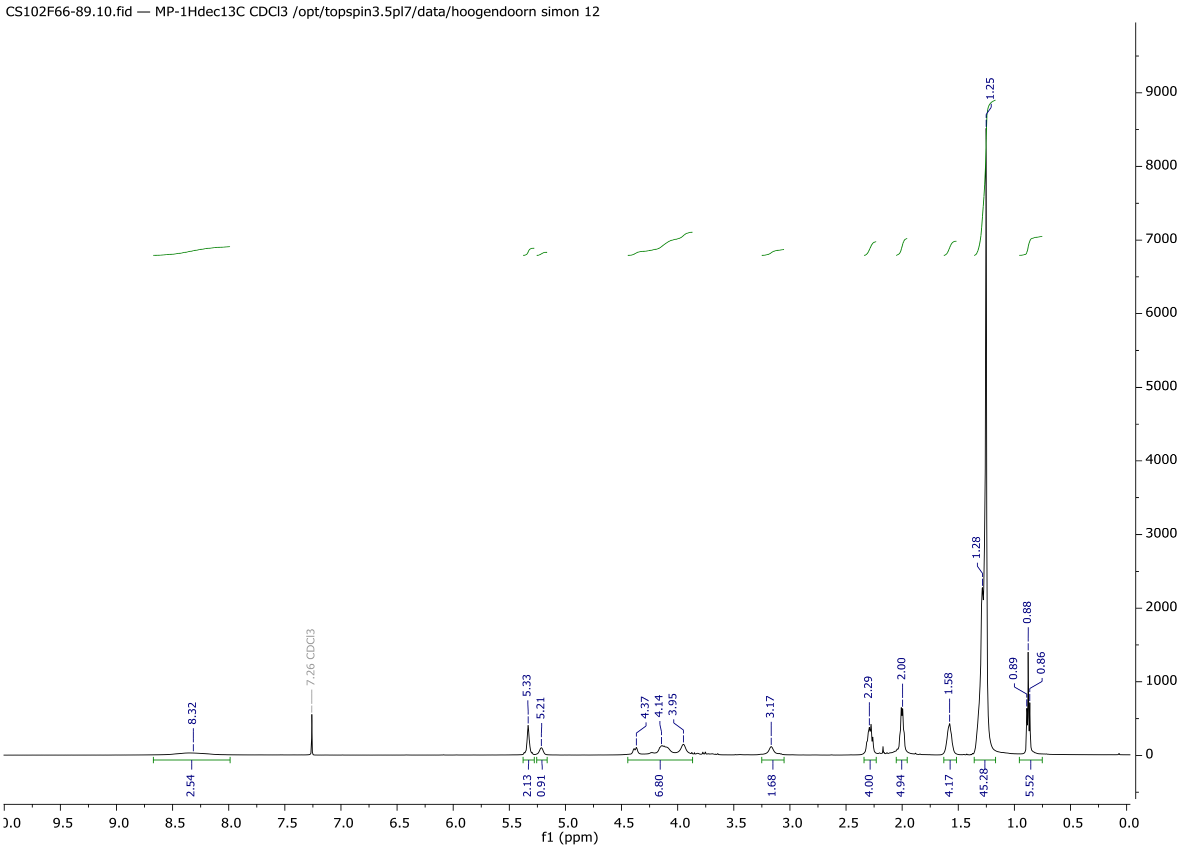

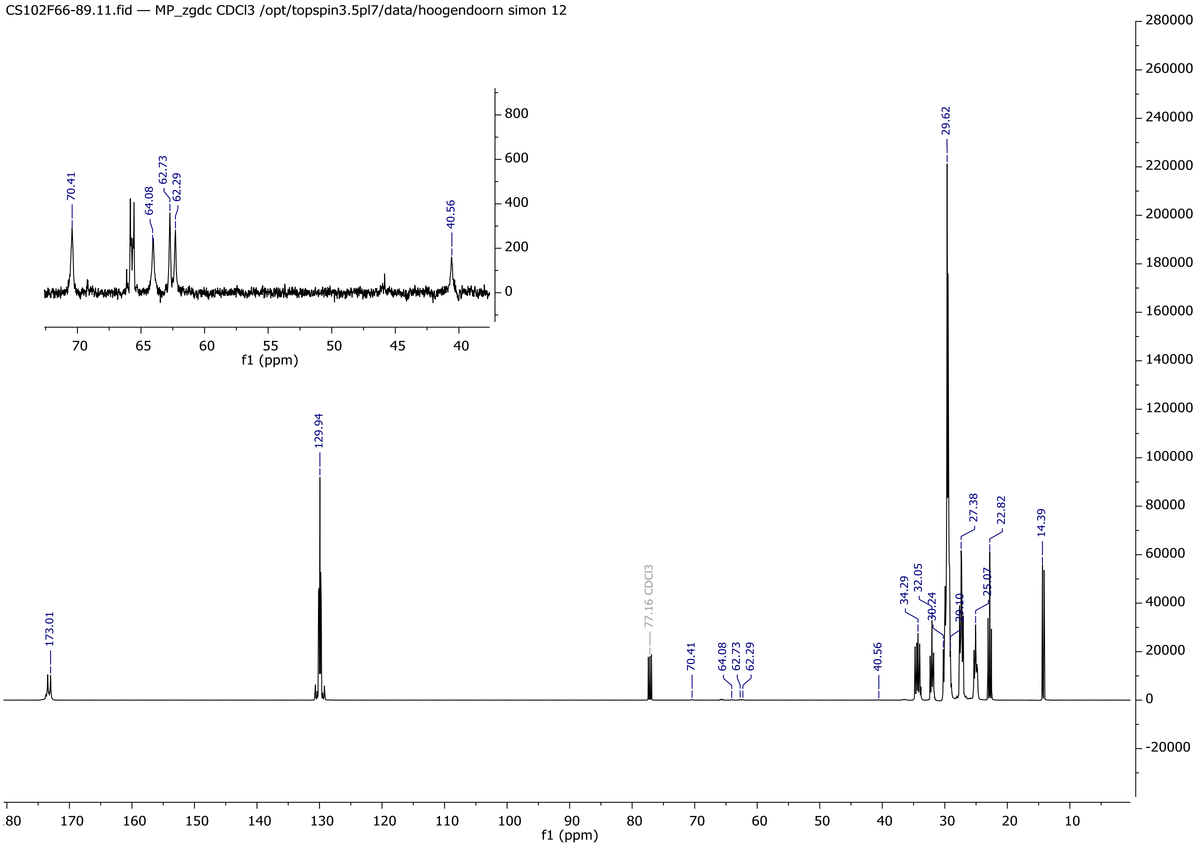

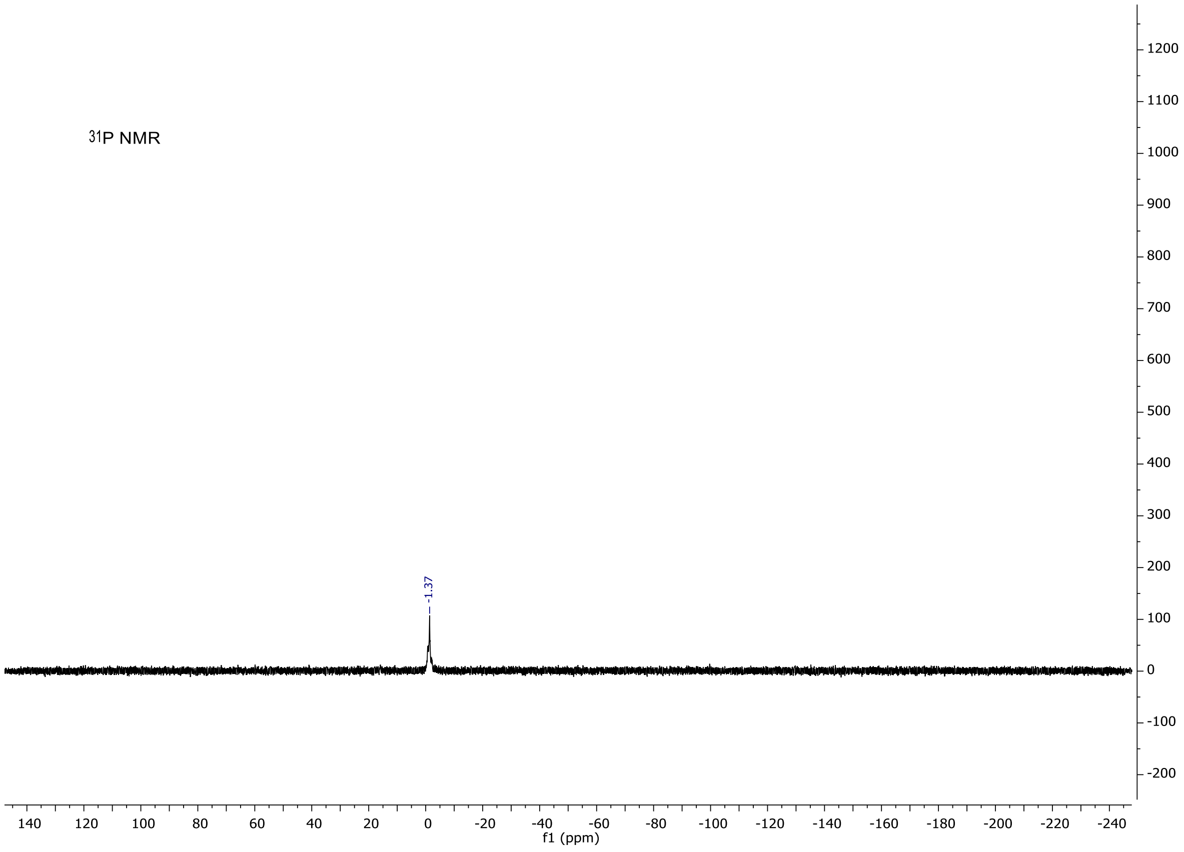

31P NMR
-1.37
f1 (ppm)

HR-MS

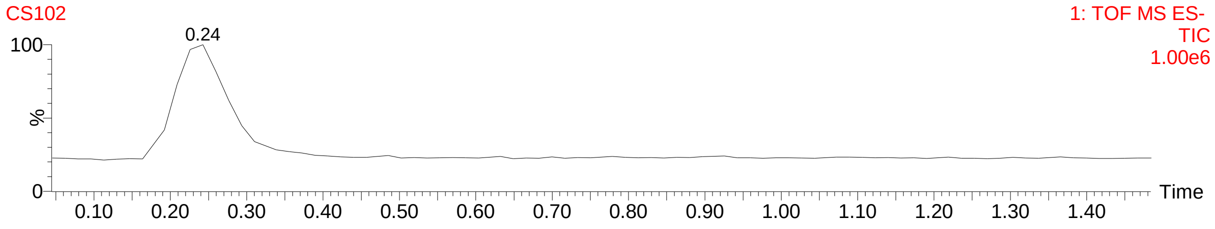

CS102
1: TOF MS ES-
TIC
1.00e6
0.24
%
Time

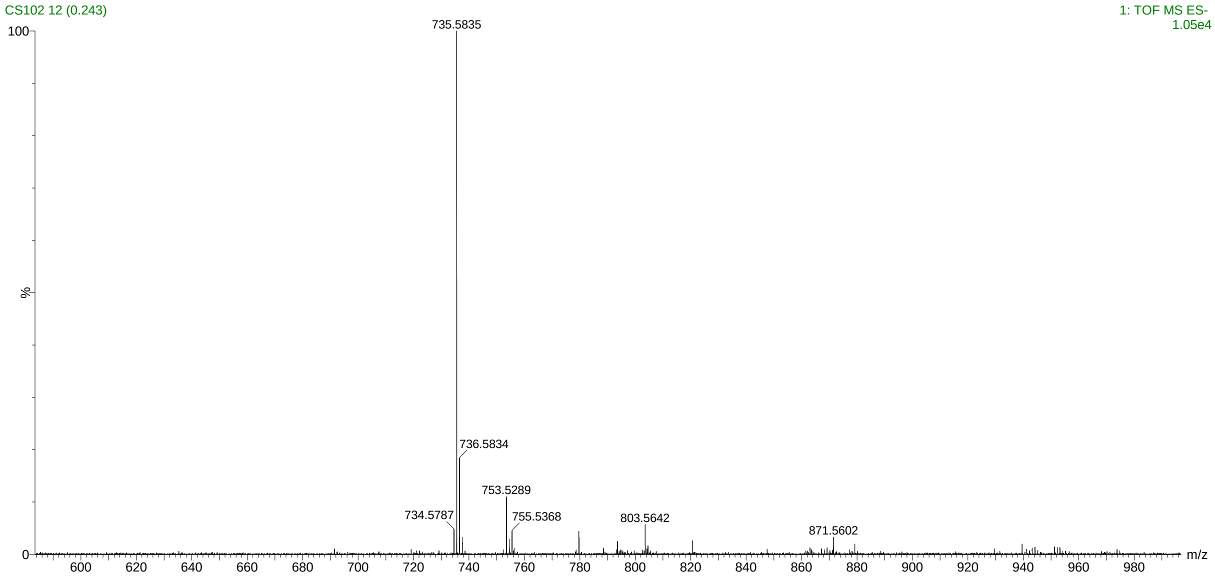

CS102 12 (0.243)
1: TOF MS ES-
1.05e4
735.5835
736.5834
753.5289
734.5787
755.5368
803.5642
871.5602
%
m/z

#### 2. MitoPE

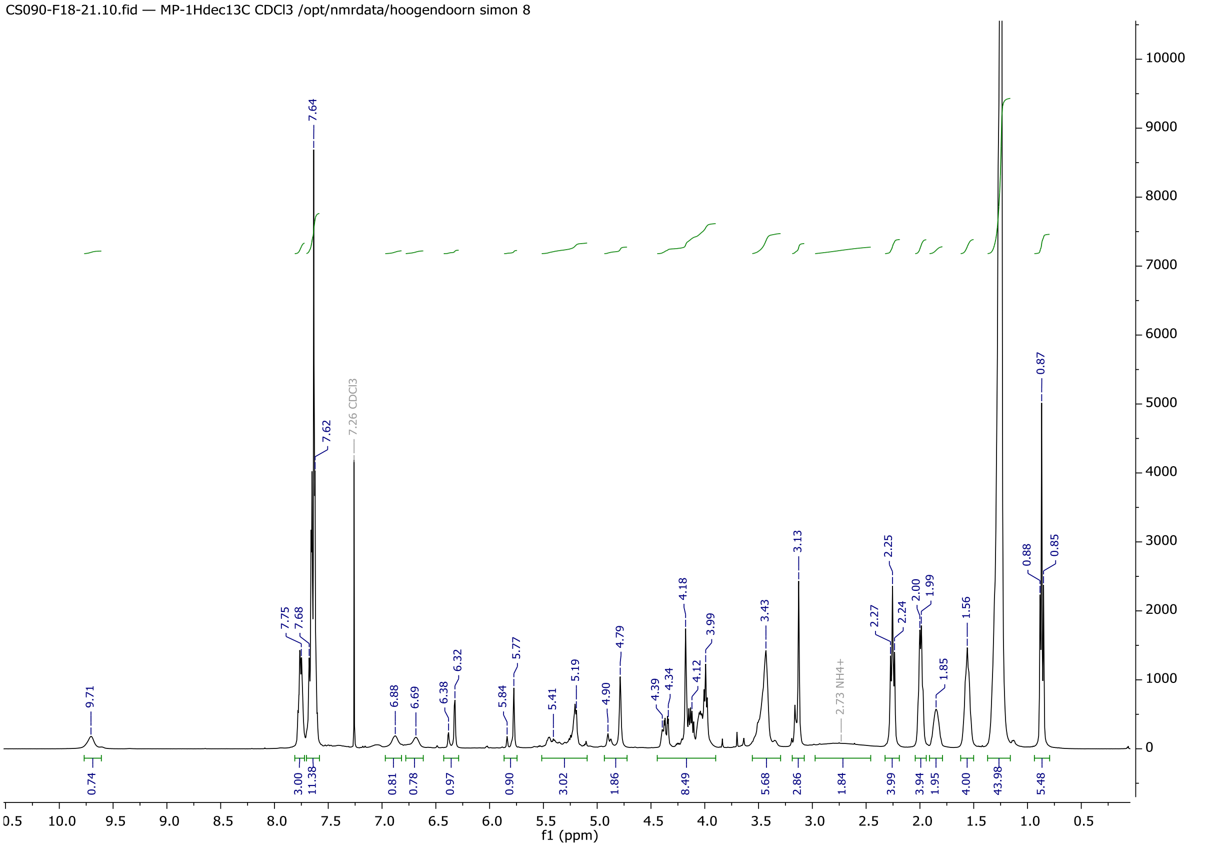

CS090-F18-21.10.fid — MP-1Hdec13C CDCl3 /opt/nmrdata/hoogendoorn simon 8
f1 (ppm)

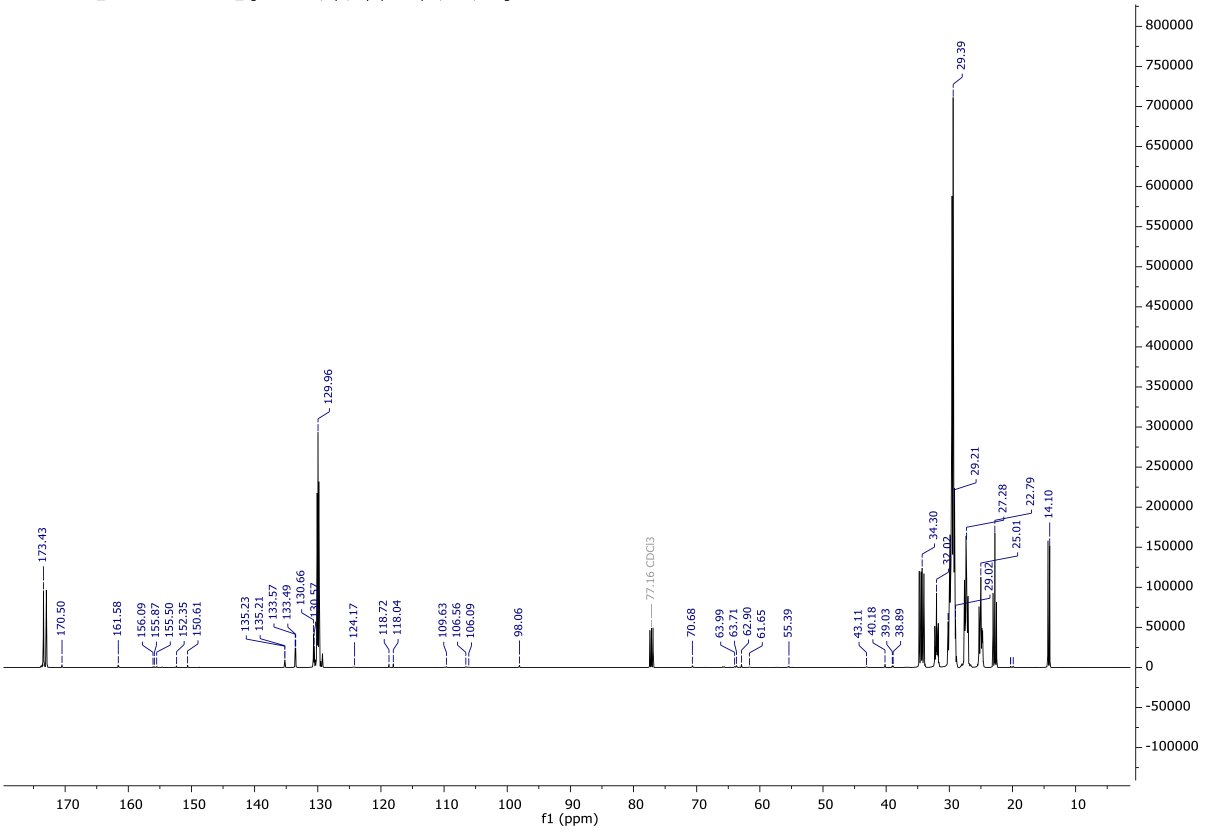

CS090-F19-21_211117.11.fid — MP_zgdc CDCl3 /opt/topspin3.5pl7/data/hoogendoorn simon 14
f1 (ppm)

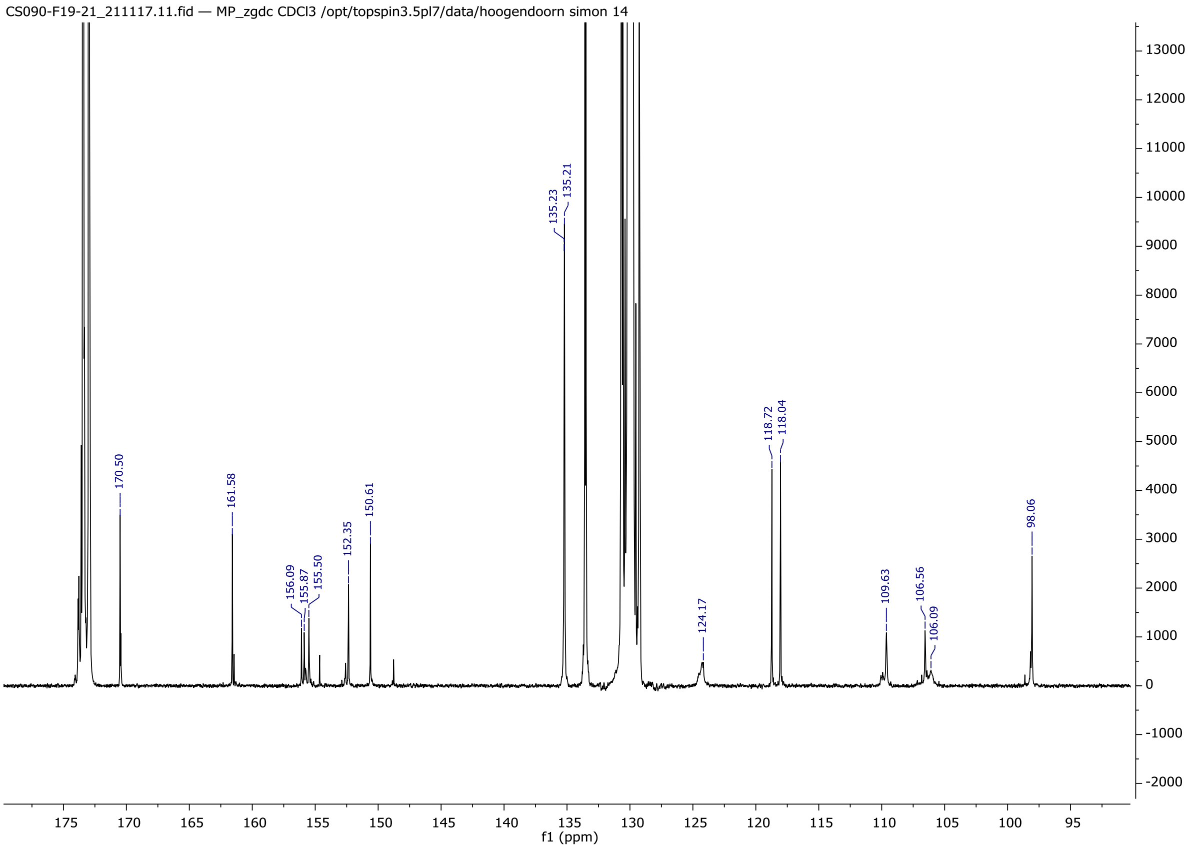
CS090-F19-21_211117.11.fid — MP_zgdc CDCl3 /opt/topspin3.5pl7/data/hoogendoorn simon 14
170.50
161.58
156.09
155.87
155.50
152.35
150.61
135.23
135.21
124.17
118.72
118.04
109.63
106.56
106.09
98.06
f1 (ppm)

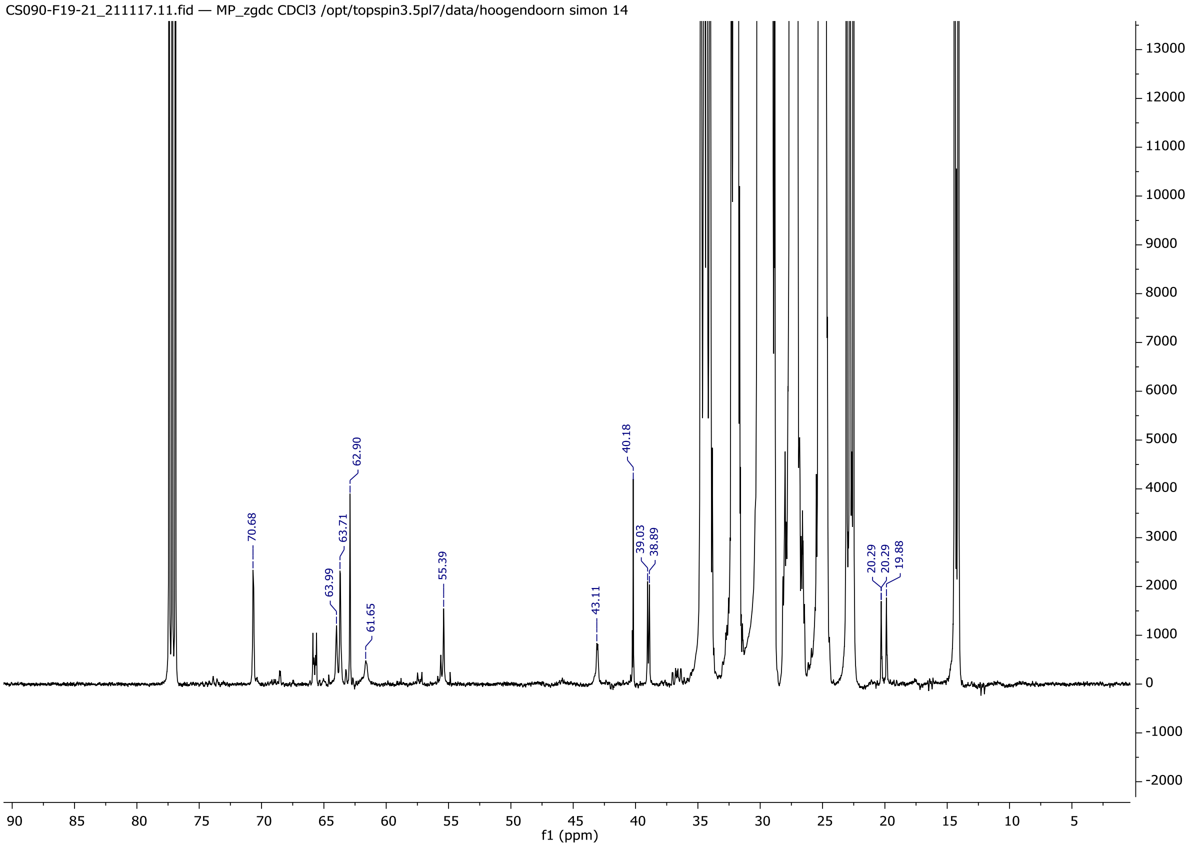
CS090-F19-21_211117.11.fid — MP_zgdc CDCl3 /opt/topspin3.5pl7/data/hoogendoorn simon 14
70.68
63.99
63.71
62.90
61.65
55.39
43.11
40.18
39.03
38.89
20.29
20.29
19.88
f1 (ppm)

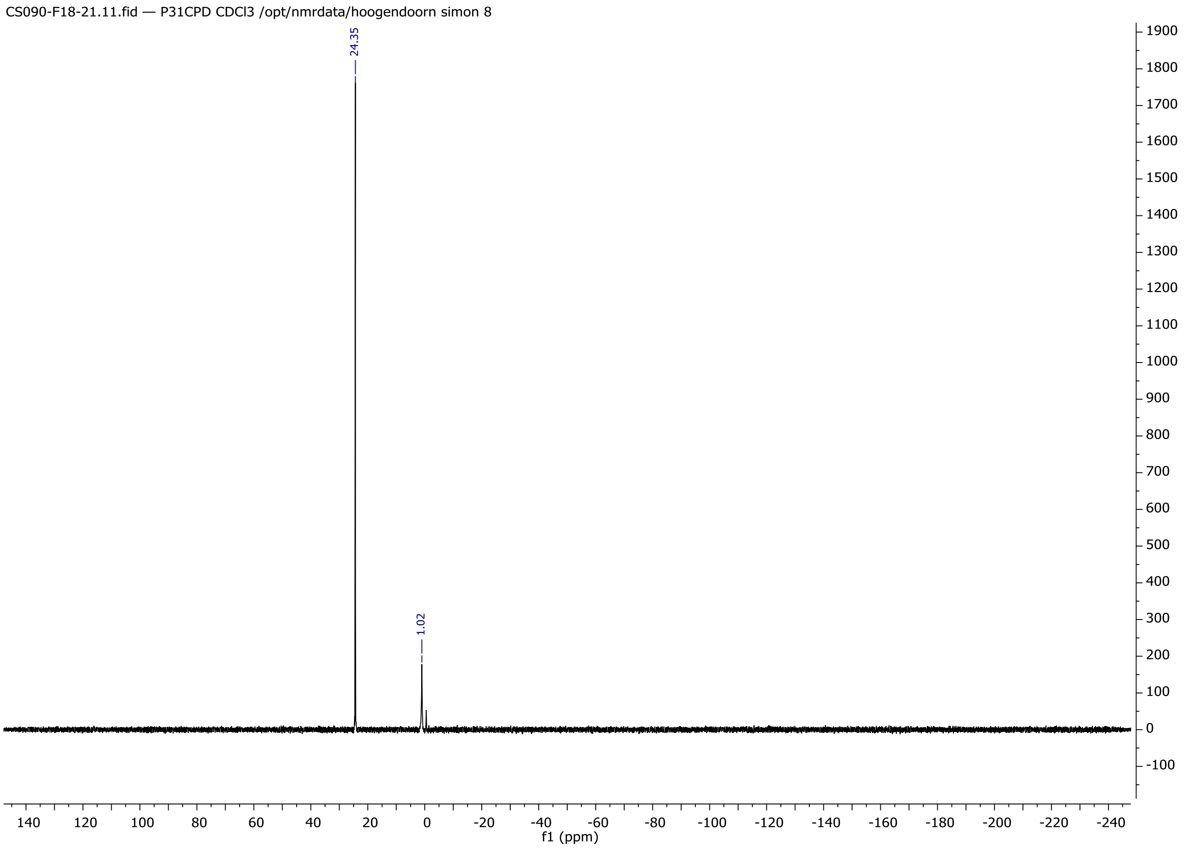
CS090-F18-21.11.fid — P31CPD CDCl3 /opt/nmrdata/hoogendoorn simon 8
24.35
1.02
f1 (ppm)

HR-MS

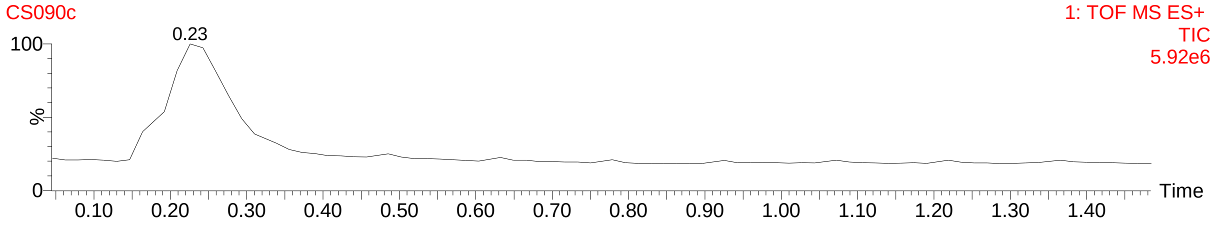
CS090c
1: TOF MS ES+
TIC
5.92e6
0.23
%
Time

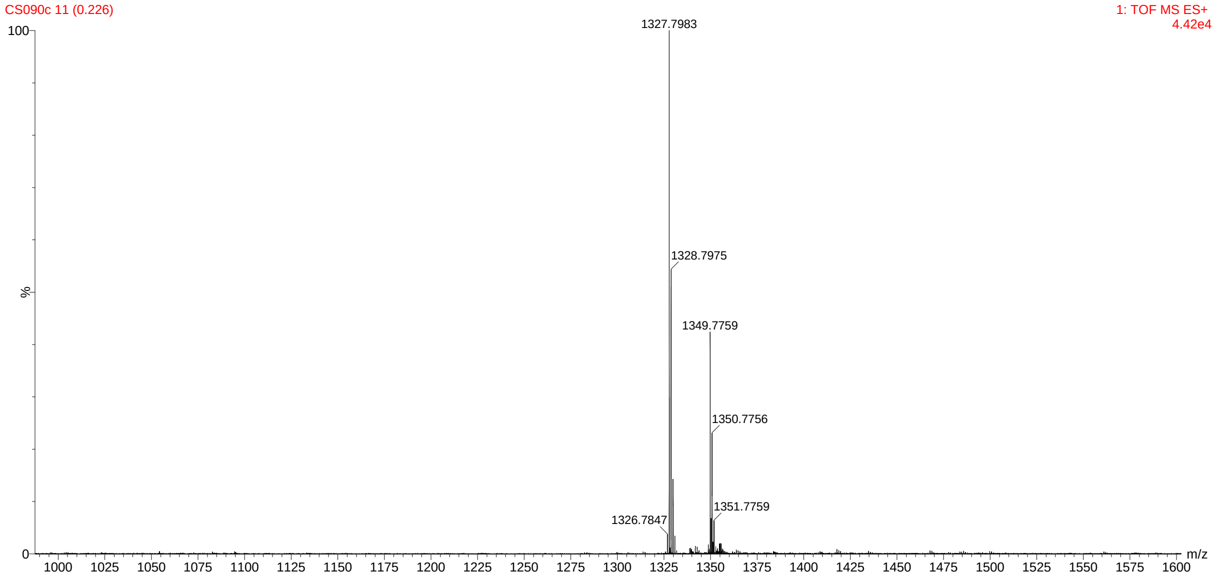
CS090c 11 (0.226)
1: TOF MS ES+
4.42e4
1327.7983
1328.7975
1349.7759
1350.7756
1351.7759
1326.7847
%
m/z

## 3. ER-PE

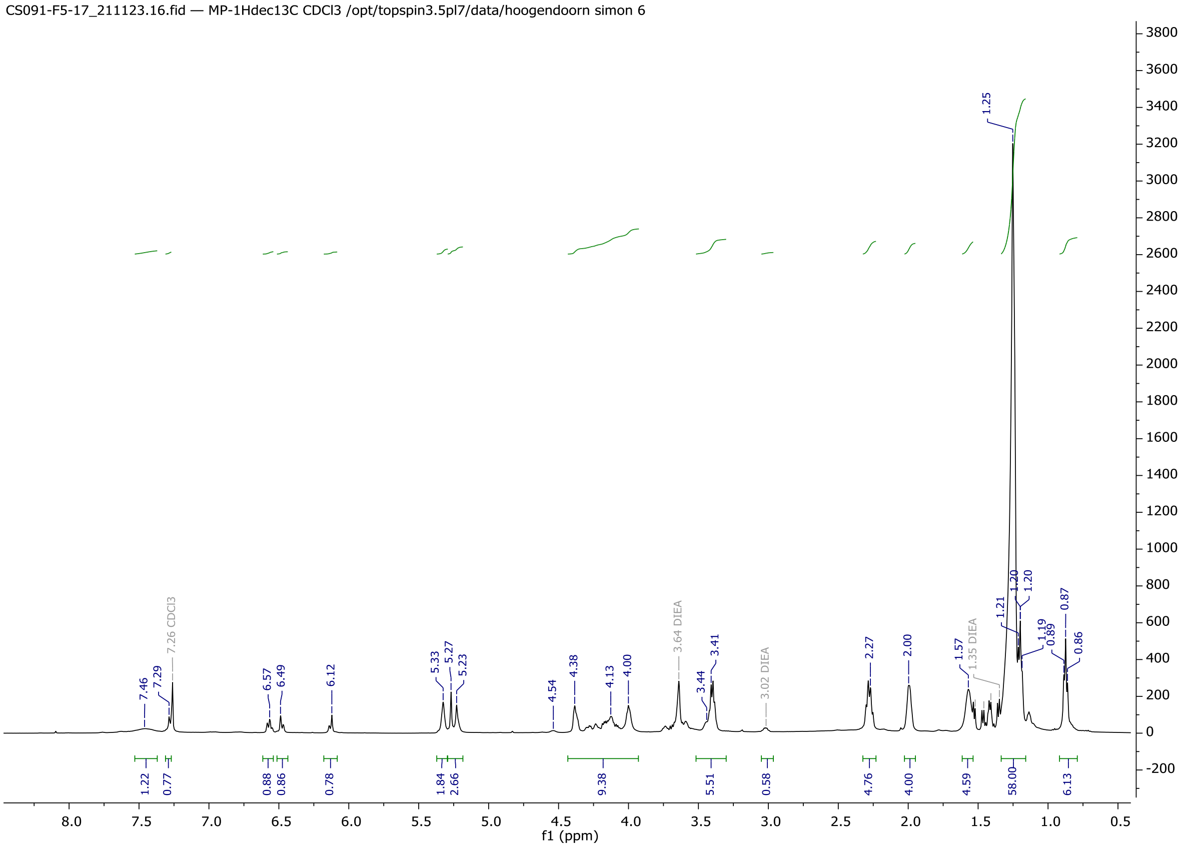
CS091-F5-17_211123.16.fid — MP-1Hdec13C CDCl3 /opt/topspin3.5pl7/data/hoogendoorn simon 6
f1 (ppm)

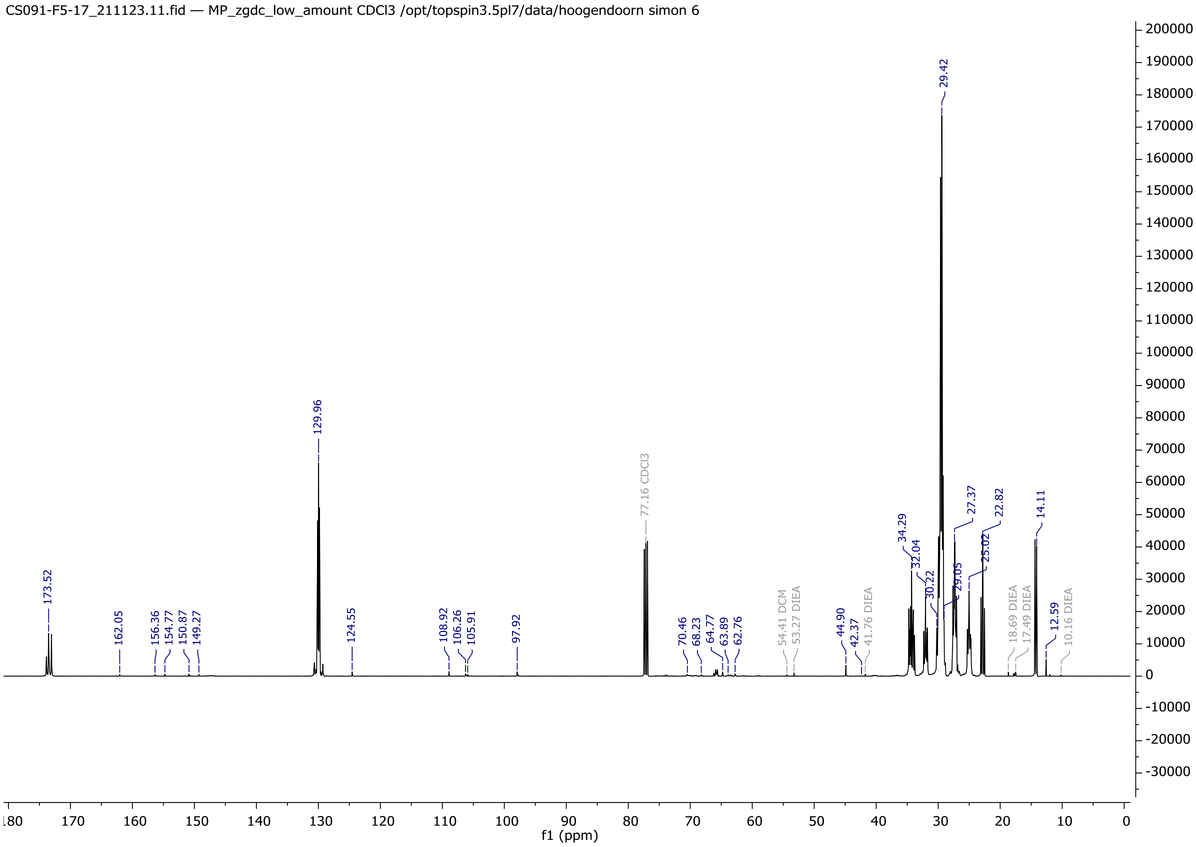
CS091-F5-17_211123.11.fid — MP_zgdc_low_amount CDCl3 /opt/topspin3.5pl7/data/hoogendoorn simon 6
f1 (ppm)

CS091-F5-17_211123.11.fid — MP_zgdc_low_amount CDCl3 /opt/topspin3.5pl7/data/hoogendoorn simon 6

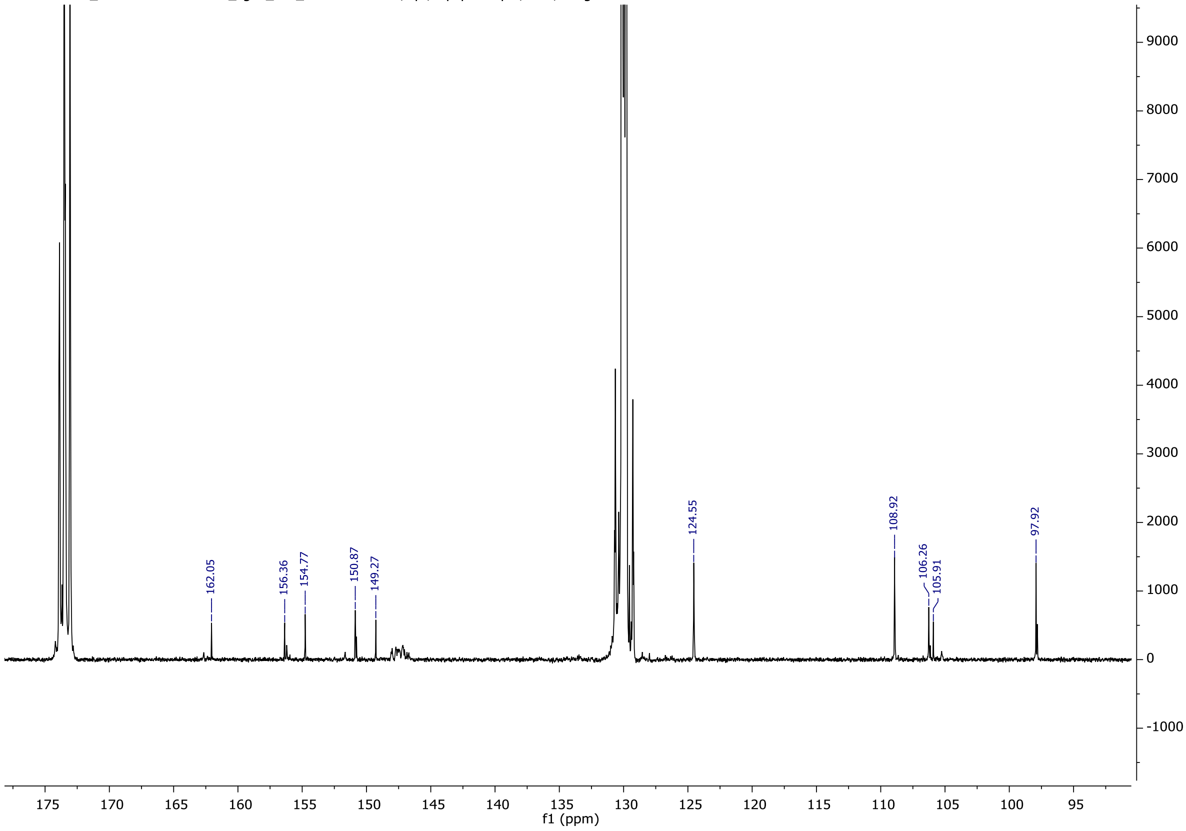

CS091-F5-17_211123.11.fid — MP_zgdc_low_amount CDCl3 /opt/topspin3.5pl7/data/hoogendoorn simon 6

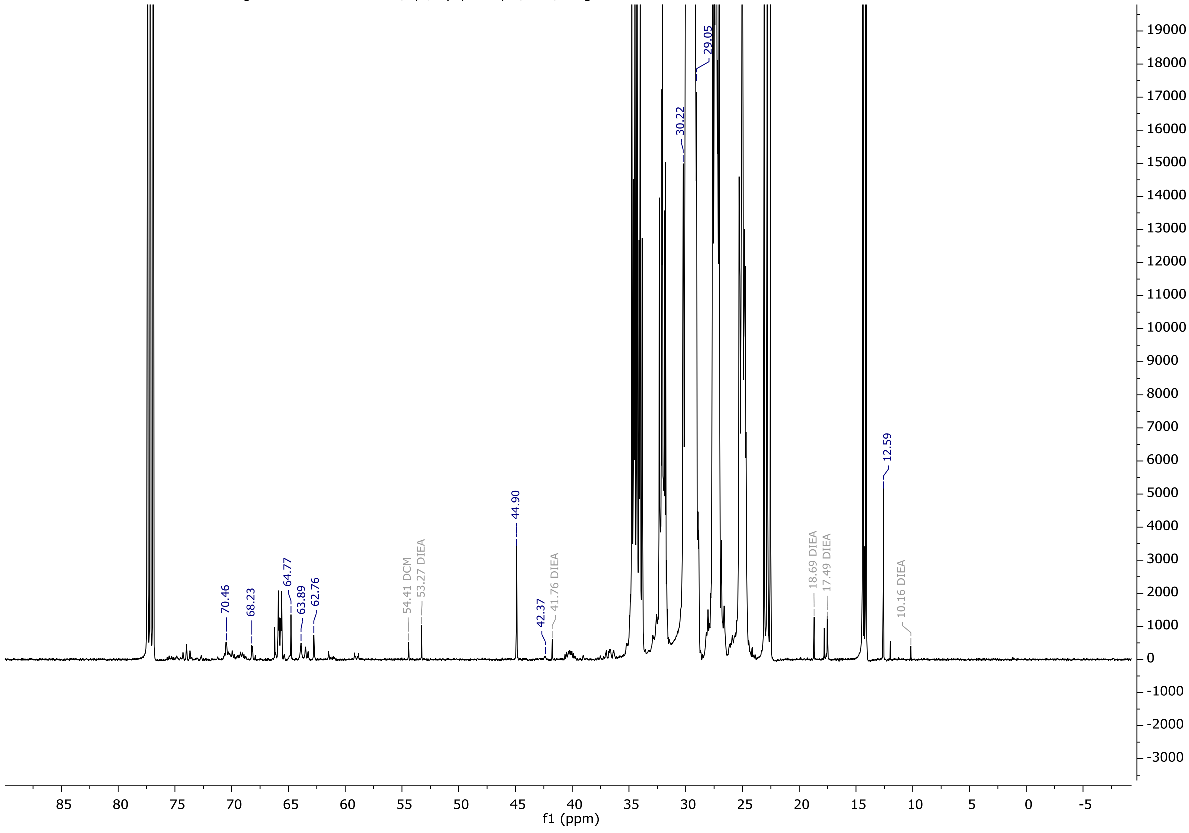

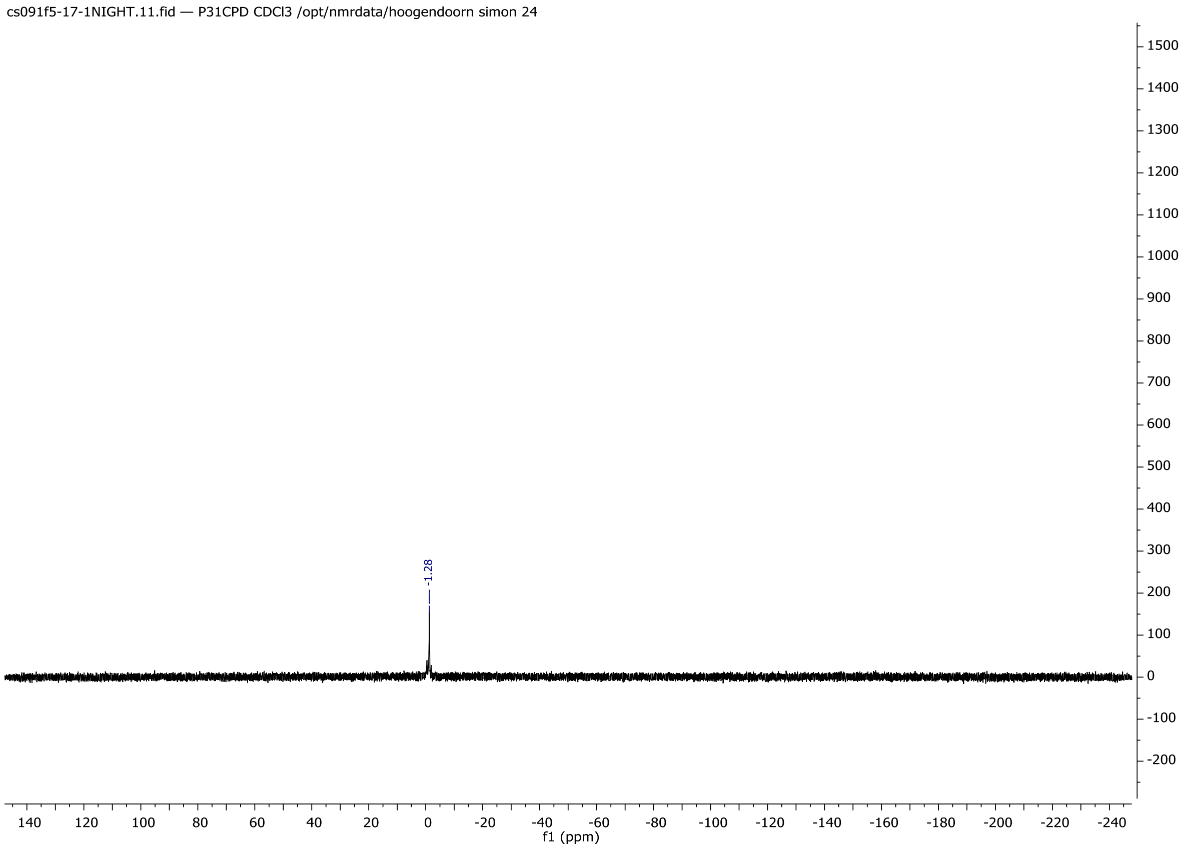
cs091f5-17-1NIGHT.11.fid — P31CPD CDCl3 /opt/nmrdata/hoogendoorn simon 24
-1.28
f1 (ppm)

# HR-MS

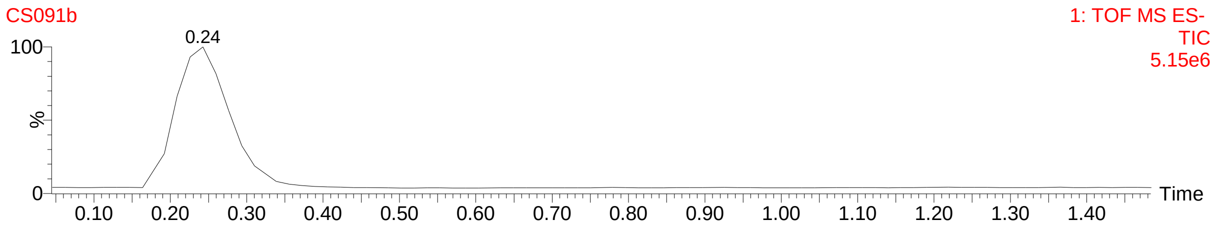
CS091b
1: TOF MS ES-
TIC
5.15e6
0.24
Time

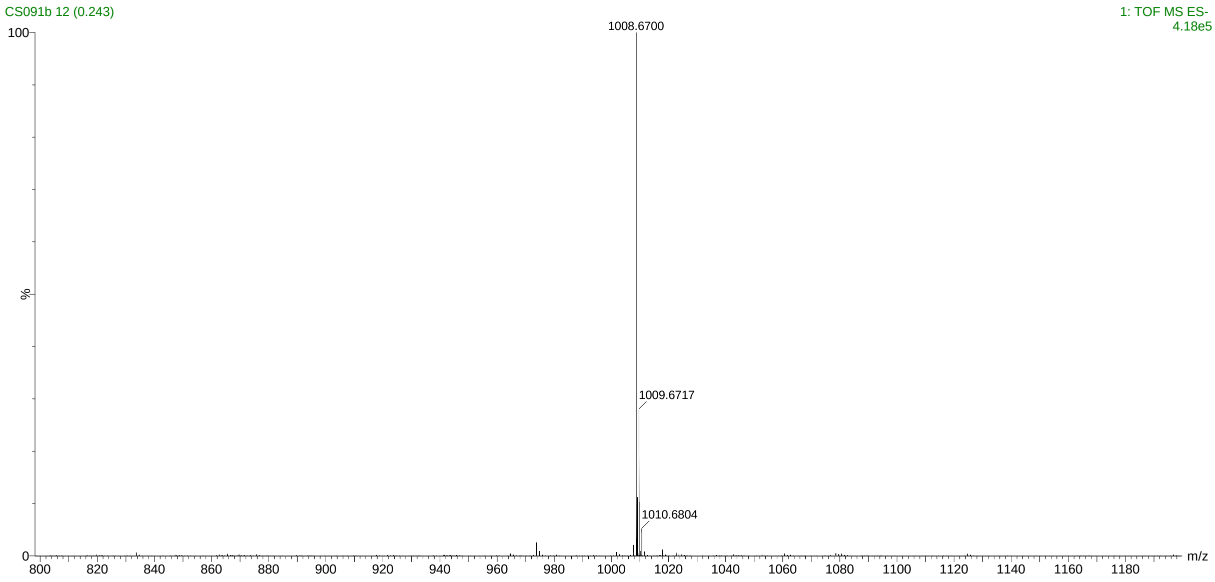
CS091b 12 (0.243)
1: TOF MS ES-
4.18e5
1008.6700
1009.6717
1010.6804
m/z

4. *tpf*ER-PE

CS104-F11-20.10.fid — MP-1Hdec13C CDCl3 /opt/nmrdata/hoogendoorn simon 4

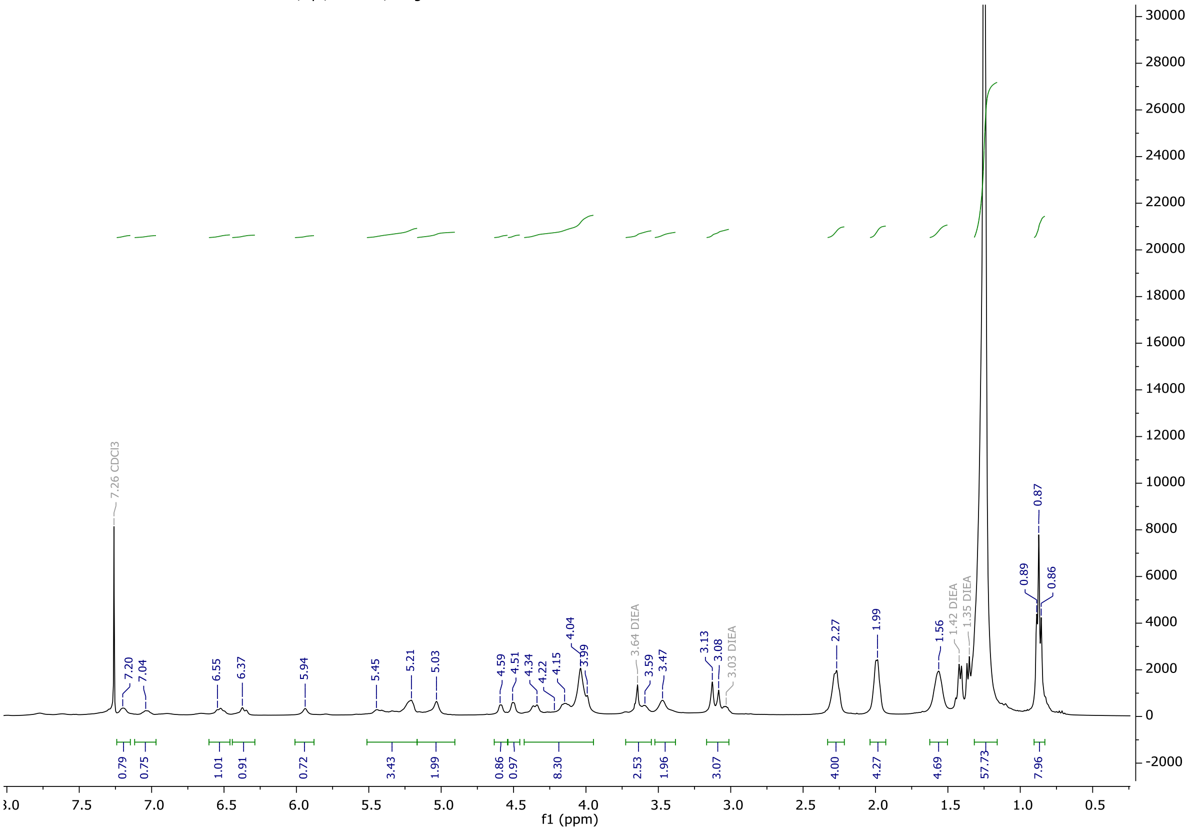

CS104-F11-20_220503.11.fid — MP_zgdc_low_amount CDCl3 /opt/topspin3.5pl7/data/hoogendoorn simon 12

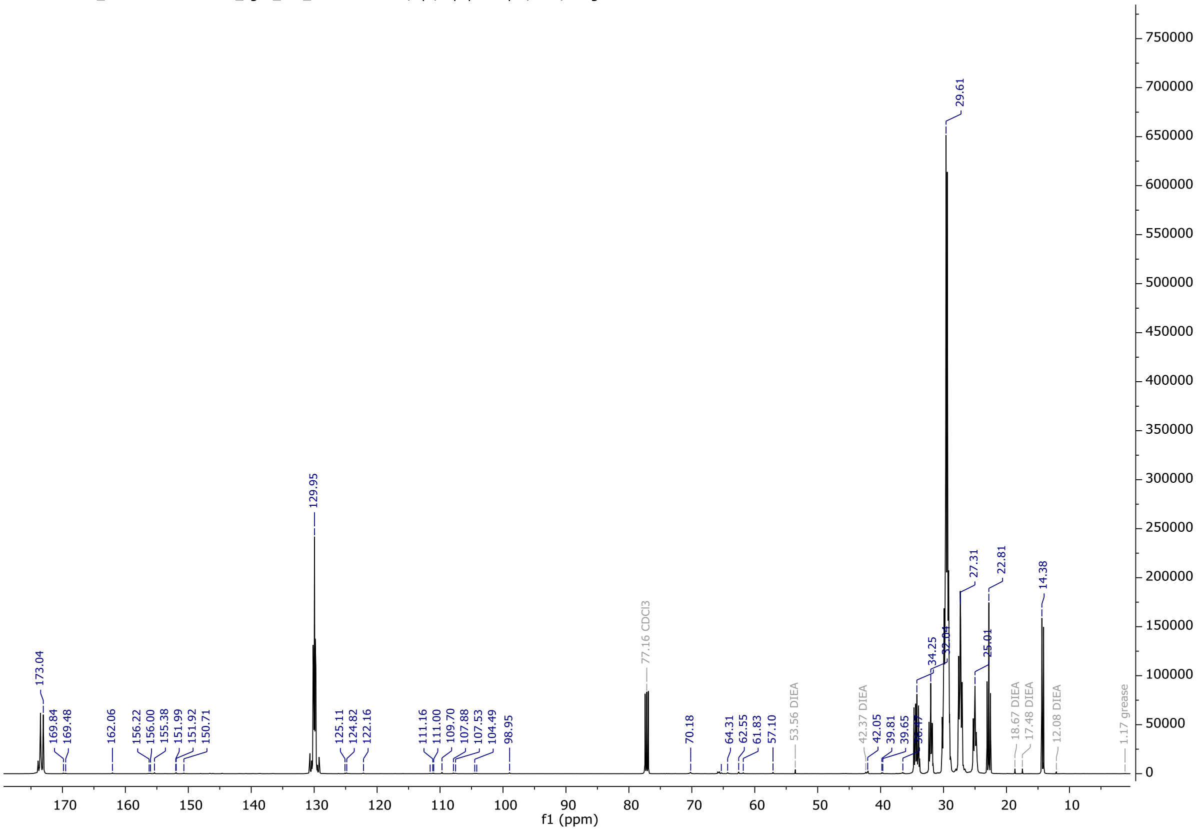

CS104-F11-20_220503.11.fid — MP_zgdc_low_amount CDCl3 /opt/topspin3.5pl7/data/hoogendoorn simon 12

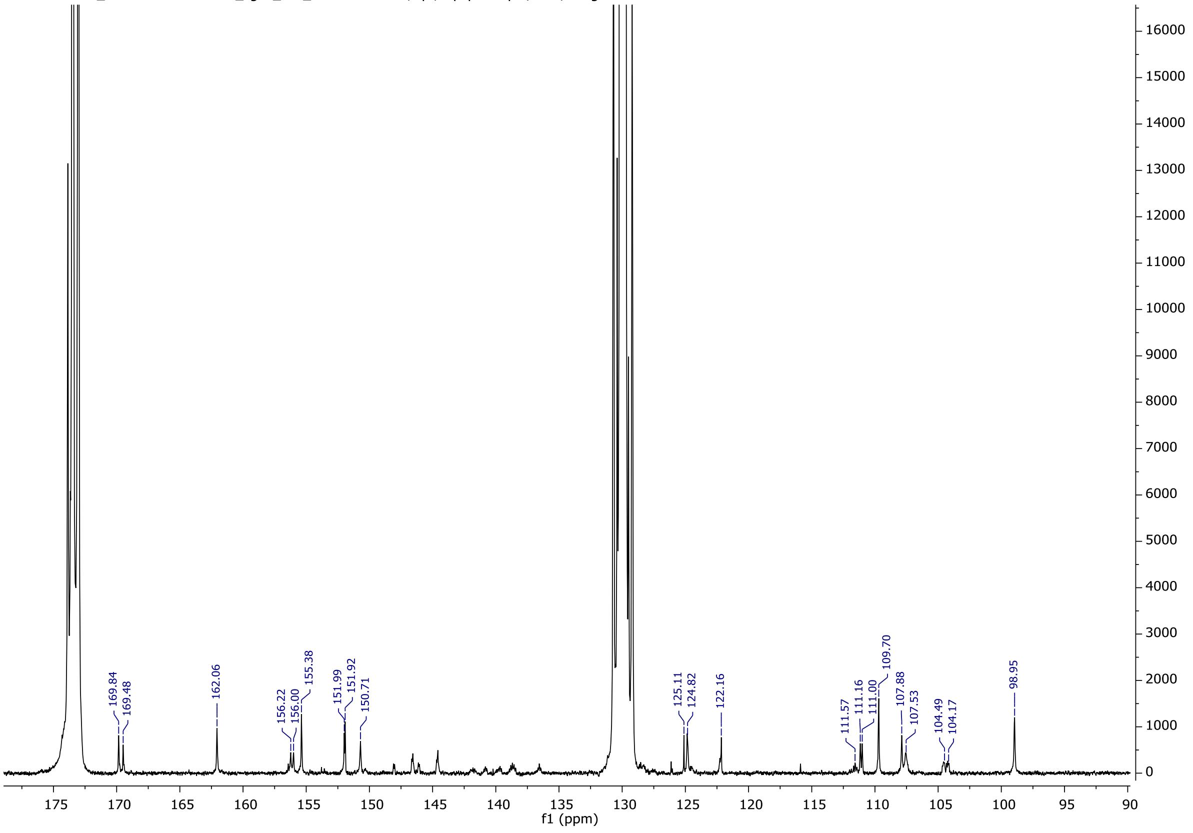

CS104-F11-20_220503.11.fid — MP_zgdc_low_amount CDCl3 /opt/topspin3.5pl7/data/hoogendoorn simon 12

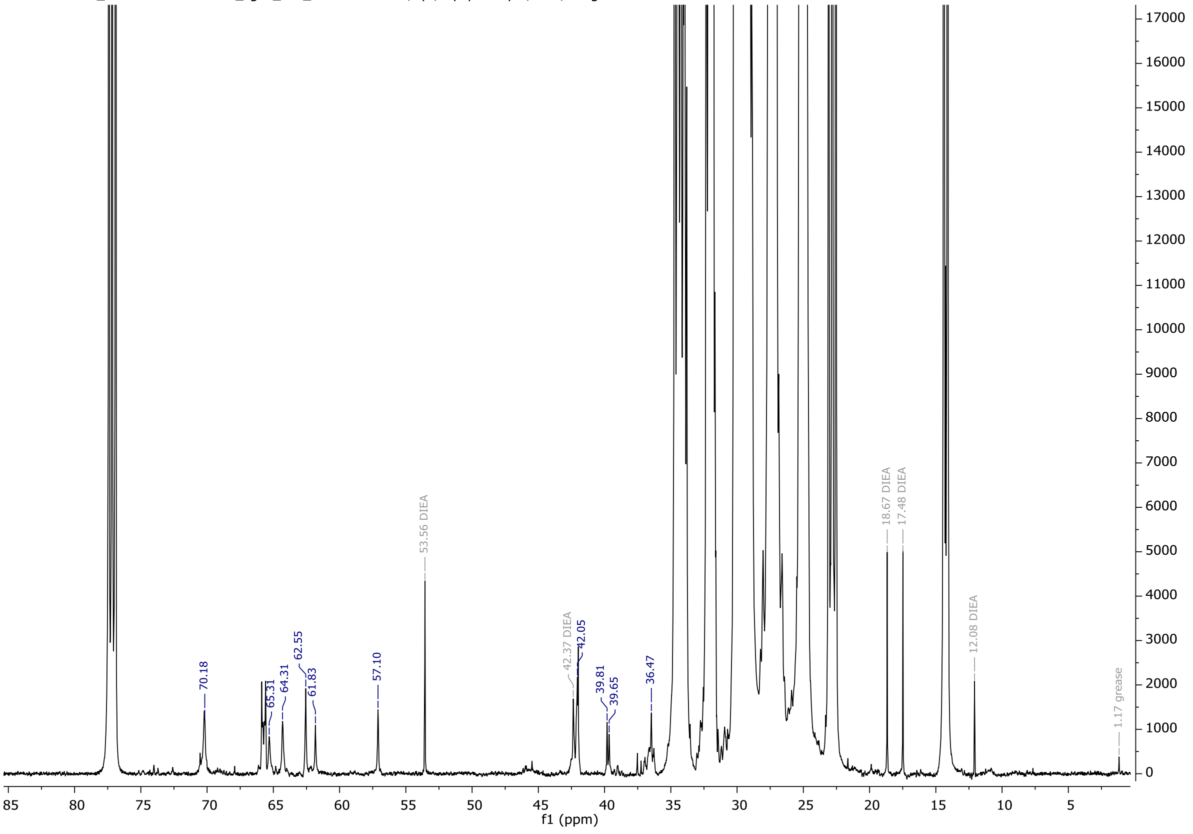

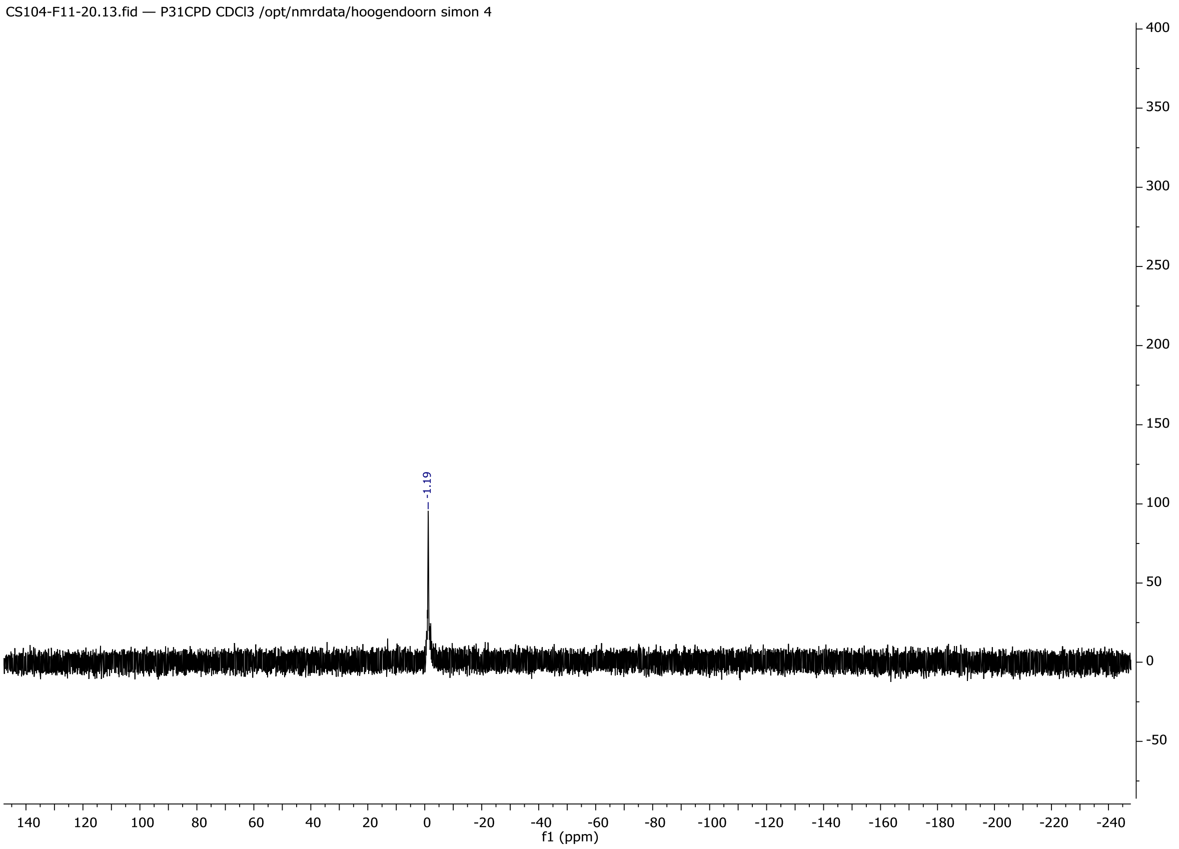
CS104-F11-20.13.fid — P31CPD CDCl3 /opt/nmrdata/hoogendoorn simon 4
-1.19
f1 (ppm)

HR-MS

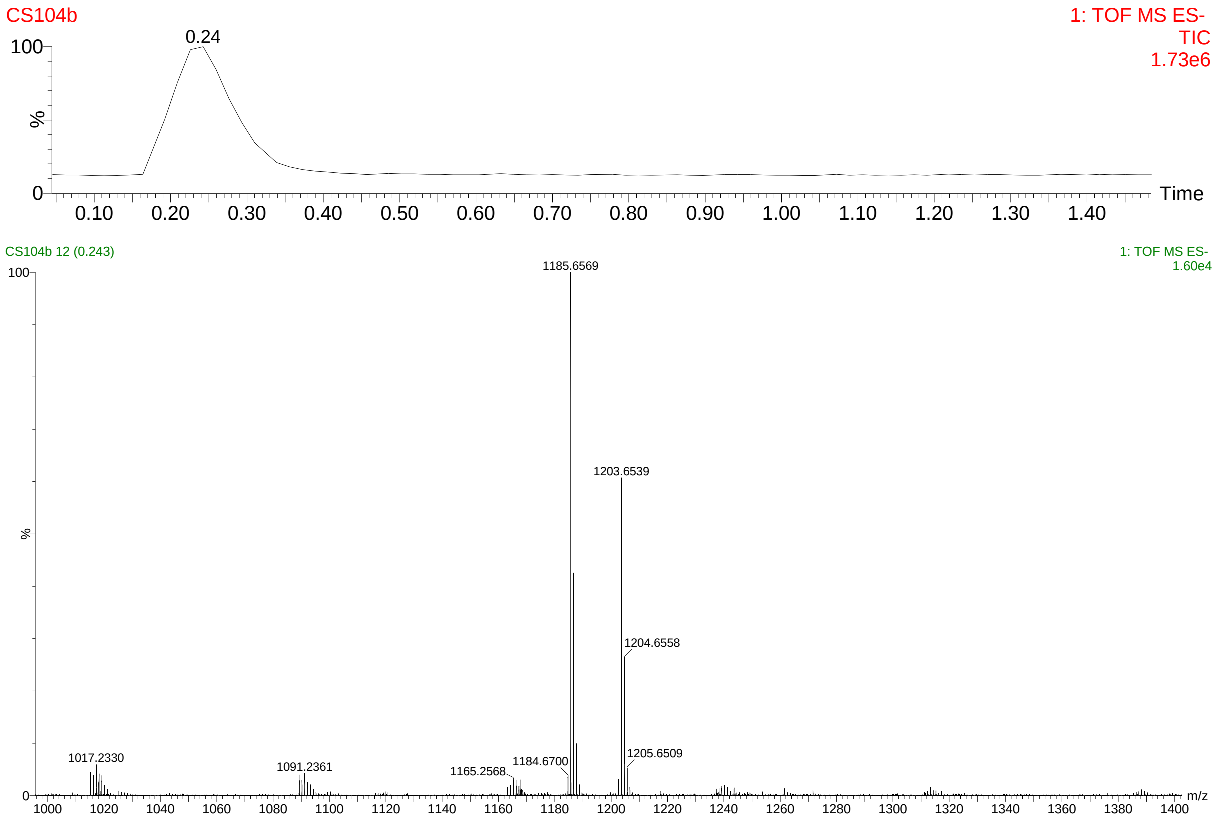
CS104b
1: TOF MS ES-
TIC
1.73e6
0.24
Time
CS104b 12 (0.243)
1: TOF MS ES-
1.60e4
1185.6569
1203.6539
1204.6558
1205.6509
1184.6700
1165.2568
1091.2361
1017.2330
m/z
